# A Hierarchical Bayesian Mixture Model for Inferring the Expression State of Genes in Transcriptomes

**DOI:** 10.1101/711630

**Authors:** Ammon Thompson, Michael R. May, Brian R. Moore, Artyom Kopp

## Abstract

Transcriptomes are key to understanding the relationship between genotype and phenotype. The ability to infer the expression state (active or inactive) of genes in the transcriptome offers unique benefits for addressing this issue. For example, qualitative changes in gene expression may underly the origin of novel phenotypes, and expression states are readily comparable between tissues and species. However, inferring the expression state of genes is a surprisingly difficult problem, owing to the complex biological and technical processes that give rise to observed transcriptomic datasets. Here, we develop a hierarchical Bayesian mixture model that describes this complex process, and allows us to infer expression state of genes from replicate transcriptomic libraries. We explore the statistical behavior of this method with analyses of simulated datasets—where we demonstrate its ability to correctly infer true (known) expression states—and empirical-benchmark datasets, where we demonstrate that the expression states inferred from RNA-seq datasets using our method are consistent with those based on independent evidence. The power of our method to correctly infer expression states is generally high and, remarkably, approaches the maximum possible power for this inference problem. We present an empirical analysis of primate-brain transcriptomes, which identifies genes that have a unique expression state in humans. Our method is implemented in the freely-available `R` package `zigzag`.

**Significance Statement:** How do the cells of an organism—each with an identical genome—give rise to tissues of incredible phenotypic diversity? Key to answering this question is the *transcriptome*: the set of genes expressed in a given tissue. We would clearly benefit from the ability to identify *qualitative* differences in expression (whether a gene is active or inactive in a given tissue/species). Inferring the expression state of genes is surprisingly difficult, owing to the complex biological processes that give rise to transcriptomes, and to the vagaries of techniques used to generate transcriptomic datasets. We develop a hierarchical Bayesian mixture model that—by describing those biological and technical processes—allows us to infer the expression state of genes from replicate transcriptomic datasets.

---

A central goal of biology is to understand the relationship between genotype and phenotype: how is it that the cells of a multicellular organism—each with an identical genome— give rise to tissues and organs of astonishing structural and functional diversity? Our current understanding of the connection between genotype and phenotype is largely based on the *transcriptome*; the set of genes that are expressed in a given tissue. Tissue-specific transcriptomes can change in two fundamental ways during development and evolution; *quantitative* changes in expression level through up- or down-regulation of genes that were already active in a given tissue, and *qualitative* changes in expression, where a gene is activated or inactivated in that tissue.

Our ability to explore the genomic basis of organismal phenotype has been greatly enhanced by the advent of RNA-sequencing (RNA-seq) techniques. However, the utility of quantitative transcriptomic approaches—*i.e.*, those based on relative differences in the expression levels of cells and tissues— is limited by both biological and technical issues. First, the relationship between the abundance of transcripts of a given gene and the corresponding abundance of the encoded protein can be obscured by post-transcriptional regulation on both physiological and evolutionary time scales (1–14). Second, the nature of RNA-seq data complicates comparison of expression levels between tissues and/or species. That is, gene-expression estimates from RNA-seq data are in relative units; the number of transcripts sampled in an RNA-seq library is not proportional to the total RNA content of a sample. Consequently, a gene with a similar number of transcripts in two different samples may have very different relative-expression levels (15).

Evaluating the qualitative expression state of genes—*i.e.*, active or inactive—in transcriptomes offers unique advantages for exploring the genotype-phenotype connection. In both development and evolution, a qualitative change in gene-expression state may be more likely to induce a qualitative change in cellular phenotype. Moreover, qualitative differences in expression state are readily comparable; *e.g.*, it is straight-forward to interpret the observation that a given gene is active in one tissue or species but inactive in another. Genes that are expressed in tissue- or cell-restricted patterns are candidates for the unique characteristics of those tissues and cells.

The potential of qualitative transcriptomic approaches is hindered by the difficulty of inferring the expression state of genes. There are three primary factors that complicate our ability to identify expression states. First, transcription is an inherently noisy process (16–18); there is compelling evidence that non-functional genes are often expressed at low levels (19, 20). Therefore, detecting transcripts of a given gene in a given tissue does not necessarily indicate that it is active. Second, we may fail to detect transcripts of a given gene owing to biological and technical factors, including its expression level, its length, and the sequencing depth of the library. Therefore, detecting zero transcripts of a given gene in a given tissue does not necessarily indicate that it is inactive. Third, even when we detect transcripts of a given gene, its measured expression level is likely to vary among libraries owing to both biological factors (*e.g.*, population-level variation) and technical factors (*i.e.*, the relative abundance of a given transcript in a given library depends on the total transcript number of that library). Therefore, the rank order in expression level of two genes in one library may differ from their rank order in a second library, which complicates methods that infer the expression state of genes based on fixed expression-level thresholds (17, 21).

Here, we present a hierarchical Bayesian model that describes the biological and technical processes that generate transcriptomic data that—by explicitly accommodating the factors described above—allows us to infer the expression state of each gene from replicate RNA-seq libraries. We present analyses of simulated datasets that validate the implementation and characterize the statistical behavior of our hierarchical Bayesian model. We also apply our method to several empirical datasets, and demonstrate that the expression states inferred using our method are consistent with expectations based on independent information, such as epigenetic marks and developmental-genetic studies. Finally, we demonstrate our method with an empirical analysis of primate-brain transcriptomes that identifies the set of genes with unique expression states in regions of the human brain.

## Inferring Gene Expression State from Transcriptomes

Here we develop a hierarchical Bayesian mixture model that describes the biological and technical processes that give rise to transcriptomic datasets with the objective of inferring the expression state of each gene. A given transcriptomic dataset is comprised of one or more replicate libraries, where each replicate library consists of the relative number of transcripts for each gene on the log scale (*e.g.*, log-TPM). Our model includes two levels: the upper level describes the distribution of the true (unobserved) expression level of each gene, and the lower level describes the variation in the observed expression levels as a consequence of biological and technical factors. To develop intuition for this model, we first describe how our inference model can be used to simulate data. We then outline the procedure for inferring the parameters of the mixture model from empirical data, and how to assess the fit of our model to empirical datasets. We provide detailed descriptions of the statistical model, inference machinery, model-comparison methods, and implementation in Sections 1.1 through 1.5 of the Supplemental Material.

### A Generative Model

To introduce our model, it is helpful to imagine using it to generate data. We begin in the upper level of the hierarchical model, which reflects the true expression level of genes in the transcriptome; this is a mixture distribution comprised of inactive (blue) and active (red) genes (Fig. 1A). For each gene, we randomly draw the expression state from this mixture distribution: specifically, a gene is inactive (active) with probability proportional to the area under the blue (red) distribution. If the selected expression state is inactive, it will either have zero transcripts (with probability proportional to the blue spike), or non-zero transcripts, in which case its expression level is drawn from the inactive (blue) normal distribution. Conversely, if the selected expression state is active, its expression level will be drawn from the active (red) normal distribution.

**Fig. 1.**
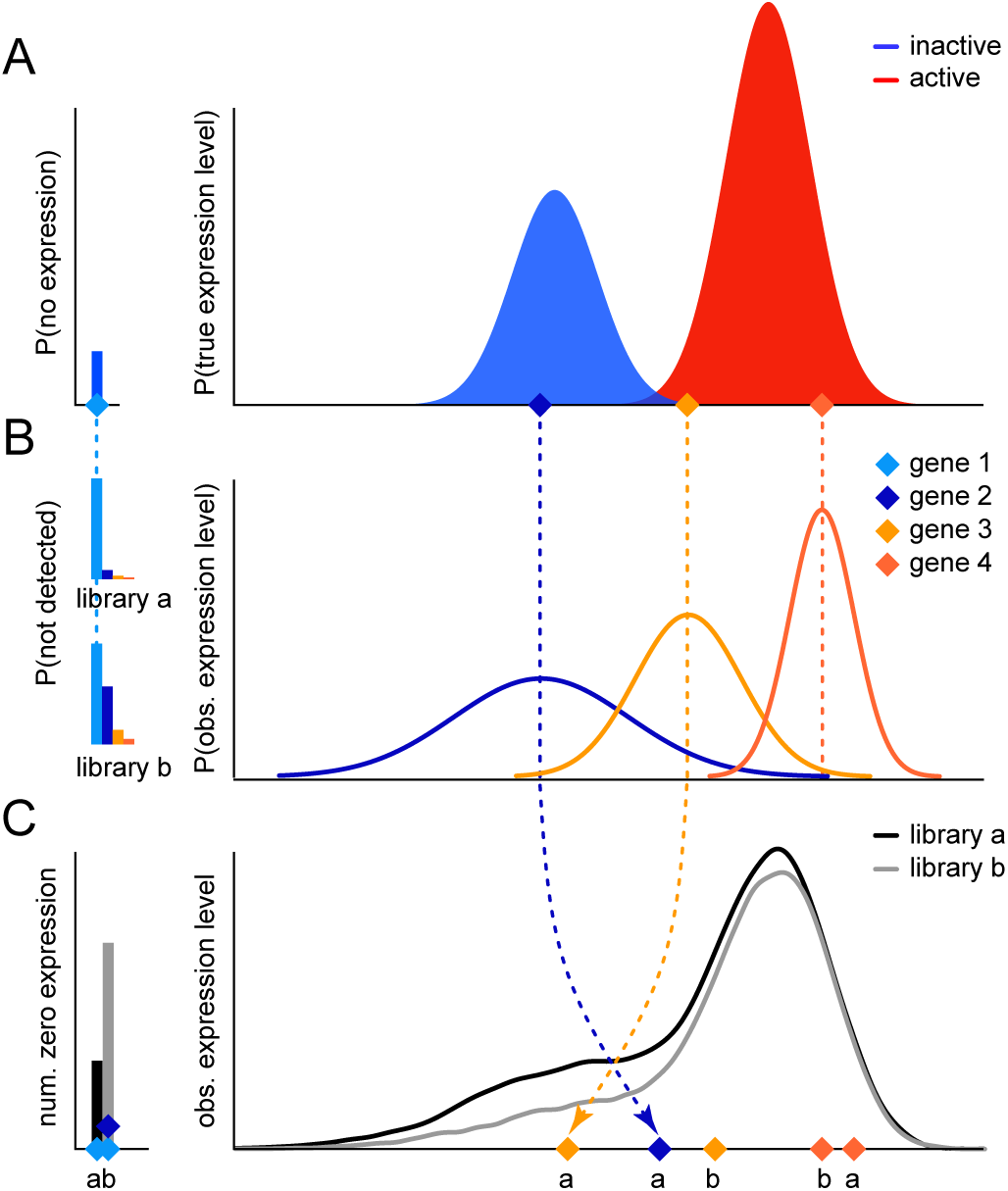
A hierarchical Bayesian mixture model for inferring the expression state (active or inactive) of genes from replicate transcriptomic libraries. We introduce our model by describing how it could be used to simulate transcriptomic libraries. Panel A depicts the true expression state—inactive(blue) or active(red)—and expression level of all genes in the transcriptome. We simulate each gene by randomly sampling from this mixture distribution. Our first four draws include two inactive genes—one with zero expression (gene 1, in the “spike” at left) and one with non-zero expression (gene 2)—and two active genes (3 and 4). Panel B depicts the probability that a gene is not detected (left) and—given detection—the observed expression level of each gene across libraries (right). For each simulated gene, we first determine whether it is detected in each library; if a gene is *not* detected in a given library, it will have an observed expression level of zero (*i.e.*, be assigned to the library-specific spikes at left). If a gene *is* detected in a given library, its observed expression level will be drawn from a normal distribution (the gene-specific distributions at right) that describes its variation across all libraries in which it is detected. These normal distributions have a mean equal to the true expression level of each gene and a gene-specific variance. Panel C depicts the observed expression level of all genes—with zero transcripts (left) or non-zero transcripts (right)—in two replicate libraries. For example, gene 1 was not detected in either library because its true expression is zero (panel A). The observed expression levels of genes 2–4 were drawn from their corresponding normal distributions (panel B), resulting in zero transcripts for gene 2 in library *b*, and non-zero transcripts for the remaining genes in both libraries. To generate a complete library with *n* genes, we repeat the above procedure *n* times. Like real datasets, transcriptomes simulated under our model have bimodal expression levels, albeit the active and inactive distributions are obscured by library-specific factors (*e.g.*, sequencing depth) and gene-specific factors (*e.g.*, gene length, true expression level, and gene-specific variance). When used as a generative model, we assume the parameter values are known and the data are unknown; conversely, when used as an inference model, we assume the data are known (observed) and the parameter values are unknown (inferred).

Having simulated the true expression level for each gene, we now simulate their observed expression levels (Fig. 1B). For each gene, we first determine whether it is detected in each transcriptomic library. The probability that a gene is detected in a given library depends on its true expression level, its length, and library-specific factors (*e.g.*, sequencing depth). For any library in which the gene is not detected, the observed expression level will be zero (Fig. 1B, left). For all libraries in which the gene is detected, we draw its observed expression level from a normal distribution, with a mean equal to its true expression level and a gene-specific variance (Fig. 1B, right).

Like empirical datasets, transcriptomic libraries simulated under our model have a characteristic bimodal distribution, with a dominant right mode and a left shoulder (Fig. 1C). We simulate a set of transcriptomic libraries by repeatedly drawing from the gene-specific distributions described above. Biological and technical sources of variation (in the lower level of our hierarchical model) largely obscure the distinct inactive and active distributions of true expression levels (in the upper level of our hierarchical model). Note that the number of genes with zero transcripts may differ among libraries owing to differences in their sequencing depth. Additionally, the rank order of the expression level of genes may vary among libraries owing to variation in the observed expression level of each gene; *e.g.*, an inactive gene may have a higher observed expression level than an active gene in a given libraries (Fig. 1C, see arrows).

### Model Parameters and Inference

Our goal is to infer the expression state of each gene from replicate (observed) transcriptomic libraries using our hierarchical Bayesian mixture model. When used as a generative model (as above), we assume that the parameter values are known and the data are unknown. To perform inference under our model, we treat the data as known (observed), and treat the parameter values as unknown. Here, we describe the parameters of the lower and upper levels of the hierarchical model, and adopt a Bayesian approach to estimate those parameters from observed transcriptomic data.

Our hierarchical Bayesian model describes the processes that give rise to our observed dataset, which we denote **X**, that is comprised of two or more replicate transcriptomic libraries. The lower level of our model describes the observed expression levels for each gene across all libraries. Specifically, we model the variation in observed expression levels for each gene across libraries with a gene-specific variance parameter. We assume that the variance parameter is inversely related to the (log) expression level, where genes of similar expression levels have similar levels of variation across replicate libraries. Additionally, the lower level includes library-specific parameters that impact the probability that a gene is detected in each library. We represent all of the parameters in the lower level of our model with the container parameter *θ*_1_.

The upper level of our hierarchical Bayesian model describes the distribution of true (unobserved) expression levels, which we denote **Y**. We assume that the true expression levels of genes can be divided into two components; those genes that are actively expressed, and those that are not actively expressed. The assignment of each gene to these in/active expression-state components is described by parameter 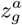, where 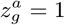 indicates that the gene is assigned to the active component, and 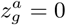 indicates that it is assigned to the inactive component. We refer to the assignments of *all* genes to the in/active-expression states as ***z***^*a*^. We further assume that inactive genes can be subdivided into two subcomponents: one with zero expression, and another with non-zero expression. Similarly, active genes may be subdivided into one or more subcomponents with distinctly different expression levels (*e.g.*, housekeeping genes may collectively have higher true expression levels relative to other genes). A given model assumes a specific number of active subcomponents, *e.g.*, the model in Figure 1A has a single active subcomponent; a model with two active subcomponents would have two red distributions. We can specify a set of distinct models with different numbers of active subcomponents, and compare their fit to a given dataset (see below). We represent all of the parameters in the upper level of our model—describing true active and inactive distributions—with the container parameter *θ*_2_.

We infer the joint posterior probability distribution of the hierarchical model parameters—including the set of parameters describing the expression state of all genes, ***z***^*a*^—given our observed transcriptomic data, **X**, by applying Bayes’ theorem:

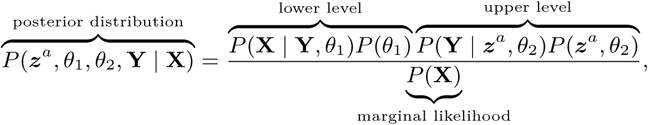

where the first term in the numerator is the joint probability of the lower level of the hierarchical model given the local model parameters, *θ*_1_, the second term is the joint probability of the upper level of the hierarchical model given the local model parameters, *θ*_2_, and the denominator is the average probability of the data under the model (the marginal likelihood).

The posterior probability distribution, *P* (***z***^*a*^, *θ*_1_, *θ*_2_, **Y** | **X**), cannot be calculated analytically because the marginal likelihood, *P* (**X**), is impossible to evaluate. Accordingly, we use a numerical algorithm—Markov chain Monte Carlo (MCMC; 22–25)—to approximate the posterior probability distribution. The MCMC algorithm samples parameter values in proportion to their posterior probability. From these MCMC samples, we compute the posterior probability that a given gene is active as the fraction of MCMC samples where 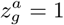. We validated our MCMC implementation by running it under the prior, and by measuring coverage probabilities using simulated data.

### Model Checking

The Bayesian approach for assessing model adequacy is called *posterior-predictive assessment* (26). This approach is based on the following premise: if our inference model provides an adequate description of the process that gave rise to our observed data, then we should be able to use that model to simulate datasets that resemble our original data. The resemblance between the observed and simulated datasets is quantified using a summary statistic. We use three summary statistics: (1) the upper-level Wasserstein statistic (which measures the discrepancy between the expected and realized *true* expression levels), (2) the lower-level Wasserstein statistic (which measures the discrepancy between the expected and realized *observed* expression levels), (3) the Rumsfeld statistic (which measures the discrepancy between the observed and expected number of undetected genes).

## Simulation Study

We explored the ability of our hierarchical Bayesian mixture model to correctly infer the expression state (active or inactive) of genes via simulation. First, we characterize the *power* to correctly identify the expression state of genes as a function of: (1) the degree of overlap between the true inactive and active distributions of expression levels, and; (2) the number of replicate transcriptomic libraries used to estimate the model parameters. We then characterize the *robustness* of expression-state estimates when the number of active subcomponents in the model is misspecified. We provide detailed descriptions of the simulation analyses and results in Sections 2.1 and 2.2 of the Supplemental Material.

### Replicate libraries improve our ability to correctly infer expression states

We expect our ability to correctly infer expression states will depend on the disparity between the true distributions of active and inactive expression levels and the number of replicate libraries. Specifically, we expect the power to increase as we: (1) decrease the degree of overlap between the true in/active distributions, and; (2) increase the number of replicate libraries used to estimate the model parameters.

We simulated data with low, moderate, and high levels of overlap between the true active and inactive distributions. For each condition, we simulated datasets comprising 2, 4, and 6 replicate libraries. For each unique combination of overlap and library number, we simulated 100 datasets. We measured power by evaluating the posterior probability of the true expression state, averaged across all of the genes in the transcriptome. Our results reveal that our method generally has good power (on average, we inferred the correct expression state for ≈ 90% of the simulated datasets), which increases with the number of replicate libraries and the disparity between true active and inactive distributions (Fig. 2, left).

**Fig. 2.**
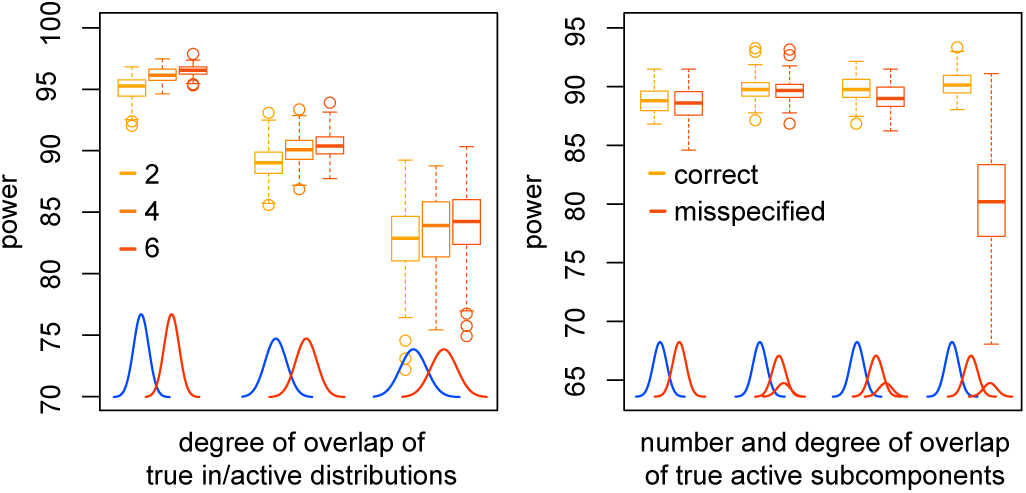
Exploring the power and robustness of our hierarchical Bayesian mixture model to correctly infer the expression state of genes from simulated data. Left panel: We explored the power of our method by simulating datasets with 2, 4, and 6 replicate libraries under varying degrees of overlap (low, moderate and high) between the true active (red) and inactive (blue) distributions. The power to infer the true expression state is generally high, and increases with the number of replicate libraries and/or the degree of separation between the true in/active distributions. Right panel: We explored the robustness of our method to model misspecification by simulating datasets with one or two active (red) subcomponents. For simulated datasets with two active subcomponents, we varied their degree of overlap (low, moderate, and high). We analyzed each simulated dataset under two models: a model with one active subcomponent, and a model with two active subcomponents. The power of the method to correctly infer expression states is robust to model overspecification: estimates from datasets with one active subcomponent are virtually identical under the correct and overspecified models (leftmost pair of boxplots). Similarly, the power of the method is robust to moderate model underspecification: estimates from datasets with two active subcomponents are virtually identical under the correct and underspecified models (two middle pairs of boxplots), except when the degree of disparity between the two active subcomponents is extreme (rightmost pair of boxplots).

### Estimates of expression state are robust to model misspecification

We expect our ability to correctly infer expression states of genes will be adversely affected when the number of assumed active subcomponents in the hierarchical mixture model differs from the true number active subcomponents, *i.e.*, when the model is misspecified. The model may either include too many active subcomponents (*overspecified*) or too few active subcomponents (*underspecified*).

We simulated datasets with one or two active subcomponents. In the latter scenario, we varied the degree of overlap (low, moderate, and high) between the active subcomponents. For each scenario, we simulated 100 datasets, each with four replicate libraries. For each simulated dataset, we inferred expression states under a model with one or two active subcomponents. When the model was overspecified (*i.e.*, with one true and two assumed active subcomponents), the expression-state estimates were virtually identical to those inferred under the correctly specified model. When the model was underspecified (*i.e.*, with two true and one assumed active subcomponents), the accuracy of expression-state estimates decreased as the disparity between the two true active subcomponents increased. These results indicate that expression-state estimates are robust to overspecificaiton and moderate underspecification, but are sensitive to severe underspecification (Fig. 2, right).

## Empirical Benchmarks

We augment our simulation study—where we assessed the ability of our method to recover true/known parameter values— with analyses of two empirical datasets where the expression states are “known” from external evidence. Specifically, we characterize the power to correctly identify the expression state of genes in human-lung transcriptomes (where expression states are predicted by epigenetic marks), and *Drosophila*-testis transcriptomes (where expression states are known from developmental-genetic studies). These special cases—where expression states have been determined by independent means— provide a rare opportunity to empirically benchmark the performance of our method. We provide detailed descriptions of the empirical analyses and results in Section 2.3 of the Supplemental Material.

### Human-lung transcriptomes

Our first empirical benchmark is a human-lung dataset comprising 427 libraries with 19,154 protein-coding genes sourced from the GTEx RNA-seq database (27, 28). We inferred the expression state of each gene using the extensive epigenomic dataset for human-lung tissues from the Roadmap Epigenomics Consortium (27, 29, 30); this dataset includes 15 epigenetic marks that are strongly associated with expression state. Using this epigenetic evidence, we were able to confidently classify the expression state of 11,968 genes (see Supplemental Material for details); we identified 7,261 active and 4,707 inactive genes (Fig. 3, left). These expression-state assignments (treated as known) provide an empirical benchmark to assess the power of our method.

**Fig. 3.**
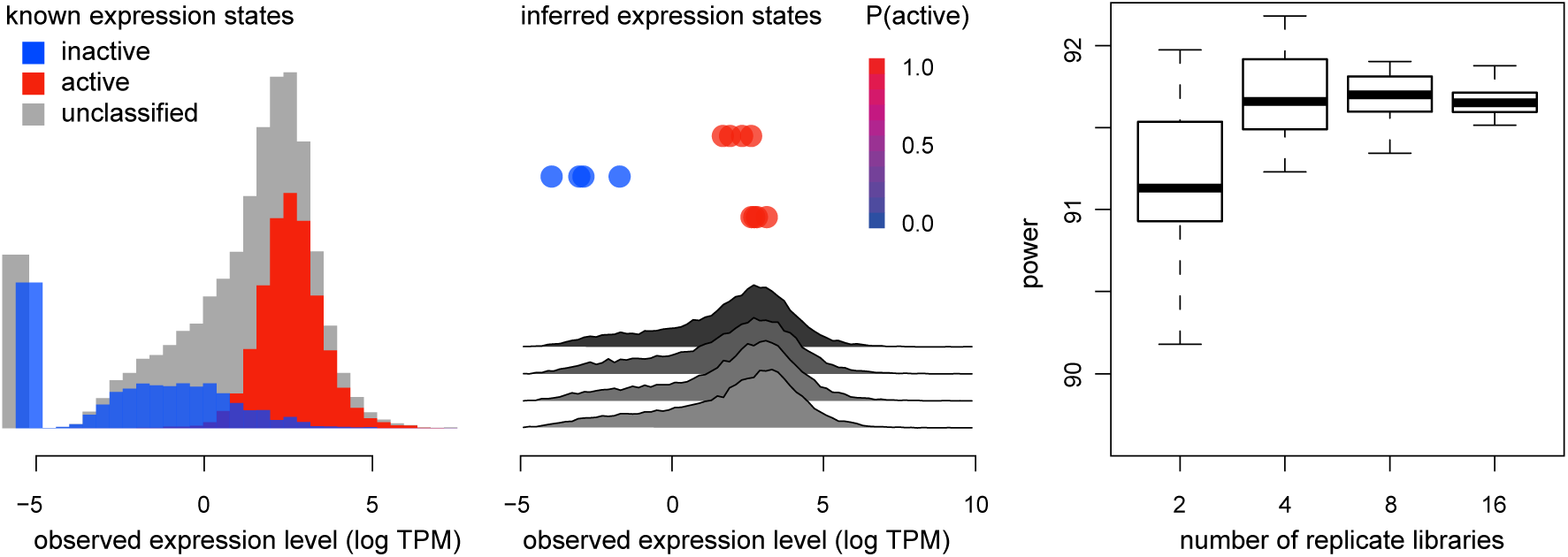
Exploring the power of our hierarchical Bayesian mixture model to accurately infer known expression states of genes in the human-lung transcriptome. Left panel: Average observed expression levels of genes in the human-lung transcriptome; active (red) and inactive (blue) genes are known from epigenetic marks, providing an empirical benchmark to assess the performance of our method (genes that could not be classified using epigenetic marks are shown in gray). Note that the expression levels of active and inactive genes fall into two distinct but overlapping distributions. Center panel: We used our model to infer the expression state of all genes from datasets consisting of 2, 4, 8 and 16 randomly selected libraries: we depict estimates for three example genes, where active (red) or inactive (blue) expression states were inferred from a dataset with four randomly selected, replicate libraries (gray distributions). Right panel: We compared our inferred expression states to the known expression states; the power of our method to correctly infer the known expression states is generally high. The use of multiple replicate libraries improves power, and this benefit is realized with only a modest number (four) of replicate libraries. Boxplots represent variation in estimates of power across the 10 sets of randomly selected datasets for each number of libraries.

Next, we used our hierarchical Bayesian mixture model to infer the expression state of all 19,154 protein-coding genes. We analyzed data subsets consisting of 2, 4, 8, and 16 randomly selected replicate libraries. For each number of libraries, we sampled 10 independent datasets (*e.g.*, 10 sets of 2 libraries, 10 sets of 4 libraries, etc.). We measured power by evaluating the posterior probability of the true expression state for each gene, averaged across all of the genes in the transcriptome. The results of these empirical analyses confirm the findings of our simulation study; the power is generally high (> 90% in all cases), and the method performs well with a modest number (four) of libraries (Fig. 3, right).

### *Drosophila*-testis transcriptomes

Our second empirical benchmark is a *Drosophila*-testis transcriptomic dataset (with four libraries) that we generated for this study. In this experiment, we assessed the power of our method to correctly identify expression states in a challenging empirical setting; *i.e.*, where genes are known to be active in a small number of cells within a tissue, with correspondingly low tissue-wide expression levels. Germline stem cells and several types of somatic cells collectively comprise the stem-cell niche at the tip of the testis (restricted to 20 to 30 cells per testis); we used developmental-genetic evidence to identify 39 active genes in the stem-cell niche (see Table S.4 for a list studies). Conversely, we identified 119 genes that are known to encode odorant and gustatory receptors that are unlikely to be active in the testis; we therefore classify these genes as inactive.

We used our method to infer the expression state of all genes in the *Drosophila*-testis dataset. We measured power by evaluating the posterior probability of the true expression state for each gene, averaged across all of the genes in the transcriptome. As previously, the power of our method is generally high, even for genes that are actively expressed in a tiny fraction of cells in the tissue (Fig. S10). Among genes that are known to be actively expressed in the stem-cell niche, the median inferred posterior probability of being in the active expression state was 0.96, with 37 of the 39 genes inferred to be active (*P* [active] > 0.5). Among olfactory- and gustatory-receptor genes, which we assume are inactive in the testis, the median inferred posterior probability of being in the active expression state was 0.005, with 111 of the 119 genes inferred to be inactive (*P* [active] < 0.5).

### Theoretical power analysis

Our analyses of simulated and empirical-benchmark datasets demonstrate that our method generally has high power to infer true/known expression states. Here, we attempt to evaluate the absolute power of our method. To this end, we first establish an upper bound on the power to infer expression states (under a method that requires *known* expression states), and then compare the power of our method to this reference.

Specifically, we imagine a threshold-based method; *i.e.*, where a gene is inferred to be active if its relative expression level exceeds a fixed threshold value. Unlike actual threshold-based methods (28, 31, 32), this “omniscient” threshold-based method knows the true expression state of each gene. Because this method is aware of the true expression states, it can choose the perfect threshold value that simultaneously maximizes the number of true active genes it infers to be active (the true-positive rate) and minimizes the number of true inactive genes it infers to be active (the false-positive rate).

We first characterize the power of the omniscient-threshold method by applying it to the empirical-benchmark datasets (where the expression state of each gene is known). Specifically, we characterize its power by plotting receiver operating characteristic (ROC) curves: for each possible threshold value, we compute the true- and false-positive rate, and plot the true-positive rate as a function of the false-positive rate (Fig. 4, orange curves). Note that a method with perfect power would exhibit an L-shaped ROC curve, as it would simultaneously achieve a 100% true-positive rate and a 0% false-positive rate. Conveniently, we can compare the power of two methods by comparing their ROC curves.

**Fig. 4.**
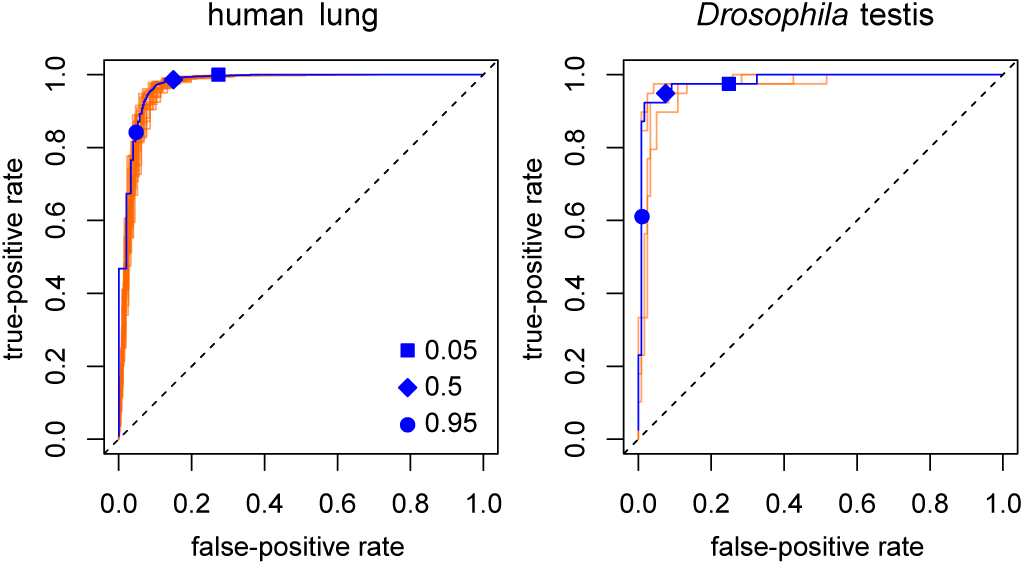
The power of our method to infer expression states approaches the practical limit for this inference problem. We used our method to infer expression states of all genes in the two empirical-benchmark datasets, human-lung (left) and *Drosophila*-testis (right) transcriptomes. A gene is assigned to the active expression state if its posterior probability of being active is greater than *P*. For all possible values of *P*, we plot the true-positive rate (the fraction of active genes correctly assigned to the active-expression state) against the false-positive rate (the fraction of inactive genes incorrectly assigned to the active-expression state) (blue curves). The resulting ROC curves characterize the discriminatory power of a method, *i.e.*, its ability to distinguish between active and inactive genes. The power of our method (which infers the *unknown* expression states) is equivalent to that of an “omniscient” threshold-based method that requires knowledge of the true expression states (orange curves, one for each library). Symbols indicate conventional *P* thresholds.

Next, we plot ROC curves for our method based on the same empirical-benchmark datasets. Our method infers the posterior probability that each gene is in/active. In principle, we could adopt any posterior-probability threshold to classify the expression state of each gene. Accordingly, we plot ROC curves by computing the true- and false-positive rate for all posterior-probability thresholds between 0 and 1 (Fig. 4, blue curves).

Remarkably, the power of our method is virtually identical to that of the omniscient threshold-based method for both the human-lung and the *Drosophila*-testis datasets (Fig. 4). These results demonstrate that our method—under the typical inference scenario, where the true expression states of genes are unknown—is able to correctly infer expression states as well as a method that requires *a priori* knowledge of the true expression states.

## Empirical Application

Our analyses of simulated and empirical-benchmark datasets demonstrate the ability of our hierarchical Bayesian mixture model to reliably infer the expression state of genes in transcriptomic libraries. Here, we provide an empirical demonstration of our method with analyses of primate-brain transcriptomes. Because the true expression state of these genes is not known from external evidence, this represents a more typical inference scenario.

We used our method to analyze a published primate-brain transcriptomic dataset (33). We inferred the expression state of all protein-coding genes in six brain regions—amygdala (AMY), ventral frontal cortex (VFC), dorsal frontal cortex (DFC), superior temporal cortex (STC), striatum (STR), and the area 1 visual cortex (V1C)—sampled from macaques, chimpanzees and humans. We then identified the subset of approximately 12,000 1:1:1 orthologous genes in the three species. From these, we identified the subset of genes with a unique expression state in humans, *i.e.*, where a given gene is inferred to be in/active in humans but not chimpanzees and macaques. Across the six brain regions, we identified 9 to 20 genes that were uniquely active in humans and 16 to 23 genes that were uniquely inactive in humans, with the greatest number of unique expression states located in the striatum (Fig. 5A).

**Fig. 5.**
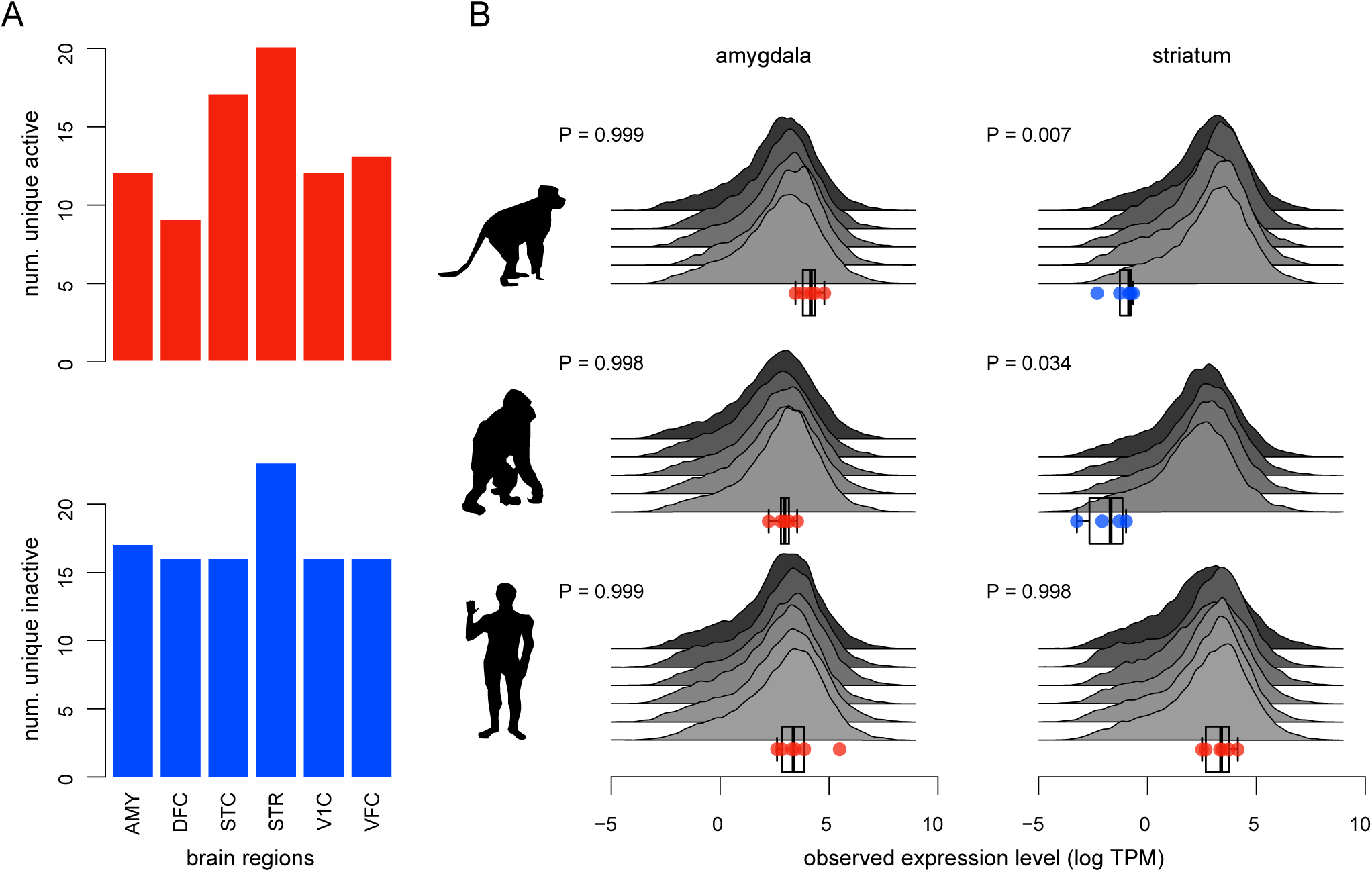
Identifying genes with unique expression states in human-brain transcriptomes. We used our method to infer the expression state of 12,000 1:1:1 orthologous genes in the transcriptomes of six brain regions of macaques, chimpanzees, and humans. We then identified the subset of these genes with unique expression states in humans. (A) Across the six brain regions, we identified between between 9 to 20 genes that were uniquely active in humans (red histogram) and 16 and 23 genes that were uniquely inactive in humans (blue histogram). (B) Here, we depict the expression state of the *Slc17a6* gene in two brain regions, amygdala and striatum, for the three species inferred from replicate transcriptomic libraries (gray distributions); active and inactive expression states are indicated with red and blue dots, respectively. In the amygdala, *Slc17a6* is active in all three species; in the striatum, *Slc17a6* is uniquely active in humans.

Genes that are uniquely active in the human brain represent factors that may be involved in human cognitive evolution. For example, we inferred that the *Slc17a6* gene is actively expressed in the human striatum but is inactive in the striatum of macaques and chimpanzees (Fig. 5B). This gene is also believed to be inactive in the mouse striatum (34), suggesting that the activation of *Slc17a6* occurred in the human lineage. This gene encodes the protein VGLUT2, which is involved in loading glutamate—a major excitatory neurotransmitter—into synaptic vesicles (34–36). These results raise the intriguing possibility that the evolutionary gain of this glutamate transporter in the human striatum may underly changes in the function of this brain region, either through the gain of a novel cell type or a change in the activity of an ancestral cell type.

## Discussion

Inferring the expression state (active or inactive) of a given gene from transcriptomic datasets is surprisingly difficult, owing both to the complexity of the underlying biological processes that give rise to transcriptomes, as well as the vagaries of the techniques that we use to generate RNA-seq libraries. Inferring the expression state of a gene based on its presence/absence in a library is unreliable: non-functional genes are often expressed at low levels, while we may fail to detect a functional gene in a given library for technical reasons. Moreover, variation in the relative expression level of a given gene among libraries will cause its rank order to vary among libraries. As a result, inferring the expression state of a given gene based on its relative-expression level in single libraries is unreliable: transcriptional noise may cause a non-functional gene to have a higher observed expression level than some functional genes that are expressed at low levels. Such considerations complicate our ability to infer expression states, especially from single libraries.

In this paper, we have developed a hierarchical mixture model that captures both important biological features— including the characteristic bimodal distribution of expression levels reflecting active expression of functional genes and background expression of nonfunctional genes (16–18, 20, 37–45)— and relevant technical factors—including differences in the detection probability of individual transcripts among replicate libraries owing to differences in their sequencing depth—that give rise to observed replicate transcriptomic libraries. We implemented our model in a Bayesian inference framework, which confers numerous benefits, including the ability to gauge uncertainty in expression-state estimates, the ability to choose among alternative models, and the ability to assess the fit of a given model to an empirical dataset. We have implemented all of the methods described in this paper in the `R` package, `zigzag`.

Encouragingly, our analyses of simulated and empirical-benchmark datasets demonstrate that our method has generally high power to recover true/known expression states, and this power increases with the number of replicate libraries. In fact, the power of our method approaches the upper bound for this inference problem (Fig. 4). Additionally, our simulations demonstrate that our method is relatively robust to model misspecification (*i.e.*, the assumed number of active subcomponents). Interestingly, our use of posterior-predictive checking indicates that our model adequately describes the processes that gave rise to all of the empirical datasets evaluated in our study (Figs. S6, S8, S11). These findings provide an empirical validation of the biological and technical features that we chose to incorporate in our model.

Our method provides a powerful means to infer expression states; this ability will play a direct role in answering many questions about the processes that give rise to transcriptomes. For example, our analyses of human-lung transcriptomes reveal that, although ≈ 98% of protein-coding genes are transcribed at detectable levels, only 67% of those genes are actively expressed in this tissue. Our method can also play a less direct—but key—role in transcriptomic/developmental-genetic pipelines, where identifying the expression states is integral to a given inference problem. For example, many developmental studies focus on actively expressed genes; our method provides a more principled alternative to conventional pre-filtering steps in quantitative RNA-seq analyses. Additionally, because expression states are inherently comparable, they can be used to address questions that involve comparisons between tissues within species. For example, we can investigate how the expression state of a gene (or set of genes) varies among a set of tissues at a given point in development. Expression states inferred using our method can also be compared across species. For example, our analysis of the primate-brain transcriptome allowed us to identify genes with unique expression states in humans, providing a narrow list of candidates that may be associated with novel brain phenotypes in humans.

The ability to identify the expression state of genes across species also lays the foundation for formal phylotranscriptomic models that describe how changes in expression state (activation and deactivation) have shaped transcriptomic diversity. Such models could be used to explore many fundamental questions, including: (1) For a given gene, what are the lineages in which it has been de/activated? (2) For a given lineage, which genes have been de/activated? (3) For the entire transcriptome, what are the relative rates of regulatory changes (activation and deactivation) and structural changes (*e.g., de novo* origination, duplication, and loss of genes)?

For many purposes, qualitative comparisons of gene-expression states between tissues and species will provide a useful complement to quantitative measures of expression level. Although it remains an open question whether changes in expression state play a particularly prominent role in phenotypic evolution, we emphasize that it is impossible to address this question without an objective method for identifying expression states. We are optimistic that—by providing a reliable and powerful means to infer the expression state of genes—our method will greatly enhance our ability to understand transcriptome evolution, and thereby illuminate the relationship between genotype and phenotype.

## Materials and Methods

We provide details of the methods and analyses in the Supplementary Material.

## ACKNOWLEDGMENTS

The authors thank Li Zhao and David Begun for technical advice and comments on the manuscript. We also thank Nerisa Riedl and Olga Barmina for technical assistance. This work was supported by NIH grant 5F32GM125107-02 to AT, NIH grant R35GM122592 to AK, and NSF grants DEB-0842181, DEB-0919529, DBI-1356737, and DEB-1457835 awarded to BRM. The authors thank Laura Crothers and Emily Delaney for comments on the manuscript.

## References

1. Geiger T, Cox J, Mann M (2010) Proteomic Changes Resulting from Gene Copy Number Variations in Cancer Cells. PLOS Genetics 6(9):e1001090.

2. Laurent JM, et al. (2010) Protein abundances are more conserved than mRNA abundances across diverse taxa. PROTEOMICS 10(23):4209–4212.

3. Schwanhäusser B, et al. (2011) Global quantification of mammalian gene expression control. Nature 473(7347):337–342.

4. Stingele S, et al. (2012) Global analysis of genome, transcriptome and proteome reveals the response to aneuploidy in human cells. Molecular Systems Biology 8(1):608.

5. Vogel C, Marcotte EM (2012) Insights into the regulation of protein abundance from proteomic and transcriptomic analyses. Nature Reviews Genetics 13(4):227–232.

6. Khan Z, et al. (2013) Primate Transcript and Protein Expression Levels Evolve Under Compensatory Selection Pressures. Science 342(6162):1100–1104.

7. Wu L, et al. (2013) Variation and genetic control of protein abundance in humans. Nature 499(7456):79–82.

8. Dephoure N, et al. (2014) Quantitative proteomic analysis reveals posttranslational responses to aneuploidy in yeast. eLife 3:e03023.

9. Artieri CG, Fraser HB (2014) Evolution at two levels of gene expression in yeast. Genome Research 24(3):411–421.

10. Battle A, et al. (2014) Impact of regulatory variation from RNA to protein. Science p. 1260793.

11. McManus CJ, May GE, Spealman P, Shteyman A (2014) Ribosome profiling reveals post-transcriptional buffering of divergent gene expression in yeast. Genome Research 24(3):422–430.

12. Liu Y, Beyer A, Aebersold R (2016) On the Dependency of Cellular Protein Levels on mRNA Abundance. Cell 165(3):535–550.

13. Ishikawa K, Makanae K, Iwasaki S, Ingolia NT, Moriya H (2017) Post-Translational Dosage Compensation Buffers Genetic Perturbations to Stoichiometry of Protein Complexes. PLOS Genetics 13(1):e1006554.

14. Wang SH, Hsiao CJ, Khan Z, Pritchard JK (2018) Post-translational buffering leads to convergent protein expression levels between primates. Genome Biology 19(1):83.

15. Lovén J, et al. (2012) Revisiting Global Gene Expression Analysis. Cell 151(3):476–482.

16. Struhl K (2007) Transcriptional noise and the fidelity of initiation by RNA polymerase II. Nature Structural & Molecular Biology 14(2):103–105.

17. Ramsköld D, Wang ET, Burge CB, Sandberg R (2009) An Abundance of Ubiquitously Expressed Genes Revealed by Tissue Transcriptome Sequence Data. PLOS Computational Biology 5(12):e1000598.

18. van Bakel H, Nislow C, Blencowe BJ, Hughes TR (2010) Most “Dark Matter” Transcripts Are Associated With Known Genes. PLOS Biology 8(5):e1000371.

19. Djebali S, et al. (2012) Landscape of transcription in human cells. Nature 489(7414):101–108.

20. Jensen TH, Jacquier A, Libri D (2013) Dealing with Pervasive Transcription. Molecular Cell 52(4):473–484.

21. Kin K, Nnamani MC, Lynch VJ, Michaelides E, Wagner GP (2015) Cell-type Phylogenetics and the Origin of Endometrial Stromal Cells. Cell Reports 10(8):1398–1409.

22. Metropolis N, Rosenbluth AW, Rosenbluth MN, Teller AH, Teller E (1953) Equation of State Calculations by Fast Computing Machines. The Journal of Chemical Physics 21(6):1087–1092.

23. Hastings WK (1970) Monte Carlo sampling methods using Markov chains and their applications. Biometrika 57(1):97–109.

24. Geman S, Geman D (1987) Stochastic Relaxation, Gibbs Distributions, and the Bayesian Restoration of Images in Readings in Computer Vision, eds. Fischler MA, Firschein O. (Morgan Kaufmann, San Francisco (CA)), pp. 564–584.

25. Green PJ (1995) Reversible jump Markov chain Monte Carlo computation and Bayesian model determination. Biometrika 82(4):711–732.

26. Gelman A, Meng XL, Stern H (1996) Posterior predictive assessment of model fitness via realized discrepancies. Statistica Sinica 6(4):733–760.

27. Consortium TG (2015) The Genotype-Tissue Expression (GTEx) pilot analysis: Multitissue gene regulation in humans. Science 348(6235):648–660.

28. Melé M, et al. (2015) The human transcriptome across tissues and individuals. Science 348(6235):660–665.

29. Ernst J, et al. (2011) Mapping and analysis of chromatin state dynamics in nine human cell types. Nature 473(7345):43–49.

30. Roadmap Epigenomics Consortium, et al. (2015) Integrative analysis of 111 reference human epigenomes. Nature 518(7539):317–330.

31. Mortazavi A, Williams BA, McCue K, Schaeffer L, Wold B (2008) Mapping and quantifying mammalian transcriptomes by RNA-Seq. Nature Methods 5(7):621–628.

32. Marguerat S, et al. (2012) Quantitative Analysis of Fission Yeast Transcriptomes and Proteomes in Proliferating and Quiescent Cells. Cell 151(3):671–683.

33. Sousa AMM, et al. (2017) Molecular and cellular reorganization of neural circuits in the human lineage. Science 358(6366):1027–1032.

34. Wallén-Mackenzie Å, Wootz H, Englund H (2010) Genetic inactivation of the vesicular glutamate transporter 2 (VGLUT2) in the mouse: What have we learnt about functional glutamatergic neurotransmission? Upsala Journal of Medical Sciences 115(1):11–20.

35. Reimer RJ, Edwards RH (2004) Organic anion transport is the primary function of the SLC17/type I phosphate transporter family. Pflugers Archiv European Journal of Physiology 447(5):629–635.

36. Wallace ML, et al. (2017) Genetically Distinct Parallel Pathways in the Entopeduncular Nucleus for Limbic and Sensorimotor Output of the Basal Ganglia. Neuron 94(1):138–152.e5.

37. Hebenstreit D, et al. (2011) RNA sequencing reveals two major classes of gene expression levels in metazoan cells. Molecular Systems Biology 7(1):497.

38. Hart T, Komori H, LaMere S, Podshivalova K, Salomon DR (2013) Finding the active genes in deep RNA-seq gene expression studies. BMC Genomics 14(1):778.

39. Piccolo SR, Withers MR, Francis OE, Bild AH, Johnson WE (2013) Multiplatform single-sample estimates of transcriptional activation. Proceedings of the National Academy of Sciences of the United States of America 110(44):17778–17783.

40. Singh A, Vargas CA, Karmakar R (2013) Stochastic analysis and inference of a two-state genetic promoter model in 2013 American Control Conference. pp. 4563–4568.

41. Rands CM, Meader S, Ponting CP, Lunter G (2014) 8.2% of the Human Genome Is Constrained: Variation in Rates of Turnover across Functional Element Classes in the Human Lineage. PLOS Genetics 10(7):e1004525.

42. Huang L, Yuan Z, Liu P, Zhou T (2015) Effects of promoter leakage on dynamics of gene expression. BMC systems biology 9:16.

43. Tiberi S, Walsh M, Cavallaro M, Hebenstreit D, Finkenstädt B (2018) Bayesian inference on stochastic gene transcription from flow cytometry data. Bioinformatics 34(17):i647–i655.

44. Wu Z, Zhang Y, Stitzel ML, Wu H (2018) Two-phase differential expression analysis for single cell RNA-seq. Bioinformatics 34(19):3340–3348.

45. Lloyd JP, Tsai ZTY, Sowers RP, Panchy NL, Shiu SH (2018) A Model-Based Approach for Identifying Functional Intergenic Transcribed Regions and Noncoding RNAs. Molecular Biology and Evolution 35(6):1422–1436.

