## Supplemental Text for "A Hierarchical Bayesian Mixture Model for Inferring the Expression State of Genes in Transcriptomes"

#### Contents

**S1 Model Description** **S2**
S1.1 Observed Transcriptomic Datasets . . . . . S2
S1.2 Bayesian Inference . . . . . S2
S1.3 Markov Chain Monte Carlo . . . . . S11
S1.4 Implementation and Validation . . . . . S17
S1.5 Assessing Model Adequacy Using Posterior Predictive Simulation . . . . . S21

**S2 Analyses** **S25**
S2.1 General Analysis Protocol . . . . . S25
S2.2 Simulation Studies . . . . . S26
S2.3 Empirical Studies . . . . . S28

### S1 Model Description

#### S1.1 Observed Transcriptomic Datasets

Our observed dataset, $\mathbf{X}$, is comprised of expression levels measured as log-transformed counts of transcripts (*i.e.*, log TPM) from $G$ genes in $R$ replicate libraries. We organize our transcriptomic dataset as a $G \times R$ matrix, where $\mathbf{X}_{g\cdot}$ represents the log-expression levels of gene $g$ across all libraries, $\mathbf{X}_{\cdot r}$ represents the log-expression levels of all genes in library $r$, and $X_{gr}$ is the log-expression level of gene $g$ in library $r$. The expression levels within RNA-seq libraries—for both protein-coding and non-coding genes—have a characteristic bimodal distribution. The components of this bimodal distribution appear symmetric on the log-scale. The left (low expression level) component consists of a single symmetric distribution that is typically centered on less than 1 TPM [1]. The bulk of the right (high expression level) component also consists of a single symmetric distribution; however, it may contain additional components comprising a smaller number of highly-expressed outlier genes, *e.g.* housekeeping genes. The expression level of individual genes varies across replicate libraries: this variation is approximately normally distributed. Genes with a high expression levels are more likely to be detected in a given library, whereas genes with lower expression levels are more likely to be undetected in a given library (*i.e.*, with zero transcripts).

#### S1.2 Bayesian Inference

The measured log-expression level of a given gene, $\mathbf{X}_{g\cdot}$, varies among libraries. To account for this variation, we imagine that these measured expression levels are manifestations of a true underlying (unobserved) log-expression level, $\mathbf{Y}$. The variation in the observed expression levels among libraries reflects both variation within natural populations, as well as the vagaries of the technical procedure that generated the libraries. In addition to adding "noise" to the measured expression levels, the technical procedure may completely fail to detect certain transcripts within a library. We assume that the true expression level of a given gene, $Y_g$, reflects either nonfunctional background expression (*e.g.*, promoter leakage), or active transcription, associated with some functional role in the cell or tissue.

Our goal is to infer the expression state ("Active/ON" or "Inactive/OFF") of each gene from the observed data, $\mathbf{X}$, under a hierarchical Bayesian model. On the upper level of the hierarchical model, we describe the distribution of the true expression level of each gene, $\mathrm{Y}_g$; on the lower level, we describe the variation in the observed expression levels, $\mathbf{X}_g$, as a consequence of biological and technical factors. The focal parameter of our model, $\boldsymbol{z}^a$, indicates the expression state of all genes; when $z_g^a = 1$, the expression state of gene $g$ is active, when $z_g^a = 0$, the expression state is inactive. We infer the joint posterior probability distribution of the hierarchical model parameters—including $\boldsymbol{z}^a$—given the observed expression-level data, $\mathbf{X}$, by applying Bayes' theorem:

$$\overbrace{P(\boldsymbol{z}^a, \theta, \mathbf{Y} \mid \mathbf{X})}^{\substack{\text{joint posterior} \\ \text{probability distribution}}} = \frac{\overbrace{P(\mathbf{X}, \mathbf{Y}, \boldsymbol{z}^a, \theta)}^{\substack{\text{joint probability} \\ \text{of the data and} \\ \text{model parameters}}}}{\underbrace{P(\mathbf{X})}_{\substack{\text{marginal probability} \\ \text{of the data}}}},$$

which is equal to the joint probability of the data and parameters, divided by the marginal probability of the data under the model (*i.e.*, the marginal likelihood). Here, $\theta$ contains the remaining model parameters, described below.

We factorize the joint probability distribution of the data and model parameters (the numerator on the right-hand side of Bayes' theorem) into two hierarchical levels—with container parameters $\theta_1$ and $\theta_2$, respectively—resulting in the joint posterior distribution:

$$P(z^a, \theta, \mathbf{Y} \mid \mathbf{X}) = \frac{\overbrace{P(\mathbf{X} \mid \mathbf{Y}, \theta_1)P(\theta_1)}^{\text{lower level}}\overbrace{P(\mathbf{Y} \mid z^a, \theta_2)P(z^a, \theta_2)}^{\text{upper level}}}{P(\mathbf{X})}. \tag{S1}$$

We describe each of the hierarchical levels of our model in detail below.

##### S1.2.1 Lower Level

The lower level of our hierarchical model describes the probability of the observed expression levels for all genes across all libraries, $\mathbf{X}$, assuming that the true expression levels of all genes, $\mathbf{Y}$, and the local model parameters, $\theta_1$, are known: $P(\mathbf{X} \mid \mathbf{Y}, \theta_1)$. We describe the data variables in Table S1. We imagine that the observed expression level for a given gene across libraries, $\mathbf{X}_{g\cdot}$, results from two sampling processes: the first process determines whether a given transcript is detected in a given library; the second process determines the variation in the observed expression levels across the libraries in which the transcript is detected.

We represent the probability of $\mathbf{X}_{g\cdot}$ as $P(\mathbf{X}_{g\cdot} \mid \mathrm{Y}_g, \sigma_g, \boldsymbol{\alpha})$, where $\boldsymbol{\alpha}$ represents factors that impact the probability that the transcript is detected in a given library, and $\sigma_g$ describes how the expression level of a given transcript varies across libraries (when the transcript is detected) (see Fig. 1B, main text). We describe the details of these parameters below and in Tables S1–S3. We assume that the observed expression levels of all genes across all libraries, $\mathbf{X}$, are independent—given the true expression levels, $\mathbf{Y}$, and the local model parameters, $\theta_1$—so that:

$$P(\mathbf{X} \mid \mathbf{Y}, \boldsymbol{\sigma}, \boldsymbol{\alpha}) = \prod_{g=1}^{G} P(\mathbf{X}_{g\cdot} \mid \mathrm{Y}_g, \sigma_g, \boldsymbol{\alpha}).$$

We treat the variation in the observed expression levels for a given gene, $\sigma_g$, as a random variable that depends on the true expression level of the gene, $\mathrm{Y}_g$, and three other parameters—$s_0, s_1, \tau$—that are shared among all genes. Accordingly, the probability of the observed variation in expression levels of gene $g$ across the libraries in which it is detected is $P(\sigma_g \mid \mathrm{Y}_g, s_0, s_1, \tau)$, and the observed variation in expression levels for all genes, $\boldsymbol{\sigma}$, has probability:

$$P(\boldsymbol{\sigma} \mid \mathbf{Y}, s_0, s_1, \tau) = \prod_{g=1}^{G} P(\sigma_g \mid \mathrm{Y}_g, s_0, s_1, \tau),$$

which assumes independence of $\sigma_g$ among genes, given $s_0$, $s_1$, and $\tau$.

The joint probability of the lower level (the first term on the right-hand side of equation S1) can therefore be written as:

$$P(\mathbf{X} \mid \mathbf{Y}, \theta_1)P(\theta_1) = \underbrace{P(\mathbf{X} \mid \mathbf{Y}, \boldsymbol{\sigma}, \boldsymbol{\alpha})}_{\text{sampling model}}\underbrace{P(\boldsymbol{\sigma} \mid \mathbf{Y}, s_0, s_1, \tau)}_{\substack{\text{mean-variance}\\\text{model}}}\underbrace{P(\boldsymbol{\alpha}, s_0, s_1, \tau)}_{\text{joint prior model}}, \tag{S2}$$

where the third term represents the joint prior distribution of the local model parameters (collectively contained in parameter $\theta_1$). We now consider each of these three terms in detail.

*Sampling model.*—The parameters of the sampling component of the hierarchical model are listed and described in in Table S2. A given gene in a given library is either detected (*i.e.*, with one or more reads) or not detected (*i.e.*, with zero reads). We denote the probability of detecting at least one read from gene $g$ in library $r$ as $\rho_{gr}$, which is a function of the true expression level of that gene, $\mathrm{Y}_g$. In principle, our sampling model should exhibit the following attributes: (1) the probability that a read maps to a given gene is proportional to the length of the transcript of that gene, $L_g$; (2) the probability that a read maps to a given gene is proportional to the expression level of that gene, $\mathrm{Y}_g$, and; (3) the probability that the gene will be detected in a given library is proportional to the sequencing depth of that library (*i.e.*, the total number reads in the library).

Accordingly, we model the distribution of the number of reads for a given gene as a Binomial random variable with parameters $n_r$ (the total number of reads in library $r$) and $p_g$ (the probability that a read maps to gene $g$). When $n_r$ is large and $p_g$ is small, the number of reads for gene $g$ in library $r$ is approximately Poisson distributed with rate parameter $\lambda \approx n_r p_g$. In practice, empirical datasets will deviate somewhat from the above expectations. Accordingly, we model $\lambda_{gr}$ as

$$\lambda_{gr} = \alpha_r L_g e^{Y_g},$$

where $\alpha_r$ represents library-specific factors (such as sequencing depth) that impact detection probability, $L_g$ describes the positive correlation between transcript length and detection probability, and $e^{Y_g}$ describes the positive correlation between the true expression level and detection probability. For a given value of $\lambda_{gr}$, we can compute the detection probability, $\rho_{gr}$ (*i.e.*, the probability of detecting at least one read for gene $g$ in library $r$), as:

$$\rho_{gr} = 1 - e^{-\lambda_{gr}}. \tag{S3}$$

For genes that are detected, we assume their expression level in a given library is normally distributed with mean $\mathrm{Y}_g$ and standard deviation $\sigma_g$, and that the observed expression levels among replicate libraries are independent conditioned on $e^{Y_g} > 0$ (see Fig. 1B, main text). Therefore, under this sampling model, the probability of the observed expression levels is:

$$P(\mathbf{X}_{g\cdot} \mid \mathrm{Y}_g, \sigma_g, \boldsymbol{\alpha}) = \begin{cases} \prod_{r=1}^{R} \mathbb{I}\left(\mathrm{X_{gr}}\right) & \text{if } e^{Y_g} = 0 \\ \prod_{r=1}^{R} \left[ (1-\rho_{gr})\delta(e^{X_{gr}}) + \rho_{gr} \frac{1}{\sqrt{2\pi\sigma_g^2}} \exp\left( \frac{-(X_{gr}-Y_g)^2}{2\sigma_g^2} \right) \right] & \text{if } e^{Y_g} > 0, \end{cases}$$

where $\delta(\cdot)$ is the Dirac delta function and

$$\mathbb{I}\left(\mathrm{X_{gr}}\right) = \begin{cases} 1 & \text{if } \mathrm{e^{X_{gr}}} = 0 \\ 0 & \text{otherwise,} \end{cases}$$

gives the probability that a gene is undetected in all libraries, given that its true expression level is zero, *i.e* $e^{Y_g} = 0$.

*Mean-variance model.*—The second term in equation S2 describes the relationship between the true expression level and the variance in observed expression levels among replicate libraries. The observed variation in $\boldsymbol{X_{g\cdot}}$ among the $R$ libraries is due to both biological variation (*i.e.*, replicate libraries are based on samples from different individuals) and technical error introduced by the RNA-seq pipeline. It is generally observed that log-transformed relative-expression levels, $\mathbf{X}$, exhibit a consistent mean-variance relationship; the variance in the expression level of a given gene between libraries is inversely proportional to its average observed expression level across all libraries. We therefore model the relationship between the variance in the expression level of gene $g$, $\sigma_g^2$, and the true expression of the gene, $Y_g$, with an exponential trend parameterized by $s_0$, and $s_1$.

Specifically, we assume that each $\sigma_g^2$ is distributed as a lognormal random variable with mean $\mu_g$ and variance $\tau$. To model the relationship between the variance in the observed expression level and the true expression level of gene $g$, we specify the mode of the lognormal distribution, $S_g$, as a function of $Y_g$:

$$S_g = \exp\left(s_0 + s_1 \mathrm{Y}_g\right).$$

The parameter $s_0$ represents the baseline variance in the expression level of all genes. The parameter $s_1$ describes the strength and direction of the relationship between the true expression level and variance (*e.g.*, $s_1 < 0$ implies that the variance decreases as the true expression level increases). The corresponding log-mean of the lognormal distribution for gene $g$, $\mu_g$, is:

$$\mu_g = s_0 + s_1 \mathrm{Y}_g + \tau.$$

We assume that each $\sigma_g^2$ is sampled independently from this lognormal distribution. Consequently, the probability of the gene-specific variances is:

$$P(\boldsymbol{\sigma} \mid \mathbf{Y}, s_0, s_1, \tau) = \prod_{g=1}^{G} \frac{1}{\sigma_g^2\sqrt{2\pi\tau}} \exp\left(-\frac{(\ln \sigma_g^2 - \mu_g)^2}{2\tau}\right).$$

Our parameterization jointly estimates the predictive value of the trend along with the individual gene-wise variances to provide robust classification estimates for even small datasets, *e.g.* two libraries.

Differential expression methods employ library-specific scaling factors in their tests [2–4]. In our model, this translates to the parameterization of the mean-variance trend-line, $S_g$, where the component of the variance trend that is independent of the expression level of genes is influenced by random effects such as scaling factors. If the scale of library expression levels is highly variable, then the trend line will be shifted upward. This should have similar impacts on estimates of expression state for genes.

*Joint prior model.*—The third and final term in equation S2 describes our prior beliefs regarding the values of the local model parameters. We assume independence of the prior probability distribution for each parameter in the lower level of the hierarchical model, so that we can write their joint prior density as:

$$P(\boldsymbol{\alpha}, s_0, s_1, \tau) = P(\boldsymbol{\alpha})P(s_0)P(s_1)P(\tau).$$

We assume that the library-specific parameters $\alpha_r$ are drawn independently from an exponential prior distribution with rate parameter $\beta^{\alpha}$, so that

$$P(\boldsymbol{\alpha} \mid \beta^{\alpha}) = \prod_{r=1}^{R} P(\alpha_r \mid \beta^{\alpha}).$$

We assume that $s_0$ is drawn from a normal prior distribution with mean $\mu^{s_0}$ and standard deviation $\sigma^{s_0}$. To express our prior belief that $s_1$ should be negative (*i.e.*, that the variance in observed expression levels scales inversely with true expression levels), we assume an exponential prior with parameter $\beta^{s_1}$ on the *negative* of $s_1$, so that $s_1 \leq 0$. Finally, we assume that the variation in the variance, $\tau$, is drawn from a gamma prior distribution with shape parameter $\alpha^{\tau}$ and rate parameter $\beta^{\tau}$.

##### S1.2.2 Upper Level

Recall that the upper level of our Bayesian hierarchical model describes the distribution of the true, unobserved expression levels, $\mathbf{Y}$. We imagine that the true expression levels of genes can be divided into two components; those genes that are actively expressed, and those that are not actively expressed. The assignment of each gene to the active or inactive expression-state components is described by parameter $z_g^a$, where $z_g^a = 1$ indicates that the gene is assigned to the active component, and $z_g^a = 0$ indicates that the gene is assigned to the inactive (*i.e.*, background) component. The central goal of this model is to infer $z_g^a$ for each gene in our dataset. Given the assigned expression state of gene $g$, we can compute the probability of its true expression level, $Y_g$, as:

$$P(Y_g \mid z_g^a, \theta_2) = \begin{cases} P(Y_g \mid \theta_i) & \text{if } z_g^a = 0 \text{ (inactive component)} \\ P(Y_g \mid \theta_a) & \text{if } z_g^a = 1 \text{ (active component),} \end{cases}$$

where $\theta_i$ are the parameters governing the distribution of inactive genes, and $\theta_a$ are parameters governing the distribution of active genes. We assume that—conditional on the assignment of genes to expression states—the $Y_g$ are independent. Therefore, the joint probability of all true expression levels, $\mathbf{Y}$, is:

$$P(\mathbf{Y} \mid \boldsymbol{z}^a, \theta_2) = \prod_{g=1}^{G} P(Y_g \mid z_g^a, \theta_2)$$

The joint probability of the upper level (the second term on the right-hand side of equation S1) can therefore be written as $P(\mathbf{Y} \mid \boldsymbol{z}^a, \theta_2) P(\boldsymbol{z}^a, \theta_2)$, where $P(\boldsymbol{z}^a, \theta_2)$ represents the joint prior distribution of the focal and local model parameters. The mixture-distribution parameters of the upper level are listed and briefly summarized in Table S3. We describe the inactive and active model components below.

*Inactive component.*—We imagine that the population of inactive genes can be subdivided into two subcomponents: one with zero expression, and another with non-zero expression (*e.g.*, as might result from promoter leakage). We model these subcomponents using a "spike-and-slab" mixture distribution, where the "spike" represents zero expression and the "slab" represents non-zero expression. The parameter $z_g^s$ indicates the allocation of inactive gene $g$ to these two subcomponents, where $z_g^s = 1$ indicates that the gene is assigned to the spike, and $z_g^s = 0$ indicates that the gene is assigned to the slab. We assume inactive genes in the slab are normally distributed with mean $\mu^b$ and standard deviation $\sigma^b$. Therefore, the probability of a given inactive gene is:

$$P(Y_g \mid z_g^a = 0, z_g^s, \mu^b, \sigma^b) = \begin{cases} \mathbb{I}(Y_g) & \text{if } z_g^s = 1 \text{ (the gene is in the spike)} \\ P(Y_g \mid \mu^b, \sigma^b) & \text{if } z_g^s = 0 \text{ (the gene is in the slab),} \end{cases}$$

where

$$\mathbb{I}(Y_g) = \begin{cases} 1 & \text{if } e^{Y_g} = 0 \\ 0 & \text{otherwise,} \end{cases}$$

describes the probability when the gene is in the spike, and

$$P(Y_g \mid \mu^b, \sigma^b) = \frac{1}{\sqrt{2\pi\sigma^{2b}}} \exp\left(-\frac{(Y_g - \mu^b)^2}{2\sigma^{2b}}\right)$$

describes the probability when the gene is in the slab.

*Active component.*—Genes within the active component may cluster into one or more subcomponents with qualitatively different expression levels. We therefore model the distribution of active genes as a mixture of $K$ normal distributions. The parameter $z_g^m$ indicates the allocation of active gene $g$ to one of these $K$ subcomponents; $z_g^m = k$ indicates that the gene is assigned to the $k^{\text{th}}$ active subcomponent. Active genes in the $k^{\text{th}}$ subcomponent are normally distributed with mean $\mu_k^m$ and standard deviation $\sigma_k^m$. Therefore, the probability of a given active gene is:

$$P(Y_g \mid z_g^a = 1, z_g^m = k, \boldsymbol{\mu}^m, \boldsymbol{\sigma}^m) = \frac{1}{\sqrt{2\pi\sigma_k^{2m}}} \exp\left(-\frac{(Y_g - \mu_k^m)^2}{2\sigma_k^{2m}}\right).$$

Several studies have shown that a normal distribution provides a good fit to the bulk of the empirical distribution of transcript abundance around the mode, but with tails that are heavier than expected in some cases, suggesting the need for additional active subcomponents [5, 6].

*Joint prior model.*—The joint prior model for the upper level of our hierarchical model includes the priors on the allocation parameters ($\boldsymbol{z}^a$, $\boldsymbol{z}^s$, $\boldsymbol{z}^m$), and the corresponding hyperparameters of those priors (*i.e.*, the hyperpriors $\boldsymbol{\omega}^a$, $\boldsymbol{\omega}^s$, and $\boldsymbol{\omega}^m$), and the priors on the various subcomponent parameters ($\mu^b$, $\sigma^b$, $\boldsymbol{\mu}^m$, $\boldsymbol{\sigma}^m$).

We assume that allocation parameter, $\boldsymbol{z}^a$, is drawn from a multinomial prior distribution with corresponding hyperparameters $\boldsymbol{\omega}^a$; $\omega_1^a$ is the prior probability that a gene is assigned to the inactive component, and $\omega_2^a$ is the prior probability that it is assigned to the active component. Rather than assume that the proportion of inactive and active genes is fixed, we treat the mixture weights $\boldsymbol{\omega}^a$ as random variables with corresponding hyperpriors. Specifically, we assume that $\boldsymbol{\omega}^a$ is drawn from a Dirichlet prior distribution with hyperparameters $\alpha^a$. We specify a similar structure on the other allocation parameters: $\boldsymbol{z}^s$ and $\boldsymbol{z}^m$ are each drawn from multinomial priors with corresponding hyperparameters $\boldsymbol{\omega}^s$ and $\boldsymbol{\omega}^m$, which in turn have hyperparameters $\alpha^s$ and $\alpha^m$.

Recall that we assume a normal distribution for the inactive genes with non-zero expression, *i.e.*, those in the "slab". We assume the mean of this normal distribution, $\mu_b$, is drawn from a gamma prior distribution with shape parameter $\alpha^b$ and rate parameter $\beta^b$, and the log of the variance, $\ln \sigma^{2b}$, is drawn from a uniform prior ranging from $a^b$ to $b^b$.

Similarly, recall that we assume a normal distribution for each of the $K$ subcomponents of active genes. We assume the mean of the $k^{\text{th}}$ subcomponent is drawn from a gamma prior distribution with shape parameter $\alpha^c$ and rate parameter $\beta^c$, plus a fixed offset value, $t_k$. The fixed offsets, $t_k$, are selected to minimize the overlap between each of the active subcomponents (Fig. S1). In principle, each active subcomponent might have its own variance parameter, $\sigma_k^{2m}$. For simplicity, we assume the $K$ active subcomponents share a variance parameter, $\sigma^{2m}$. We assume that the log of this variance parameter is drawn from uniform prior distribution ranging from $a^c$ to $b^c$.

Collectively, the joint prior density for the local parameters of the upper level of our hierarchical model is:

$$P(\boldsymbol{z}^a, \theta_2) = \underbrace{P(\boldsymbol{z}^a \mid \boldsymbol{\omega}^a)P(\boldsymbol{\omega}^a)}_{\substack{\text{active/inactive}\\ \text{mixture model}}} \underbrace{P(\boldsymbol{z}^s \mid \boldsymbol{\omega}^s)P(\boldsymbol{\omega}^s)P(\mu^b)P(\sigma^b)}_{\text{inactive mixture model}} \underbrace{P(\boldsymbol{z}^m \mid \boldsymbol{\omega}^m)P(\boldsymbol{\omega}^m)P(\boldsymbol{\mu}^m)P(\boldsymbol{\sigma}^{2m})}_{\text{active mixture model}}$$

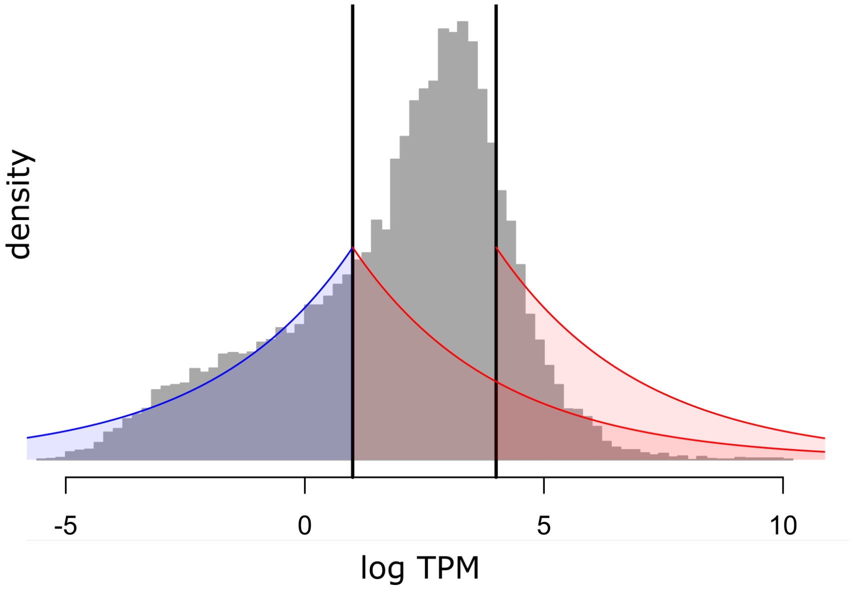


**Figure S1: Prior thresholds for the means of the in/active mixture components.** The gray distribution indicates the observed log-expression levels for a library; black vertical lines indicate the location of two thresholds that specify the upper boundary (in the case of the inactive component), and the lower boundary (in the case of active components). These thresholds represent our prior knowledge that the location of a given component mean does not exceed the threshold value. The red and blue shaded regions indicate the prior distributions for inactive component mean (blue) and the two active subcomponent means (red).

**Table S1: Notation for data variables of our hierarchical Bayesian mixture model.**

| Data Variable | Interpretation |
|---|---|
| $g$ | The index of a specific gene |
| $G$ | The number of genes |
| $r$ | The index of a specific library |
| $R$ | The number of libraries |
| $\mathbf{X}$ | All of the libraries, a $G \times R$ matrix |
| $\mathbf{X}_{g\cdot}$ | All of the transcripts for gene $g$ across libraries |
| $\mathbf{X}_{\cdot r}$ | All of the transcripts for library $r$ |
| $\mathbf{Y}$ | The true (unobserved) expression levels for all genes, a $G \times 1$ column vector |
| $Y_g$ | The true (unobserved) expression level for gene $g$ |
| $\mathbf{L}$ | The length of each transcript, a $G \times 1$ column vector |

**Table S2: Parameters of (and default values for) the sampling component of our hierarchical Bayesian mixture model.**

| Sampling Variable | Default Value | Interpretation |
|---|---|---|
| $\alpha$ | Gamma($\alpha^{\alpha} = 1, \beta^{\alpha} = 1/10$) | Library-specific parameter that is proportional to the number undetected genes in that library |
| $\sigma_g^2$ | | The variance in the observed expression level of gene $g$ among libraries |
| $s_0$ | Normal($\mu^{s_0} = -1, \sigma^{s_0} = 2$) | The base-line variance in expression levels |
| $s_1$ | $-s_1 \sim$ Gamma($\alpha^{s_1} = 1, \beta^{s_1} = 2$) | The influence of the true expression level, $Y_g$ on the variance in expression level among libraries |
| $\tau$ | Gamma($\alpha^{\tau} = 1, \beta^{\tau} = 1$) | The variation in gene-specific variances |

**Table S3: Parameters of (and default values for) the mixture component of our hierarchical Bayesian mixture model.**

| Mixture Variable | Default Value | Interpretation |
|---|---|---|
| $\omega^a$ | Dir($\alpha_1^a = 2, \alpha_2^a = 2$) | The prior probability that any gene is active or inactive |
| $\boldsymbol{z}^a$ | | The allocation of genes to the active component |
| $z_g^a$ | | The allocation of gene $g$ to the active component ($z_g^a = 1$) or the inactive component ($z_g^a = 0$) |
| $\omega^s$ | Dir($\boldsymbol{\alpha}^s = 1$) | The prior probability that an inactive gene is in the spike or slab |
| $\boldsymbol{z}^s$ | | The allocation of inactive genes among spike-and-slab subcomponents |
| $z_g^s$ | | The allocation of inactive gene $g$ to the spike ($z_g^s = 1$) or slab ($z_g^s = 0$) inactive subcomponents |
| $\mu^b$ | Gamma($\alpha^b = 1, \beta^b = 1/3$) | The mean (log) expression level for inactive genes, given that they are expressed |
| $\log \sigma^{2b}$ | Unif($a^b = \log 0.01, b^b = \log 5$) | The variance of the (log) expression level for inactive genes, given that they are expressed |
| $\omega^m$ | Dir($\alpha_1^m = \alpha_1^a/K, ..., \alpha_K^m = \alpha_1^a/K$) | The prior probability that an active gene is in each active subcomponents |
| $K$ | | The number of active subcomponents |
| $\omega_k^m$ | | The prior probability that an active gene is assigned to the $k^{\text{th}}$ active subcomponent |
| $\boldsymbol{z}^m$ | | The allocation of active genes among the active subcomponents |
| $z_g^m$ | | The allocation of active gene $g$ to an active subcomponent |
| $\boldsymbol{\mu}^a$ | Gamma($\alpha^c = 1, \beta^c = 1/3$) | The mean of the (log) expression level for active genes |
| $\mu_k^a$ | | The mean of the (log) expression level for active genes in subcomponent $k$ |
| $\boldsymbol{\log \sigma^{2a}}$ | Unif($a^c = \log 0.01, b^c = \log 5$) | The standard deviations of the (log) expression level for active genes |
| $\sigma_k^a$ | | The variance of the (log) expression level for active genes in subcomponent $k$ |

##### S1.2.3 Simulating Data Under the Model

Our motivation for developing the hierarchical Bayesian mixture model is to estimate the model parameters—particularly to the infer the in/active expression state of each gene (parameter $z^a$)—from the observed transcriptomic datasets. However, we can also use this model to simulate datasets for a given set of model parameters. When we use our hierarchical model for inference, we fix the values of our observed data and estimate (posterior-probability) distributions of the model parameters; conversely, when we use our model for simulation, we fix the values of the model parameters and generate distributions of simulated transcriptomic datasets (see Fig. 1, main text). Simulation may be useful both for characterizing the statistical behavior of the model, and also for assessing the adequacy (absolute fit) of the model to a given empirical dataset (we describe both applications elsewhere). Here, we describe a procedure for simulating transcriptomic datasets under our Bayesian hierarchical mixture model, given a particular set of parameter values, $\theta$ (except for $\sigma_g^2$). The procedure begins by simulating true expression levels for each gene according to the upper level of the hierarchical model, and then simulates observed expression levels for each library according to the lower level of the hierarchical model.

We simulate a transcriptome one gene at a time as follows:

(1) for gene $g$, assign the gene to the active component with probability $\omega_2^a$; otherwise, assign the gene to the inactive component

(2) if the gene is assigned to the active component, assign it to active subcomponent $k$ with probability $\omega_k^m$

(3) if the gene is assigned to the inactive component, assign it to the spike subcomponent with probability $\omega_1^s$; otherwise, assign the gene to the slab subcomponent

(4) draw a true expression level, $Y_g$ according to the parameter of the subcomponent to which the gene is assigned

(5) draw the variance in the observed expression level from the lognormal distribution given $Y_g$ and the remaining parameters, $s_0$, $s_1$, and $\tau$

(6) for each library $r$, the gene is detected with probability $\rho_{gr}$; otherwise, it is not detected

(7) if the gene is detected in a library, draw an observed expression level from a normal distribution with mean $Y_g$ and variance $\sigma_g^2$

(8) repeat the above procedure for each gene.

#### S1.3 Markov Chain Monte Carlo

The joint posterior density, $P(z^a, \theta, \mathbf{Y} \mid \mathbf{X})$ (equation S1), cannot be calculated analytically because the marginal likelihood, $P(\mathbf{X})$, is impossible to evaluate. Accordingly, we use Markov chain Monte Carlo (MCMC) to draw samples from the joint posterior distribution using the Metropolis–Hastings, Gibbs, and Green algorithms [7–10]. To do so, we specify a stationary discrete-time Markov chain with a state-space that includes all of the model parameters, and with transition rules that guarantee that the stationary distribution of the Markov chain is the joint posterior distribution of those model parameters.

To construct a Markov chain that draws samples from distribution $P(z^a, \theta, \mathbf{Y} \mid \mathbf{X})$, we initialize the state of the chain with random starting values for the vector of model parameters, which we represent with the generic parameter $\Theta = \{z^a, \theta, \mathbf{Y}\}$. Then, for each of many iterations, we propose a new set of parameter values, $\Theta'$, from a proposal distribution $f(\Theta)$ and "accept" the proposed parameter values (set $\Theta \to \Theta'$) with probability [8, 10]:

$$A = \min\left[1, \overbrace{\frac{P(\Theta' \mid \mathbf{X})}{P(\Theta \mid \mathbf{X})}}^{\substack{\text{posterior}\\ \text{ratio}}} \times \overbrace{\frac{f(\Theta)}{f(\Theta')}}^{\substack{\text{proposal}\\ \text{ratio}}} \times \overbrace{J(\Theta')}^{\text{Jacobian}}\right].$$

(For all of the moves we describe, the Jacobian is 1, so we omit it from the remainder of this section.) If this process is repeated for an adequate number of iterations, the sampled values of $\Theta$ constitute a valid—albeit autocorrelated—sample from the distribution $P(\Theta \mid \mathbf{X})$. Because the acceptance rule is based on the ratio of the posterior probabilities of the proposed and current states, the Metropolis–Hastings algorithm obviates the need to calculate the intractable marginal likelihood, $P(\mathbf{X})$.

For our current application, we initialize the chain by drawing a value for each model parameter from its corresponding prior distribution. For some initial states of the gene-wise parameters, the chain will not converge in a practical amount of time. To facilitate convergence, $\alpha_r$ is drawn from a Gamma$(1,1)$, $s_0$ from a Normal$(0,1/10)$, $s_1$ from a Gamma$(1,5)$, $\tau$ from a Gamma$(1,5)$ and for each gene $Y_g$ is drawn from Normal$(\bar{X}_g, s_g)$ which is parameterized by the observed gene-wise means and standard deviations in the data. If $X_g$ is detected in only one library, $s_g$ is drawn from a LogNormal$(\log(\bar{s}), \mathrm{sd}(s))$, with parameters determined by the mean and standard deviation of variances observed across all genes with detected expression in $X_g$. If $X_g$ is undetected in all libraries then $Y_g$ is drawn from the slab part of the inactive component *i.e.* from a Normal$(\mu^b, \sigma^b)$.

To propose a change to the state of the Markov chain we: (1) choose a single parameter in $\theta \in \Theta$; (2) propose a new value parameter value, $\theta'$, by drawing from a parameter-specific proposal distribution, $f(\theta)$; (3) and accept or reject the proposed value, $\theta'$, according to the acceptance rule, $A$, described above. We now describe each of the parameter-specific proposal distributions, $f(\theta)$.

*Proposals on $s_0$, $\mu^b$, $\mu_k^m$.*—All of these model parameters, which we refer to generically as $\theta$, can take values between $-\infty$ and $\infty$. To propose new parameter values, we draw $\theta'$ from a normal distribution centered on the current value:

$$\theta' \sim \text{Normal}(\theta, s),$$

where the variance $s$ represents a tuning parameter that controls the magnitude of the proposed changes to $\theta$: increasing/decreasing $s$ results in larger/smaller proposals. Because this proposal distribution is symmetric (*i.e.*, $f[\theta \mid \theta'] = f[\theta' \mid \theta]$), the acceptance probability is simply:

$$A = \min\left[1, \frac{P(\theta' \mid \Theta^c, \mathbf{X})}{P(\theta \mid \Theta^c, \mathbf{X})}\right],$$

where $\Theta^c$ refers to all of the remaining model parameters (*i.e.*, excluding $\theta$).

*Proposals on (log) $\sigma^{2b}$, $\sigma^{2m}$.*—Recall that we specify uniform priors on the log scale for variance parameters, which we refer to generically as $\theta$. We therefore propose changes to these parameters on the log scale (*i.e.*, $\theta$ is the log of the variance parameter). To propose new parameter values, we draw $\theta'$ from a uniform distribution centered on the current value:

$$\theta' \sim \text{Uniform}(\theta - \delta, \theta + \delta),$$

where $\delta$ represents a tuning parameter than controls the magnitude of the proposed changes to $\theta$: increasing/decreasing $\delta$ results in larger/smaller proposals. Because this proposal distribution is symmetric on the log scale (*i.e.*, $f[\theta \mid \theta'] = f[\theta' \mid \theta]$), the acceptance probability is simply:

$$A = \min \left[ 1, \frac{P(\theta' \mid \Theta^c, \mathbf{X})}{P(\theta \mid \Theta^c, \mathbf{X})} \right]$$

*Proposals on $\sigma_g^2$, $\alpha_r$, $s_1$, $\tau$.*—All of these model parameters, which we refer to generically as $\theta$, can take values $> 0$. To propose new parameter values, we draw a (log) scaling factor from a uniform distribution:

$$\begin{aligned} u &\sim \text{Uniform}(-\delta/2, \delta/2) \\ \theta' &= \theta e^u, \end{aligned}$$

where $\delta$ represents a tuning parameter that controls the magnitude of the proposed changes to $\theta$: increasing/decreasing $\delta$ results in larger/smaller proposals. Because this proposal distribution is asymmetric (*i.e.*, $f[\theta \mid \theta'] \neq f[\theta' \mid \theta]$), the acceptance probability is:

$$\begin{aligned} A &= \min \left[ 1, \frac{P(\theta' \mid \Theta^c, \mathbf{X})}{P(\theta \mid \Theta^c, \mathbf{X})} \times \frac{f(\theta)}{f(\theta')} \right] \\ &= \min \left[ 1, \frac{P(\theta' \mid \Theta^c, \mathbf{X})}{P(\theta \mid \Theta^c, \mathbf{X})} \times e^u \right]. \end{aligned}$$

*Joint proposal on $s_0$, and $\tau$.*—The posterior distribution of $s_0$ and $\tau$ often has a strong negative covariance, which can make independent moves inefficient for these two parameters. To improve efficiency, we generate a random uniform variate as above; $u$ which is then used to transform the two parameters:

$$\begin{aligned} \tau' &= \tau e^u \\ s_0' &= s_0 - u \end{aligned}$$

with a tuning parameter, $\delta$, specific to this combined move. The proposal ratio is also $e^u$ and the acceptance probability for $\boldsymbol{\theta'} = \{s_0', \tau'\}$ is:

$$\begin{aligned} A &= \min \left[ 1, \frac{P(\boldsymbol{\theta'} \mid \Theta^c, \mathbf{X})}{P(\boldsymbol{\theta} \mid \Theta^c, \mathbf{X})} \times \frac{f(\boldsymbol{\theta})}{f(\boldsymbol{\theta'})} \right] \\ &= \min \left[ 1, \frac{P(\boldsymbol{\theta'} \mid \Theta^c, \mathbf{X})}{P(\boldsymbol{\theta} \mid \Theta^c, \mathbf{X})} \times e^u \right]. \end{aligned}$$

*Proposal on $z_g^a$.*—A given gene belongs to both a component *and* a subcomponent, *e.g.*, a gene in the inactive component must belong to either the spike or the slab subcomponents, and a gene in the active component must belong to one of the $K$ active subcomponents. Therefore, in order to propose a change to the component to which a given gene belongs, we must simultaneously propose to reassign that gene to a new subcomponent. To propose reassignments of a given gene to either the active or inactive expression states, we first exhaustively evaluate the posterior probability of the assignments to all possible *subcomponents* (*i.e.*, both the active and inactive subcomponents). These joint posterior probabilities, $P(z_g^a, z_g^s, z_g^m \mid \mathbf{X}, \Theta^c)$ (conditional on the remaining model parameters $\Theta^c$) are:

$$\begin{aligned} P(z_g^a, z_g^s, z_g^m \mid \mathbf{X}, \Theta^c) &\propto P(\mathrm{Y}_g \mid z_g^a, z_g^s, z_g^m, \Theta^c) P(z_g^a, z_g^s, z_g^m \mid \Theta^c) \\ &\propto \underbrace{P(\mathrm{Y}_g \mid z_g^s, \Theta^c) P(z_g^s \mid z_g^a = 0, \Theta^c) P(z_g^a = 0 \mid \Theta^c)}_{\text{joint posterior probability of being inactive and } z_g^s} \\ &\quad + \underbrace{P(\mathrm{Y}_g \mid z_g^m, \Theta^c) P(z_g^m \mid z_g^a = 1, \Theta^c) P(z_g^a = 1 \mid \Theta^c)}_{\text{joint posterior probability of being active and } z_g^m}, \end{aligned}$$

where the sum in the above equation reflects the fact that the assignment of a given gene to the active or inactive components is mutually exclusive.

We then randomly propose a new set of assignments, $z_g^a$, $z_g^s$, and $z_g^m$, in proportion to their joint posterior probabilities:

$$f(z_g^a, z_g^s, z_g^m) = P(z_g^a, z_g^s, z_g^m \mid \mathbf{X}, \Theta^c).$$

The acceptance probability for this Gibbs proposal is:

$$\begin{aligned} A &= \min\left[1, \frac{\cancel{P(z_g^{a\prime}, z_g^{s\prime}, z_g^{m\prime} \mid \mathbf{X}, \Theta^c)}}{\cancel{P(z_g^a, z_g^s, z_g^m \mid \mathbf{X}, \Theta^c)}} \times \frac{\cancel{P(z_g^a, z_g^s, z_g^m \mid \mathbf{X}, \Theta^c)}}{\cancel{P(z_g^{a\prime}, z_g^{s\prime}, z_g^{m\prime} \mid \mathbf{X}, \Theta^c)}}\right] \\ &= 1. \end{aligned}$$

*Proposal on $z_g^s$.*—We propose reassignments of a given inactive gene between the spike or the slab, $z_g^s$, depending on whether it is currently in the spike or the slab. If gene $g$ is currently in the slab, we propose that $z_g^{s\prime} = 0$ and $Y_g^\prime = -\infty$. Otherwise, we propose that $z_g^{s\prime} = 1$ and draw a new value of $Y_g^\prime$ from the normal distribution that describes the slab and a new value of $\sigma_g^{2\prime}$ given $Y_g^\prime$ from a log normal distribution. Together, these rules allow us to compute the acceptance probability of reassigning an inactive gene $g$ from the spike to the slab as:

$$A = \min\left[1, \frac{P(\mathbf{X}_{g\cdot} \mid Y_g^\prime, \sigma_g^{2\prime}, \boldsymbol{\alpha}) P(Y_g^\prime \mid z_g^{s\prime} = 1, \mu^b, \sigma^{2b}) P(z_g^{s\prime} = 1 \mid \boldsymbol{\omega}^s)}{P(\mathbf{X}_{g\cdot} \mid Y_g, \sigma_g^2, \boldsymbol{\alpha}) P(Y_g \mid z_g^s = 0, \mu^b, \sigma^{2b}) P(z_g^s = 0 \mid \boldsymbol{\omega}^s)} \times \frac{1}{P(Y_g^\prime \mid z_g^{s\prime} = 1, \mu^b, \sigma^{2b}) P(\sigma_g^{2\prime} \mid Y_g^\prime)}\right].$$

Similarly, the acceptance probability of reassigning an inactive gene $g$ from the slab to the spike is:

$$A = \min\left[1, \frac{P(\mathbf{X}_{g\cdot} \mid Y_g^\prime, \sigma_g^2, \boldsymbol{\alpha}) P(Y_g^\prime \mid z_g^{s\prime} = 0, \mu^b, \sigma^{2b}) P(z_g^{s\prime} = 0 \mid \boldsymbol{\omega}^s)}{P(\mathbf{X}_{g\cdot} \mid Y_g, \sigma_g^2, \boldsymbol{\alpha}) P(Y_g \mid z_g^s = 1, \mu^b, \sigma^{2b}) P(z_g^s = 1 \mid \boldsymbol{\omega}^s)} \times \frac{P(Y_g \mid z_g^s = 1, \mu^b, \sigma^{2b}) P(\sigma_g^2 \mid Y_g)}{1}\right].$$

*Proposal on $z_g^m$.*—We propose reassignments of a given active gene to one of the $K$ subcomponents, $z_g^m$, by first exhaustively evaluating the posterior probability of all possible assignments, and then randomly propose a new assignment to an active subcomponent in proportion to its posterior probabilities. The proposal distribution, $f(z_g^m)$, is therefore:

$$f(z_g^m) = P(Y_g \mid z_g^m, \Theta^c) P(z_g^m \mid \Theta^c).$$

The acceptance probability for this Gibbs proposal is:

$$\begin{aligned} A &= \min\left[1, \frac{P(Y_g \mid z_g^{m\prime}, \Theta^c) P(z_g^{m\prime} \mid \Theta^c)}{P(Y_g \mid z_g^m, \Theta^c) P(z_g^m \mid \Theta^c)} \times \frac{f(z_g^m)}{f(z_g^{m\prime})}\right] \\ &= \min\left[1, \frac{\cancel{P(Y_g \mid z_g^{m\prime}, \Theta^c) P(z_g^{m\prime} \mid \Theta^c)}}{\cancel{P(Y_g \mid z_g^m, \Theta^c) P(z_g^m \mid \Theta^c)}} \times \frac{\cancel{P(Y_g \mid z_g^m, \Theta^c) P(z_g^m \mid \Theta^c)}}{\cancel{P(Y_g \mid z_g^{m\prime}, \Theta^c) P(z_g^{m\prime} \mid \Theta^c)}}\right] \\ &= 1 \end{aligned}$$

*Proposal on $\boldsymbol{\omega}^a$.*—To propose changes to the in/active mixture weights, $\boldsymbol{\omega}^a$, we first compute the number of genes that are assigned to each expression state:

$$n_i = \sum_{g=1}^{R} z_g^a = i,$$

such that the number of inactive genes is $n_1$ and the number of active genes is $n_2$. Recall that the prior distribution on the assignment of genes to the active or inactive expression states is multinomial, and that the prior distribution on the mixture weights is Dirichlet with parameter $\alpha^a$. The conditional posterior probability of $\boldsymbol{\omega}^a$ is:

$$P(\boldsymbol{\omega}^a \mid \boldsymbol{z}^a, \alpha^a) \propto P(\boldsymbol{z}^a \mid \boldsymbol{\omega}^a) P(\boldsymbol{\omega}^a \mid \alpha^a).$$

Because the Dirichlet distribution is conjugate to the multinomial distribution, this conditional posterior distribution is also a Dirichlet distribution:

$$\boldsymbol{\omega}^a \mid \boldsymbol{z}^a, \alpha^a \sim \text{Dirichlet}(\alpha^a + n_1, \alpha^a + n_2).$$

We therefore propose a new vector of mixture weights in proportion to their (conditional) posterior probability by drawing from this Dirichlet distribution, such that:

$$\begin{aligned} f(\boldsymbol{\omega}^a) &= P(\boldsymbol{\omega}^a \mid \boldsymbol{z}^a, \alpha^a) \\ &\propto P(\boldsymbol{z}^a \mid \boldsymbol{\omega}^a) P(\boldsymbol{\omega}^a \mid \alpha^a). \end{aligned}$$

The acceptance probability for this Gibbs proposal is:

$$\begin{aligned} A &= \min\left[1, \frac{P(\boldsymbol{z}^a \mid \boldsymbol{\omega}^{a\prime}) P(\boldsymbol{\omega}^{a\prime} \mid \alpha^a)}{P(\boldsymbol{z}^a \mid \boldsymbol{\omega}^a) P(\boldsymbol{\omega}^a \mid \alpha^a)} \times \frac{f(\boldsymbol{\omega}^a)}{f(\boldsymbol{\omega}^{a\prime})}\right] \\ &= \min\left[1, \frac{\cancel{P(\boldsymbol{z}^a \mid \boldsymbol{\omega}^{a\prime}) P(\boldsymbol{\omega}^{a\prime} \mid \alpha^a)}}{\cancel{P(\boldsymbol{z}^a \mid \boldsymbol{\omega}^a) P(\boldsymbol{\omega}^a \mid \alpha^a)}} \times \frac{\cancel{P(\boldsymbol{z}^a \mid \boldsymbol{\omega}^a) P(\boldsymbol{\omega}^a \mid \alpha^a)}}{\cancel{P(\boldsymbol{z}^a \mid \boldsymbol{\omega}^{a\prime}) P(\boldsymbol{\omega}^{a\prime} \mid \alpha^a)}}\right] \\ &= 1. \end{aligned}$$

*Proposal on $\boldsymbol{\omega}^s$*.—To propose changes to the spike-and-slab inactive mixture weights, $\boldsymbol{\omega}^s$, we first compute the number of inactive genes that are assigned to each of the inactive subcomponents:

$$n_i = \sum_{g=1}^{R} z_g^a = i,$$

such that the number of inactive genes in the spike is $n_1$ and the number of inactive genes in the slab is $n_2$. Recall that the prior distribution on the assignment of inactive genes to the spike or slab is multinomial, and that the prior distribution on the mixture weights is Dirichlet with parameter $\alpha^s$. The conditional posterior probability of $\boldsymbol{\omega}^s$ is:

$$P(\boldsymbol{\omega}^s \mid \boldsymbol{z}^s, \alpha^s) \propto P(\boldsymbol{z}^s \mid \boldsymbol{\omega}^s) P(\boldsymbol{\omega}^s \mid \alpha^s).$$

Because the Dirichlet distribution is conjugate to the multinomial distribution, this conditional posterior distribution is also a Dirichlet distribution:

$$\boldsymbol{\omega}^s \mid \boldsymbol{z}^s, \alpha^s \sim \text{Dirichlet}(\alpha^s + n_1, \alpha^s + n_2).$$

We therefore propose a new vector of mixture weights in proportion to their (conditional) posterior probability by drawing from this Dirichlet distribution, such that:

$$\begin{aligned} f(\boldsymbol{\omega}^s) &= P(\boldsymbol{\omega}^s \mid \boldsymbol{z}^s, \alpha^s) \\ &\propto P(\boldsymbol{z}^s \mid \boldsymbol{\omega}^s) P(\boldsymbol{\omega}^s \mid \alpha^s). \end{aligned}$$

The acceptance probability for this Gibbs proposal is:

$$\begin{aligned} A &= \min\left[1, \frac{P(\boldsymbol{z}^s \mid \boldsymbol{\omega}^{s\prime}) P(\boldsymbol{\omega}^{s\prime} \mid \alpha^s)}{P(\boldsymbol{z}^s \mid \boldsymbol{\omega}^s) P(\boldsymbol{\omega}^s \mid \alpha^s)} \times \frac{f(\boldsymbol{\omega}^s)}{f(\boldsymbol{\omega}^{s\prime})}\right] \\ &= \min\left[1, \frac{\cancel{P(\boldsymbol{z}^s \mid \boldsymbol{\omega}^{s\prime}) P(\boldsymbol{\omega}^{s\prime} \mid \alpha^s)}}{\cancel{P(\boldsymbol{z}^s \mid \boldsymbol{\omega}^s) P(\boldsymbol{\omega}^s \mid \alpha^s)}} \times \frac{\cancel{P(\boldsymbol{z}^s \mid \boldsymbol{\omega}^s) P(\boldsymbol{\omega}^s \mid \alpha^s)}}{\cancel{P(\boldsymbol{z}^s \mid \boldsymbol{\omega}^{s\prime}) P(\boldsymbol{\omega}^{s\prime} \mid \alpha^s)}}\right] \\ &= 1. \end{aligned}$$

*Proposal on $\boldsymbol{\omega}^m$*.—To propose changes to the active mixture weights, $\boldsymbol{\omega}^m$, we first compute the number of active genes that are assigned to each of the $K$ active subcomponents:

$$n_i = \sum_{g=1}^{R} z_g^a = i,$$

such that the number of active genes in the $k^{\text{th}}$ subcomponent is $n_k$. Recall that the prior distribution on the assignment of active genes to subcomponents is multinomial, and that the prior distribution on the mixture weights is Dirichlet with parameter $\alpha^m$. The conditional posterior probability of $\boldsymbol{\omega}^m$ is:

$$P(\boldsymbol{\omega}^m \mid \boldsymbol{z}^m, \alpha^m) \propto P(\boldsymbol{z}^m \mid \boldsymbol{\omega}^m) P(\boldsymbol{\omega}^m \mid \alpha^m).$$

Because the Dirichlet distribution is conjugate to the multinomial distribution, this conditional posterior distribution is also a Dirichlet distribution:

$$\boldsymbol{\omega}^m \mid \boldsymbol{z}^m, \alpha^m \sim \text{Dirichlet}(\alpha^m + n_1, \ldots, \alpha^m + n_K).$$

We therefore propose a new vector of mixture weights in proportion to their (conditional) posterior probability by drawing from this Dirichlet distribution, such that:

$$\begin{aligned} f(\boldsymbol{\omega}^m) &= P(\boldsymbol{\omega}^m \mid \boldsymbol{z}^m, \alpha^m) \\ &\propto P(\boldsymbol{z}^m \mid \boldsymbol{\omega}^m) P(\boldsymbol{\omega}^m \mid \alpha^m). \end{aligned}$$

The acceptance probability for this Gibbs proposal is:

$$\begin{aligned} A &= \min \left[ 1, \frac{P(\boldsymbol{z}^m \mid \boldsymbol{\omega}^{m\prime}) P(\boldsymbol{\omega}^{m\prime} \mid \alpha^m)}{P(\boldsymbol{z}^m \mid \boldsymbol{\omega}^m) P(\boldsymbol{\omega}^m \mid \alpha^m)} \times \frac{f(\boldsymbol{\omega}^m)}{f(\boldsymbol{\omega}^{m\prime})} \right] \\ &= \min \left[ 1, \cancel{\frac{P(\boldsymbol{z}^m \mid \boldsymbol{\omega}^{m\prime}) P(\boldsymbol{\omega}^{m\prime} \mid \alpha^m)}{P(\boldsymbol{z}^m \mid \boldsymbol{\omega}^m) P(\boldsymbol{\omega}^m \mid \alpha^m)}} \times \cancel{\frac{P(\boldsymbol{z}^m \mid \boldsymbol{\omega}^m) P(\boldsymbol{\omega}^m \mid \alpha^m)}{P(\boldsymbol{z}^m \mid \boldsymbol{\omega}^{m\prime}) P(\boldsymbol{\omega}^{m\prime} \mid \alpha^m)}} \right] \\ &= 1. \end{aligned}$$

#### S1.4 Implementation and Validation

We implemented all of the methods described in this paper in our open-source `R` package, `zigzag`, which is available at https://github.com/ammonthompson/zigzag. This implementation allows users to specify the (hyper)parameters of the hierarchical mixture model, conveniently summarize focal-parameter estimates, assess model adequacy using posterior-predictive simulation, perform MCMC diagnosis, and plot MCMC output.

##### S1.4.1 Specifying the Number of Active Subcomponents

Our Bayesian hierarchical mixture model includes $K$ active subcomponents of genes. Each active subcomponent is a normal distribution with an independent mean, $\mu_k^a$, and shared variance, $\sigma^{2a}$. The ability to specify models with variable numbers of active subcomponents raises two practical issues: (1) how to specify thresholds for each of the $K$ subcomponents, $t_k$, and; (2) how to select an appropriate value of $K$ for a given dataset.

Recall that $\mu_k^a$ is drawn from a Gamma prior with an offset value, $t_k$. For a model with $K$ active subcomponents, there are $K$ offset values, where the $k^{\text{th}}$ value specifies the lower boundary for the $k^{\text{th}}$ component mean. These offsets control the degree of overlap between adjacent active subcomponents, which facilitates identification of the individual subcomponents. For the analysis of empirical datasets, users specify offset values using an empirical Bayesian approach. Specifically, thresholds are specified based on visual inspection of the empirical expression levels for each library.

Each unique value of $K$ corresponds to a different inference model (*i.e.*, with $1, 2, \ldots, K$ active subcomponents). The relative fit of these candidate models to a given dataset can be compared using the posterior-predictive simulation approach described above. We recommend that users increment values of $K$ until the improvement in model adequacy plateaus. We provide examples of the procedures for specifying thresholds and assessing model adequacy in the `zigzag` documentation.

##### S1.4.2 Autotuning

In theory, a properly constructed MCMC will provide an arbitrarily precise approximation of the joint posterior probability distribution if the simulation is run for infinite time. For finite MCMC samples, the efficiency of the simulation depends on the properties of the proposal distribution, $f(\theta)$. For Metropolis–Hastings proposals, the sampling efficiency depends on the scale of proposed changes: when proposed changes are too large, they will be accepted infrequently, causing the chain to mix slowly over the joint posterior distribution. Conversely, when the proposed changes are too small, the acceptance rates will be too high, causing the chain to again mix slowly over the joint posterior distribution because of the granularity of the changes. Accordingly, acceptance rates of intermediate value provide more efficient sampling from the joint posterior probability distribution. In theory, the optimal acceptance rate for Metropolis–Hastings proposals is $\approx 44\%$ for scalar parameters [11].

We adopt an autotuning procedure that dynamically changes the scale of proposed changes (as determined by their tuning parameters, which we generically denote $\Delta$) until the acceptance rate reaches the target value of 44%. To achieve this, we use a "pre-burnin" phase during which the tuning parameters for Metropolis–Hastings proposals are iteratively adjusted based on the acceptance rate of a sliding window. Specifically, at pre-specified intervals during the pre-burnin phase, *e.g.*, every $i$ iterations, we compute the acceptance rate, $a$, for each proposal from the previous $j < i$ proposals. We then compute the new value of the tuning parameter, $\Delta'$, based on the current value of the tuning parameter $\Delta$, the acceptance rate within the previous window, $a$, and the target acceptance rate, $\hat{a}$ [11]

$$\Delta' = \Delta \times \frac{\tan(\pi \frac{a}{2})}{\tan(\pi \frac{\hat{a}}{2})}.$$

##### S1.4.3 Validation Experiment 1: Comparing Analytical and Numerical Prior Distributions

We performed tests to validate our implementation of the Metropolis–Hastings MCMC algorithm. Specifically, we used our MCMC algorithm to approximate the joint *prior* probability distribution. This experiment allows us to assess whether the MCMC algorithm is implemented correctly because it is targeting a *known* probability distribution; *i.e.*, the specified (known) prior distributions of the model parameters. We approximate the joint prior probability distribution by fixing the probability of the observed expression levels, $P(\mathbf{X} \mid \mathbf{Y}, \theta_1, \theta_2)$, to 1, regardless of the values of $\mathbf{Y}$, $\theta_1$ and ,$\theta_2$ (this is equivalent to running the MCMC simulation without data). After approximating the joint prior probability distribution, we simply compare the *estimated* marginal prior probability distribution for each parameter to the corresponding *analytical* (known) marginal prior distribution. If the MCMC algorithm is implemented correctly, the analytical and estimated prior distributions for each parameter will be identical. For this experiment, we assumed default values for all (hyper)parameters, and ran the MCMC for 240 million iterations, sampling every 4800 iterations. Our results confirm that the analytical and estimated marginal prior distributions are essentially identical (given the finite samples) (Fig. S2).

##### S1.4.4 Validation Experiment 2: Computing Coverage Probabilities

In a Bayesian model, the marginal posterior probability distribution for a given parameter reflects our beliefs about that parameter given the data. The $X$% credible interval (CI) of a marginal posterior distribution is the central interval that contains $X$% of that posterior distribution. If the model is correct, the true value of a parameter should be contained within the $X$% of its CI exactly $X$% of the time [12]. We validated the correctness of our MCMC algorithm by simulating data under the model and computing the frequency with which the true values of each parameter were contained within their respective 95% credible intervals.

For each of $i = 200$ simulated datasets, we first simulated the true parameter values, $\theta^i$, from their corresponding prior distributions. We diagnosed MCMC performance by computing the Effective Sample Size (ESS) and Potential Scale Reduction Factor (PSRF) diagnostics for all parameters for each analysis of the simulated datasets. All parameters for all analyses had ESS $> 100$, except for two cases (where ESS $> 80$), which we considered sufficient for this experiment. For each simulated dataset, we performed two replicate MCMC simulations, and computed PSRF values for each parameter from each pair of replicate analyses: all PSRF values were close to 1. For each of the $i$ datasets, we parameterized our model with the true parameter values, $\theta^i$, and simulated a corresponding transcriptomic dataset, $\mathbf{X}^i$. We then inferred the posterior distribution of model parameters for each simulated dataset, $P(\theta^i \mid \mathbf{X}^i)$. Finally, we computed the 95% credible interval for the marginal posterior distribution of each parameter, and measured the frequency with which the true value of each parameter was contained within its corresponding 95% credible interval (*i.e.*, we computed the coverage probability). These experiments confirm that the true parameter is contained in the 95% CI $\approx$ 95% of the time (Fig. S3).

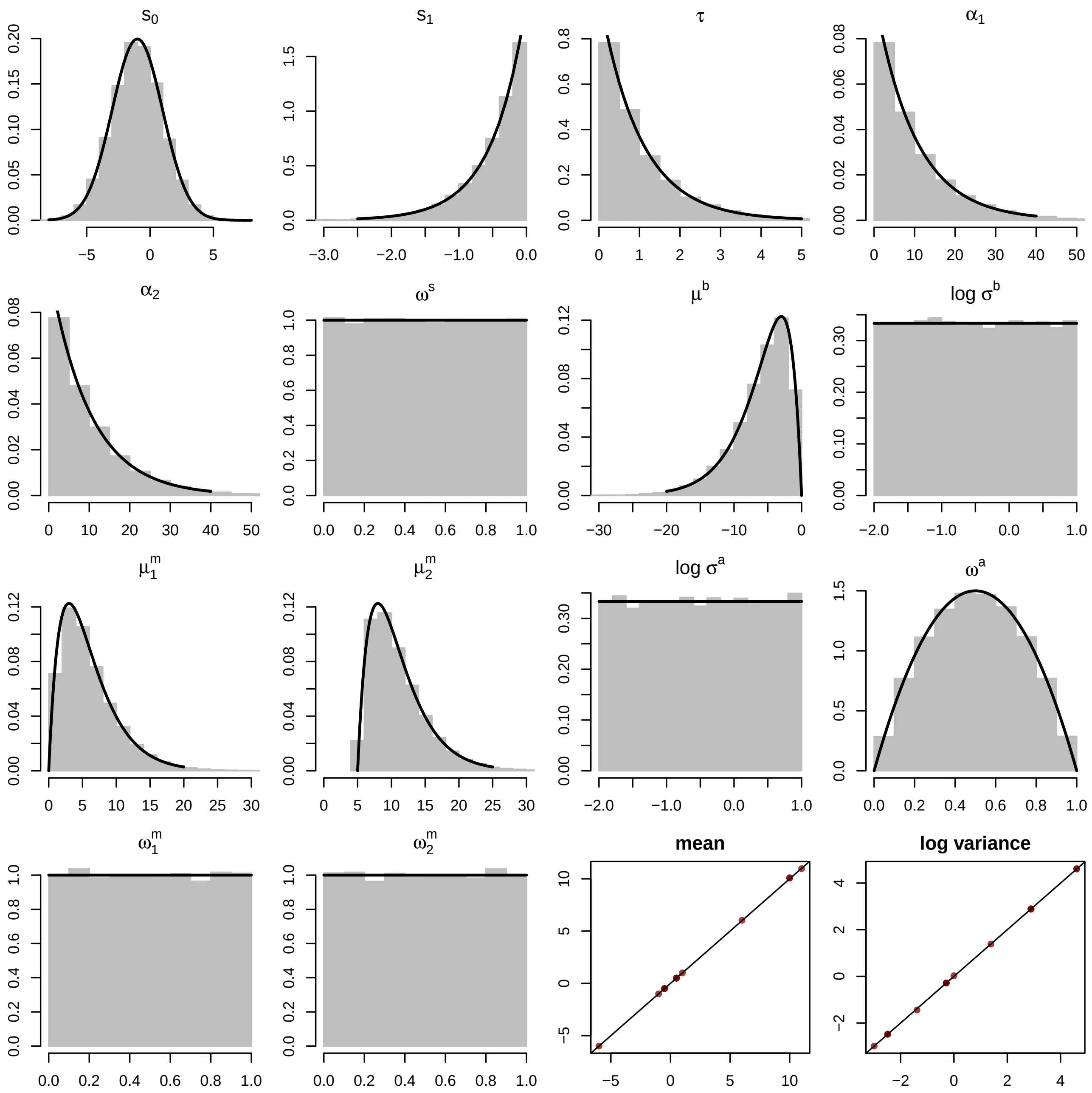


**Figure S2: Validating the MCMC implementation in `zigzag` by comparing estimated and analytical prior distributions.** We compared the analytical (known) prior probability densities (black lines) to the corresponding estimated marginal prior probability densities (gray histograms) for each parameter of our Bayesian hierarchical mixture model. The final two panels compare the means and log-variances of the estimated and known priors for all parameters; as expected of a correctly implemented MCMC algorithm, these values fall along the one-to-one line.

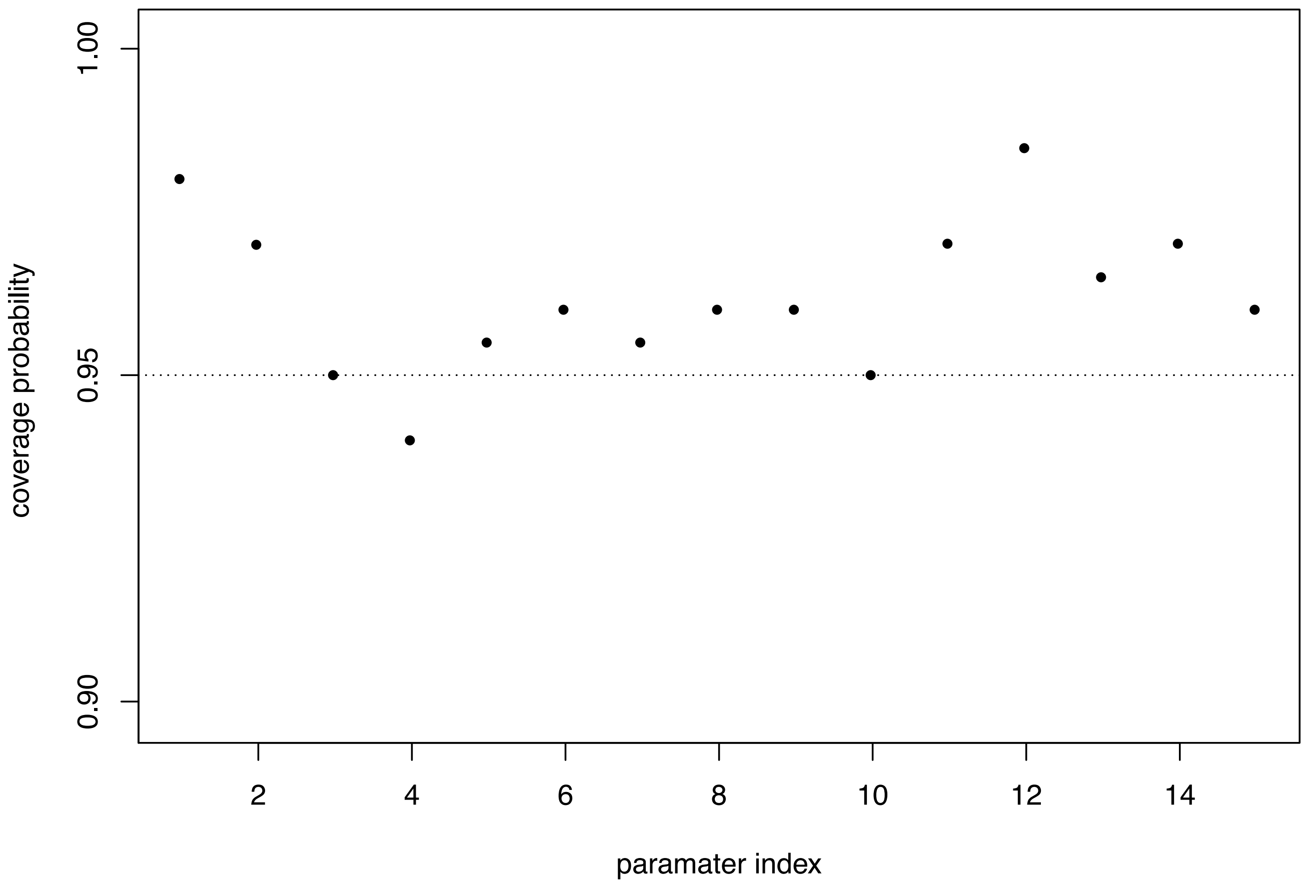


**Figure S3: Validating the MCMC implementation in `zigzag` by computing coverage probabilities.** We calculated the coverage probability for each of the 15 parameters in our hierarchical Bayesian mixture model by computing the frequency with which the true value of each parameter was contained within its corresponding 95% credible interval (dots).

#### S1.5 Assessing Model Adequacy Using Posterior Predictive Simulation

Model-based inference is based on the premise that our inference model provides an adequate description of the process that gave rise to our observations. The Bayesian approach for assessing model adequacy is called *posterior-predictive assessment* [13]. This approach is based on the following idea: if the inference model provides an adequate description of the process that gave rise to our observed data, then we should be able to use that model to simulate datasets that resemble our original dataset. We use a summary statistic to quantify the resemblance between the original and simulated datasets. By repeatedly simulating datasets from the joint posterior distribution of the model parameters inferred from the observed data, we construct a predictive distribution of summary statistics that characterize the resemblance between the simulated and observed data.

In outline, our procedure for assessing model adequacy involves the following steps:

(1) estimate the joint posterior probability distribution of the model parameters from the observed data, $P(\theta \mid \mathbf{X})$

(2) sample a vector of model parameters, $\theta_i$, from the joint posterior distribution

(3) compute $D_i^{\text{obs}}$, the "realized discrepancy" between the observed dataset, $\mathbf{X}$, and the fully specified model with parameters $\theta_i$ (we describe how to calculate the discrepancy below)

(4) simulate a dataset, $\mathbf{X}_i^{\text{sim}}$, equal in size to the original (with the same number of genes and replicate libraries, etc.)

(5) compute $D_i^{\text{sim}}$, the "realized discrepancy" between simulated dataset, $\mathbf{X}_i^{\text{sim}}$, and the fully specified model with parameters $\theta_i$

(6) compute the summary statistic as the difference of the two discrepancies, $S_i = D_i^{\text{obs}} - D_i^{\text{sim}}$

(7) repeat steps 2 through 6 $n$ times

The $n$ replicates collectively comprise a posterior-predictive distribution of summary statistics, $S$. When a simulated dataset resembles the observed dataset, the corresponding summary statistic (the difference in the discrepancy values for the observed and simulated datasets) will be close to zero. Accordingly, the 95% posterior-predictive interval of $S$ will include zero when the model is adequate. We can quantify the degree of model adequacy by computing the posterior-predictive $p$-value, which is simply the fraction of summary statistics that is less than or equal to zero.

We developed summary statistics that allow us to assess the adequacy of both the lower and upper levels of our hierarchical Bayesian mixture model. Specifically, we assess model adequacy using three summary statistics: (1) the lower-level Wasserstein statistic, which measures the discrepancy between the observed expression levels for detected genes, $\mathbf{X}$, and the expected expression levels for detected genes given the model parameters, $E[\mathbf{X} \mid \theta]$; (2) the upper-level Wasserstein statistic, which measures the discrepancy between the true expression levels $\mathbf{Y}$ and the parameters of the upper level of the mixture model, $\theta_2$, and; (3) the Rumsfeld statistic, which measures the discrepancy between the fraction of undetected transcripts in each library and the expected fraction of undetected transcripts in each library given the model parameters. We describe how to calculate these three summary statistics below.

##### S1.5.1 The Upper-Level Wasserstein Statistic

The Wasserstein metric measures the distance between two cumulative distributions (Fig. S4); we use it here to measure the discrepancy between the distribution of *measured* expression levels for detected genes ($e^{X_{gr}} > 0$) and the distribution of the *expected* expression levels of those genes given the model parameters, $\theta_i$.

Because the focal parameter, $z_g^a$, is most directly influenced by the unobserved true expression of genes, $Y_g$, the central statistic for assessing model adequacy is the $W^{\mathrm{U}}$ statistic. This statistic measures the discrepancy between the posterior distribution of $\mathbf{Y}$ and the local parameters of the upper level of our hierarchical model, $\theta_2$, for a given posterior sample, $i$.

The cumulative distribution of $\mathbf{Y}$ is:

$$F(y) = \frac{1}{G} \sum_{g=1}^{G} \left[ \mathrm{Y}_g < y \right].$$

and the cumulative distribution implied by $\theta_{2,i}$ is:

$$H(y \mid \theta_{2,i}) = \int_{-\infty}^{y} h(y \mid \theta_{2,i}) \mathrm{d}y,$$

where $H(y \mid \theta^{2,i})$ is the $i^{\mathrm{th}}$ sample from the posterior distribution. We measure the Wasserstein discrepancy between $Y$ and the sampled local model parameters, $\theta_{2i}$, as:

$$D^{\mathrm{U}}(Y) = \int_{-\infty}^{\infty} \left| F(y) - H(y \mid \theta_{2,i}) \right| \mathrm{d}y.$$

We then compute the difference between the Wasserstein discrepancies measured for simulated $Y$ and $Y_i$ sampled from the posterior distribution (Fig. S4):

$$W_i^{\mathrm{U}} = D^{\mathrm{U}}(\mathbf{Y}_i) - D^{\mathrm{U}}(\mathbf{Y}^{sim})$$

##### S1.5.2 The Lower-Level Wasserstein Statistic

Next, we compute the *cumulative* distribution function for *measured* expression levels for a given dataset, $\mathbf{X}$. The cumulative distribution function for a given library, $r$, describes the probability that a measured expression level is less than or equal to a given value, $x$, which we compute as:

$$F_r(x) = \frac{1}{G^*} \sum_{g^*=1}^{G^*} \left[ X_{g^* r} < x \right],$$

where $g^*$ is the subset of genes that have non-zero expression in the library (and $G^*$ is the number of genes with non-zero expression in the library).

Next, we compute the cumulative distribution function for the *expected* expression levels for a given dataset, $\mathbf{X}$, given the $i^{\mathrm{th}}$ posterior sample of lower-level model parameters, $\theta_{1,i}$. For a given library, we compute this as:

$$E_r(x \mid \theta_i) = \frac{1}{G} \sum_{g=1}^{G} \rho_{gr} \left[ \mathrm{Y}_{g,i} < x \right],$$

because the expected expression level for a detected gene is equal to its true (unobserved) expression level, $\mathrm{Y}_{g,i}$, and the probability that it is detected is $\rho_{gr}$ (equation S3).

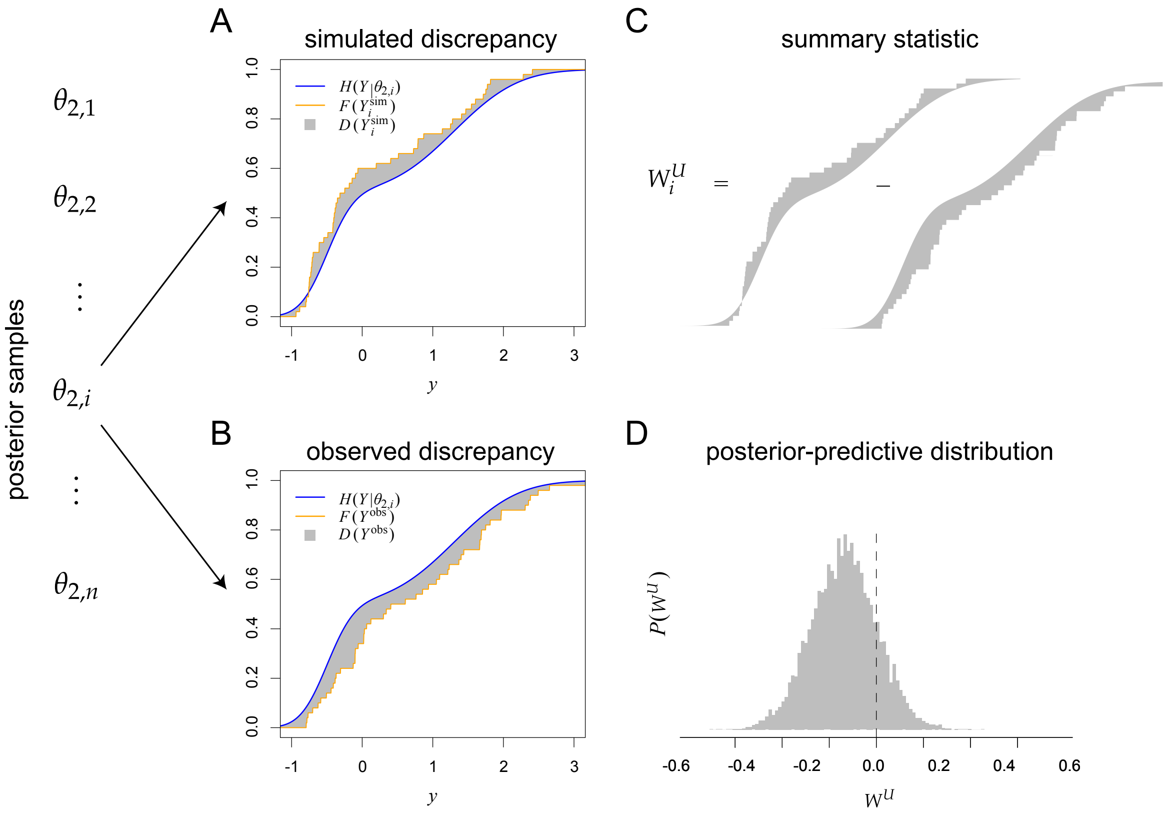


**Figure S4: Using posterior-predictive simulation to assess adequacy of the upper level of the hierarchical mixture model.** We imagine that we have previously performed MCMC simulation to draw $n$ samples from the joint posterior distribution. We then assess the adequacy of the upper level of our hierarchical model as follows. A. For the $i^{\text{th}}$ sample, we compute the *simulated* discrepancy by: (1) simulating a vector of true expression levels, $\text{Y}_i^{\text{sim}}$, and calculating the corresponding empirical cumulative distribution (orange curve). (2) Next, we compute the *expected* cumulative distribution of expression levels given the sampled parameter values, $\theta_{2,i}$ (blue curve). (3) We then compute the simulated discrepancy as the distance between the orange (simulated) and blue (expected) curves (gray area). B. We perform a similar procedure to compute the *observed* discrepancy by: (1) computing the empirical cumulative distribution function of the *sampled* true expression levels, $\text{Y}_i$ (orange curve). (2) We use the same *expected* cumulative distribution of expression levels computed previously (blue curve). (3) We then compute the simulated discrepancy as the distance between the orange (observed) and blue (expected) curves (gray area). C. Finally, we compute the upper-level Wasserstein summary statistic $W_i^U$ as the difference between the simulated and observed discrepancies. D. We repeat the previous steps for each of the $n$ samples to generate the posterior-predictive distribution of $W_i^U$. The model is considered adequate if the 95% posterior-predictive interval includes zero.

The Wasserstein discrepancy is the absolute value of the average distance between $F_r(x)$ and $E_r(x \mid \theta_{1,i})$ among $R$ replicate libraries:

$$\overline{w}^{\text{L}}(x) = \left| \frac{1}{R} \sum_{r=1}^{R} \left[ F_r(x) - E_{r,i}(x \mid \theta_{1,i}) \right] \right| .$$

By integrating over all values of $x$ we compute the discrepancy:

$$D^{\text{L}}(\mathbf{X} \mid \theta_i) = \int_{-\infty}^{\infty} \overline{w}^{\text{L}}(x) \mathrm{d}x.$$

This statistic evaluates the similarity between the average gene-expression level in the data and the expected distribution implied by the true expression level for the $i^{\text{th}}$ sample, $\mathbf{Y}_i$.

The resulting Wasserstein summary statistic for simulated dataset $\mathbf{X}_i^{\text{sim}}$ is then:

$$W_i^L = D^L(\mathbf{X} \mid \theta_{1,i}) - D^L(\mathbf{X}_i^{\text{sim}} \mid \theta_{1,i}). \tag{S4}$$

##### S1.5.3 The Rumsfeld Statistic

The Rumsfeld metric measures the distance between the fraction of undetected transcripts and the *expected* fraction of undetected transcripts given the model parameters, $\theta_i$ and $\mathbf{Y}_i$. We compute this statistic as follows. First, we compute the fraction of undetected genes in a given library, $U_r$. Next, we compute the expected fraction of undetected genes in that library, given the model parameters $\theta_i$. Genes are expected to go undetected in a library either because the true expression is zero, or because the transcripts are not sampled (equation, S3). The probability that a gene is undetected in a library because it has a true expression level of zero, denoted here as $P_i^s$, is:

$$P_i^s = P(e^{Y_{g,i}} = 0 \mid \theta_i) = \omega_i^s(1 - \omega_i^a).$$

The expected fraction of genes that have zero expression in a given library is thus computed as:

$$E_r(U \mid \theta_i) = P_i^s + (1 - P_i^s)\frac{1}{G}\sum_{g=1}^{G} \exp(-\alpha_r L_g e^{Y_{g,i}}).$$

The Rumsfeld metric is the absolute difference between $U_r$ and $E_{r,i}(U \mid \theta_i)$, averaged over all $R$ replicate libraries:

$$D^R(\mathbf{X} \mid \theta_i) = \frac{1}{R}\sum_{r=1}^{R} \left| U_r - E_{r,i}(U \mid \theta_i) \right|.$$

The resulting Rumsfeld summary statistic for simulated dataset $\mathbf{X}_i^{\text{sim}}$ is then:

$$R_i = D^R(\mathbf{X} \mid \theta_i) - D^R(\mathbf{X}_i^{\text{sim}} \mid \theta_i). \tag{S5}$$

### S2 Analyses

#### S2.1 General Analysis Protocol

In this study, we performed analyses of simulated and empirical datasets. Here, we describe our general analysis protocol: specifically, we describe aspects of the prior specification, MCMC simulation, MCMC diagnosis, and model comparison that were common to all of the analyses in this study. Details unique to specific analyses are described where appropriate.

*Prior specification.*—We used default (hyper)prior values for all model parameters (Tables S2–S3).

*MCMC settings.*—For each analysis, we performed two replicate MCMC simulations to estimate the joint posterior distribution. Each MCMC simulation consisted of two phases: a pre-burnin phase, during which tuning parameters were adjusted to achieve an acceptance rate of 44%, and a sampling phase, during which we drew samples from the joint posterior distribution. We specified a pre-burnin phase of 10,000 generations, where each generation consists of 24 independent moves (roughly the number of parameters in the model) with a tuning interval of 200 generations; at each tuning interval, we computed the acceptance rate for each proposal from the 100 most recent proposals. We specified a sampling phases that ranged from 40,000 to 240,000 generations depending on the dataset, thinning the chain by sampling every 20 or 50 generations.

*MCMC diagnosis.*—We assessed the reliability of each MCMC simulation by computing the Effective Sample Size (ESS) diagnostic using the `R` package `coda` [14]; we considered an MCMC to have failed if the ESS value for any parameter was less than 100. For empirical analyses, we compared the joint posterior distributions obtained from each of the two replicate MCMC simulations using the potential scale reduction factor (PSRF) diagnostic [15]. We considered a set of replicate MCMC simulations to have failed if PSRF $> 1.2$; in such cases, we reran the effected analyses until we achieved an acceptable value of the PSRF diagnostic.

#### S2.2 Simulation Studies

We performed simulation studies to understand the statistical behavior of our hierarchical Bayesian mixture model. We expect that the ability to correctly assign genes to the active or inactive expression-state depends on the disparity between the true active and inactive components, the number of replicate libraries, and the fit of the model to the data (*i.e.*, the number of true and assumed active subcomponents, $K$). We performed experiments to understand: (1) the power to discriminate between in/active expression state as a function of the degree of overlap between true active and inactive distributions and the number of replicate libraries; (2) the impact of model (mis)specification—*i.e.*, the assumed and true number of active subcomponents—on the ability to assign genes to the correct expression state, and; (3) the ability of summary statistics to assess model fit with posterior-predictive simulation.

For each experiment, we simulated 100 datasets, each consisting of 5000 genes. To ensure the realism of the simulated datasets, we drew simulating model parameters from distributions that resembled the corresponding marginal posterior distributions from our empirical analyses (see below). In particular, we drew the parameter values as follows:

$$\begin{aligned} \boldsymbol{\omega}^a &\sim \text{Dirichlet}(10, 10) \\ \boldsymbol{\omega}^s &\sim \text{Dirichlet}(5, 20) \\ \boldsymbol{\omega}^m &\sim \text{Dirichlet}(20, 5) \\ \tau &\sim \text{Uniform}(0.5, 1.5) \\ s_0 &\sim \text{Uniform}(-2.5, -1.5) \\ s_1 &\sim \text{Uniform}(-1.0, -0.1) \\ \alpha_r &\sim \text{Uniform}(5, 30) \end{aligned}$$

The remaining model parameters, $\mu^b, \boldsymbol{\mu^a}, \sigma^{2b}, \sigma^{2a}$, were set to values specific for each of the experiments, detailed below. For each gene, we sampled a gene length, $L_g$, from the distribution of gene lengths for the human genome (GRCH 38).

*Experiment 1: Power.*—This experiment explored the ability to correctly assign genes to the (in)active expression state, where we simulated datasets with differential degrees of overlap, determined by the variance of the inactive and active components, $\sigma^{2b}$ and $\sigma^{2a}$, respectively. We assumed that the mean of the inactive component was $\mu^b = -2$, and that there were two active subcomponents with means $\mu_1^a = 2$ and $\mu_2^a = 6$. We varied the component variances over three values, $\sigma^{2b} = \sigma^{2a} = \{1, 2, 3\}$ with prior on log $\sigma^{2a,b}$ set to $\text{Unif}(\log 0.1, \log 8)$, and the number of replicate libraries over three values, $R = \{2, 4, 6\}$. We analyzed each simulated dataset using the generic protocol described above. We measured power by computing the fraction of genes that were correctly assigned to their true in/active expression state. Results show that increasing number of replicate libraries improves accuracy, while the the amount of overlap between distributions imposes an intrinsic upper limit on the ability to distinguish between active and inactive genes (Fig. 2, main text).

*Experiment 2: Model Misspecification.*—This experiment explored the ability to correctly assign genes to the in/active expression state when the model was either underspecified (*i.e*, where the assumed number of active subcomponents is less than the true number) or overspecified (*i.e*, where the assumed number of active subcomponents exceeds the true number). We expect the impact of model misspecification to depend on the disparity between the generating and inference model. For example, if the true model has two active subcomponents, $K = 2$, that are quite similar, we expect that an underspecified inference model with one active subcomponent should perform reasonably well at inferring the true expression state of each gene. Conversely, if the two active subcomponents are very distinct, the underspecified model should perform poorly. We explored these predictions by simulating datasets with $K = 1$ or 2 active subcomponents and analyzing them using underspecified and overspecified models; the underspecified analyses assumed $K = 1$ when analyzing data generated with $K = 2$, and the overspecified analyses assumed $K = 2$ when the analyzing data generated with $K = 1$. Further, we assumed that the mean of the inactive component was $\mu^b = -2$, the mean of the first active subcomponent was $\mu_1^a = 2$, and the variance of each component was $\sigma^{2b} = \sigma^{2a} = 2$. To simulate increasingly distinct active subcomponents, we iteratively increased the mean of the second active subcomponent over three values, $\mu_2^b = \{3, 4, 6\}$. Each simulated dataset consisted of $R = 4$ replicate libraries. We analyzed each simulated dataset using the generic protocol described above. We assessed the consequences of model misspecification by computing the average probability genes were correctly assigned to the true in/active expression state. Our results indicate that model overspecificaiton (assuming too many active subcomponents) has virtually no impact on estimates of expression state. Similarly, we found that underspecification (assuming too few active subcomponents) has no detectable impact on estimates: when the active subcomponents strongly overlap, a single active subcomponent provides a reasonable approximation. However, estimation of expression state becomes increasingly impacted as the active subcomponents become increasingly distinct (Fig. 2, main text).

##### S2.3 Empirical Studies

For all empirical analyses, we assumed default priors for all parameters with three exceptions: (1) allocation weights for each component had equal prior probabilities: $\{1 - \omega^a, \omega_1^m, ..., \omega_K^m\} \sim \text{Dir}(2, 2, ..., 2)$; (2) $s_0 \sim N(0, 10)$, and; (3) $s_1 \sim \text{Gamma}(1, 1/10)$. We iteratively increased the number of active subcomponents until we achieved model adequacy (assessed using posterior-predictive simulation). We specified the threshold for an additional active subcomponent based on inspection of the posterior-predictive distribution simulated under the simpler, immediately adjacent model. For example, if posterior-predictive simulation indicated that a model with $k = 2$ active subcomponents was inadequate, we specified a new, more complex model with $k = 3$ active subcomponents, where the threshold for the third active subcomponent was located at the point where the posterior-predictive distribution departed the most strongly from the observed distribution. [Note that we also evaluated models with mean offsets, rather than fixed thresholds; these analyses produced identical results, but the MCMC simulations were less efficient. Accordingly, we report results only for models with fixed thresholds.]

*Posterior-predictive checking.*—For each posterior sample, we simulated 1,000 datasets from the posterior distribution and computed posterior-predictive statistics as described above.

*Assessing sensitivity of expression-state estimates to the assumed model.*—To determine the sensitivity of focal-parameter estimates to the assumed number of active subcomponents, we compared estimates of $\text{P}(z_g^a \mid X)$ between each model. We $P(\boldsymbol{z}^a | \boldsymbol{X}, M_k)$ plotted against $P(\boldsymbol{z}^a | \boldsymbol{X}, M_{k+1})$ for all genes. These comparisons indicate intermediate probabilities are most sensitive to model complexity.

###### S2.3.1 Analyses of the Human-Lung Transcriptomic Dataset

*Transcriptomic data.*—We downloaded human-lung transcriptomic data from the GTEx project database (https://gtexportal.org/home/) [16–18]. We wrote a custom script that computed the average length of each gene from all corresponding transcript lengths in the Gencode V. 19 annotation.

*Epigenomic data.*—Epigenomic data consist of myriad chromatin signatures associated with the expression state of genes, such as histone-tail acetylation and DNAse digestion sensitivity. The Roadmap Epigenomics Project used machine learning to classify 15 discrete states of non-overlapping 200 bp windows in the genome based on a large number of chromatin epigenetics marks [19, 20]. These epigenetics marks include histone modifications, DNA methylation, and DNA accessibility that were used to annotate the regulatory state at each 200 bp window in the genome. The results of these annotations are publicly available in CoreMarks mnemonics bed files (http://www.roadmapepigenomics.org/). These mnemonics are described in [21] (see Fig. S5). We downloaded CoreMarks mnemonics bed files from the Roadmap Epigenomics Project database; these mnemonics are based on coordinates relative to human annotation version GRCH37 [21] for lung data.

We used the `closest` function in the `Bedtools` suite to count all of the chromatin mnemonics that overlapped with exon coordinates for each gene. We defined active genes as those with at least one active promoter or promoter-flanking mark overlapping its exons (states 1 and 2), at least one active transcription mark (states 3 through 5), and no marks associated with repressed expression (states 9 through 15) [21]. We defined inactive genes as those having no active promoter or transcription marks (states 1 through 5) and at least one heterochromatin, bivalent enhancer/TSS, Polycombrepressed, or quiescent mark (states 9 through 15) overlapping at least one exon. Of the 19,154 protein-coding genes in the human-lung transcriptome, we were able to classify 62% of the genes as either active

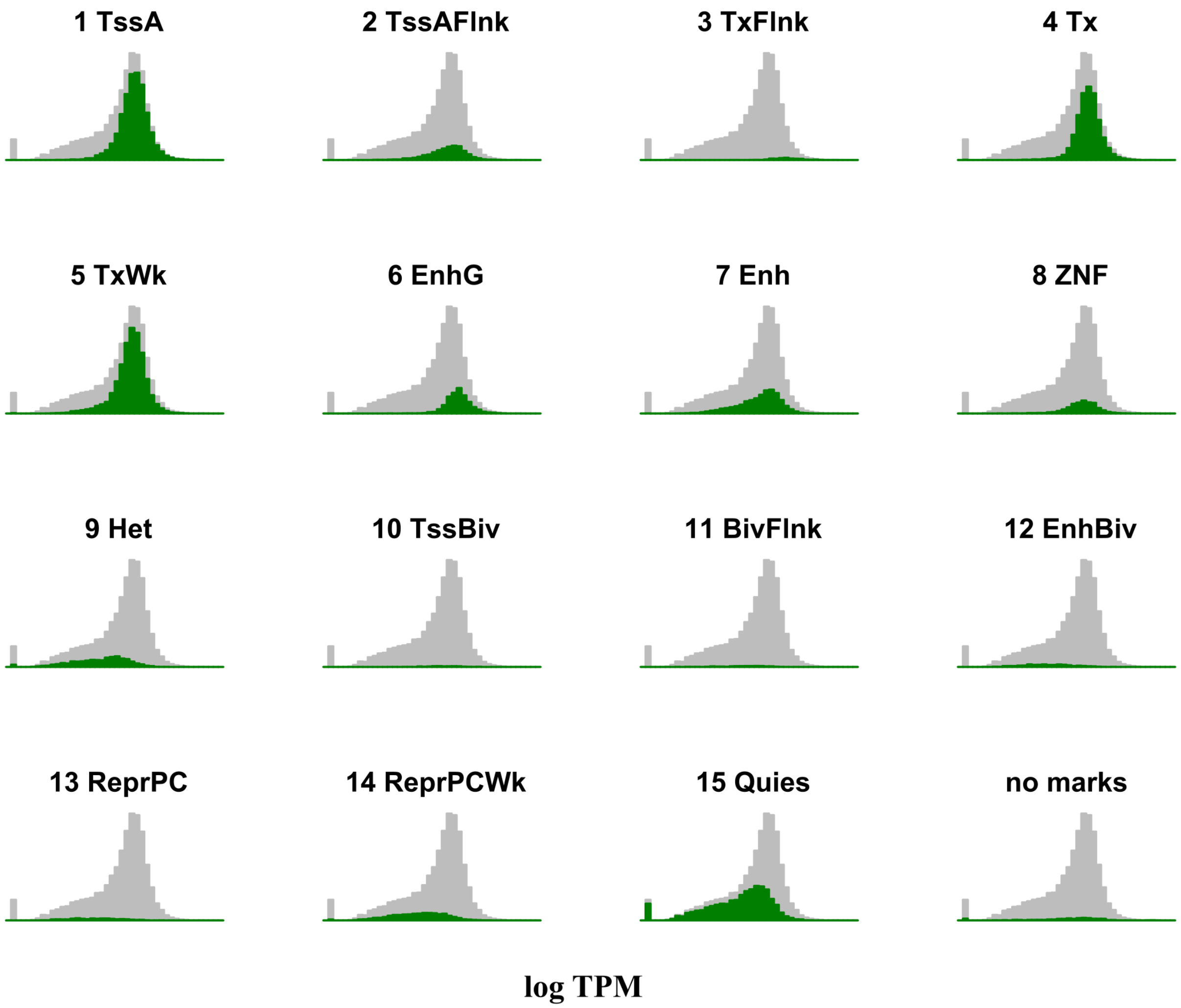


**Figure S5: Expression levels for genes in the human-lung transcriptome exhibiting each of the 15 epigenetic marks**. Expression-level distributions of genes that contain at least one exon with the corresponding epigenomic mark (green) overlayed against the expression level for the overall transcriptome in the GTEx dataset [16]. These 15 subsets of genes are not mutually exclusive; genes often have more than one type of epigenetic mark overlapping at least one exon.

(7,261) or inactive (4,707). Expression levels of active and inactive genes defined in this way have distinct unimodal distributions (Fig. S5).

*Model selection.*—Posterior-predictive simulation indicated that the improvement in model adequacy plateaued for model with two or more active subcomponents (Fig. S6).

*Sensitivity analyses.*—As in our simulation study, our analyses of the human-lung dataset indicate that expression-state estimates under the hierarchical Bayesian mixture model are robust to model overspecification: estimates under models with two, three or four active subcomponents are virtually identical (Fig. S7). Conversely, expression-state estimates under a model with one active subcomponent depart quite strongly from those inferred with two or more active subcomponents, especially for genes with intermediate probabilities of being active.

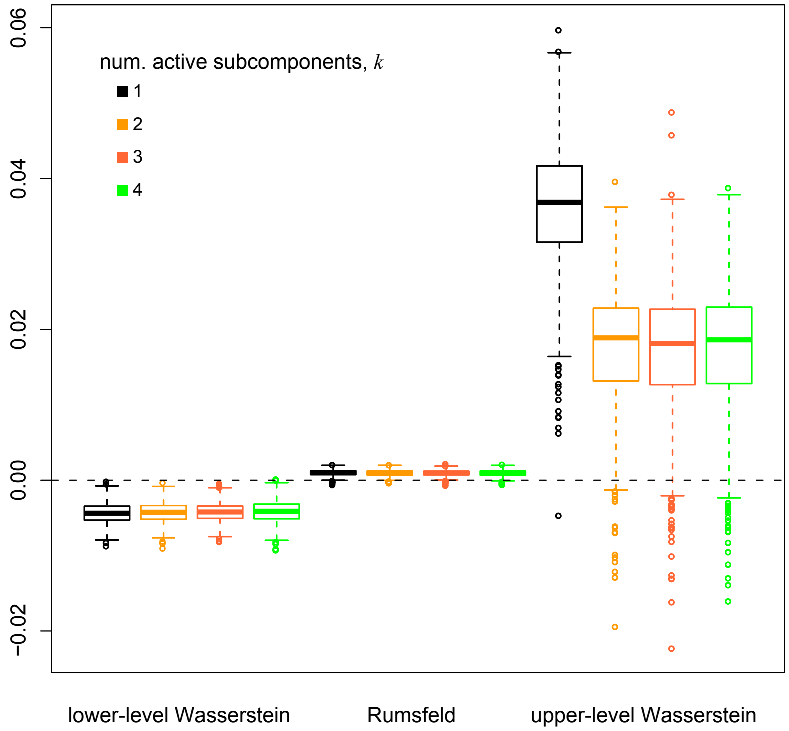


**Figure S6: Selecting among hierarchical mixture models for analyes of the human-lung transcriptomic dataset.** We assessed the fit of the human-lung dataset to our hierarchical Bayesian mixture model with $k = \{1, 2, 3, 4\}$ active subcomponents using posterior-predictive simulation. Here, we depict boxplots for three summary statistics; the lower-level Wasserstein, the upper-level Wasserstein, and the Rumsfeld statistics. These analyses indicate that a model with at least two active subcomponents provides an adequate fit to these data.

*Power.*—We assessed the power of our method to infer the known expression state of genes in the human-lung transcriptome. For a given gene, we record the posterior probability inferred for its known expression state; and then compute the average of this value for all genes. Specifically, we compute:

$$\text{power} = \frac{1}{|A| + |I|} \left[ \sum_{a \in A} P(z_a^a = 1 \mid X) + \sum_{i \in I} P(z_i^a = 0 \mid X) \right],$$

where $|A|$ and $|I|$ are the numbers of known active and inactive genes, respectively.

*Theoretical maximum power.*—We can imagine a threshold-based method for inferring expression states: such a method would specify a precise expression level, and all genes above (below) that expression level would be classified as active (inactive). Recall that the expression state of genes in the human-lung transcriptome are known (from epigenetic marks). Accordingly, for our hypothetical threshold-based method, there is a threshold value that will maximize the number of genes correctly assigned to the known in/active expression state. We performed exactly this experiment: for each library, we used this threshold-based method to classify the expression state of genes for all threshold values between –10 and 10 log TPM (in increments of 0.5 log TPM). For each threshold value, we computed the fraction of active genes correctly classified as active (true-positive rate) and the fraction of inactive genes incorrectly classified as active (false-positive rate). We then inferred the expression state of each gene in the human-lung transcriptomic dataset using our hierarchical Bayesian mixture model. The same threshold procedure was applied to the probability of active expression between 0 and 1

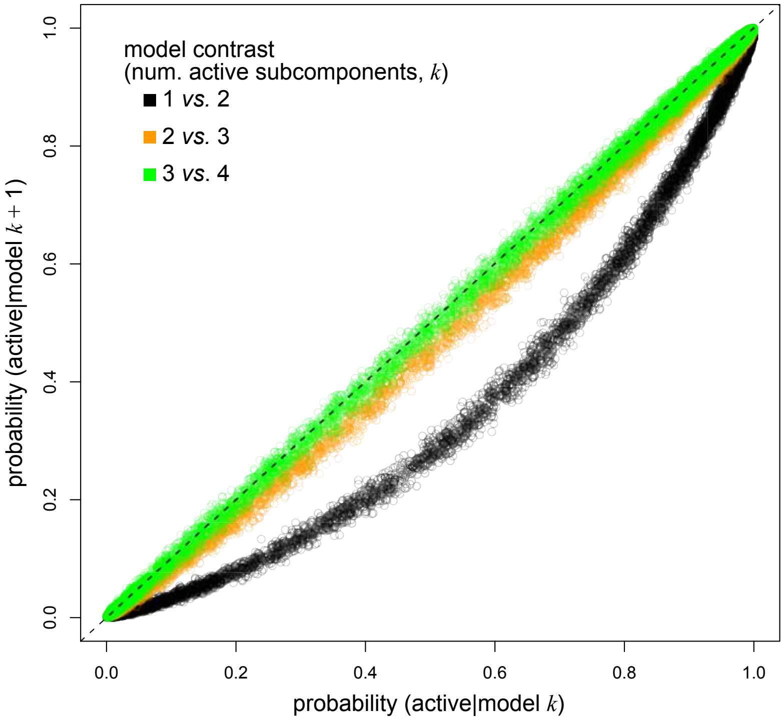


**Figure S7: Sensitivity of expression-state estimates for the human-lung dataset to the assumed number of active subcomponents.** We compared expression-state estimates for genes in the human-lung transcriptome under models with $k = \{1, 2, 3, 4\}$ active subcomponents: the probabilities of assigning genes to the active expression state under a given model and its immediately adjacent, more complex model are plotted on the x- and y-axes, respectively. Specifically, for each gene, $g$, in the the human-lung transcriptome, we plot the probability of active expression, $z_g^a = 1$, inferred under models with 1 *vs.* 2 (black), 2 *vs.* 3 (orange), and 3 *vs.* 4 (green) active subcomponents. Models with $k = \{2, 3, 4\}$ active subcomponents produced nearly identical expression-state estimates for all genes. Conversely, estimates under a model with a single active subcomponent appear to be biased (*i.e.*, causing us to underestimate the uncertainty of the assignment of genes to the in/active expression state).

in increments of 0.01. Finally, we compared the performance of these two methods: the maximum power of the hypothetical threshold-based method—*i.e.*, where an optimal threshold value is selected based on the known expression states—is comparable to that of our hierarchical Bayesian mixture model (which does not rely on the true expression states: Fig. 5A, main text).

##### S2.3.2 Analyses of the *Drosophila*-Testis Transcriptomic Dataset

*Transcriptomic data.*—We dissected tissue samples from $\approx$ 25 mixed-stage adult male fly testes from each of two strains of *D. melanogaster*: Iso-1 and ED10, providing 4 RNA samples. We extracted tissues in phosphate-buffered saline (137 mM NaCl, 10mM Phosphate, 2.7mMKCl, pH 7.4) and then homogenized these tissues in 1 mL Trizol on ice. We added 1 uL LPA (GeneEluteTM; Sigma-Aldrich, Burlington MA) to the homogenate and incubated for 5 min. at room temperature. We then centrifuged for 15 min. at 16,000 Xg at 4 C, extracted the upper-phase supernatant and it transferred to 500 uL isopropanol, which we stored overnight at -20 C. Next, we centrifuged the sample for 30 min. at 16,000 Xg at 4 C, removed the supernatant, added 1 mL 70% EtOH, loosened the pellet, and then spun it down for 10 min. at 15,000 Xg at 4 C. We removed the ethanol and dried the pellet on ice for less than 10 min. We dissolved the dried RNA pellet in DEPC water (30–50 uL), measured the RNA quality using Agilent RNA 6000 Nano kit on BioAnalyzer (Agilent, Waldbronn, Germany), and quantified the amount of RNA using Qubit RNA High Sensitivity Assay kit on Qubit (Life tech-

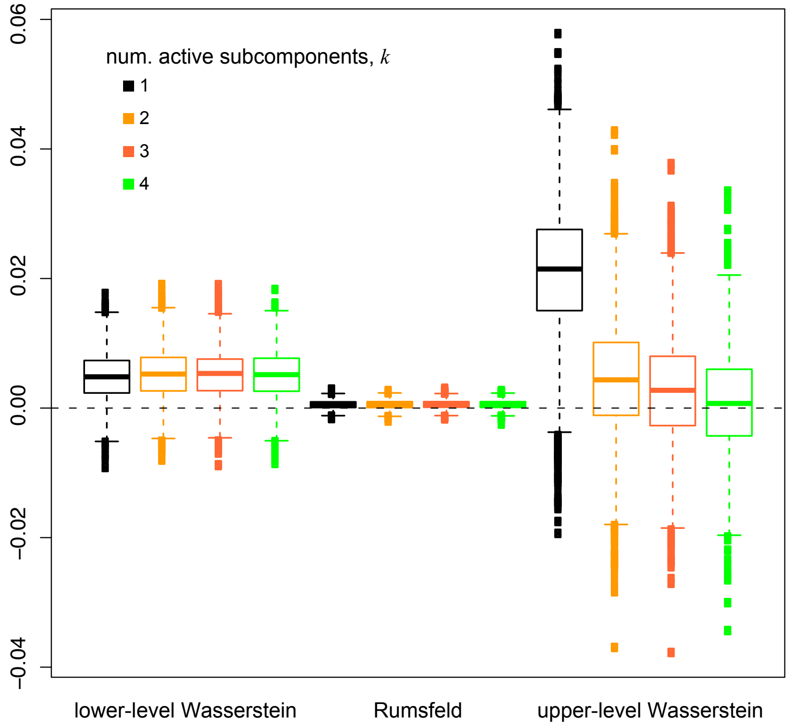


**Figure S8: Selecting among hierarchical mixture models for analyes of the *Drosophila*-testis transcriptomic dataset.** We assessed the fit of the *Drosophila*-testis dataset to our hierarchical Bayesian mixture model with $k = \{1, 2, 3, 4\}$ active subcomponents using posterior-predictive simulation. Here, we depict boxplots for three summary statistics; the lower-level Wasserstein, the upper-level Wasserstein, and the Rumsfeld statistics. These analyses indicate that a model with at least three active subcomponents provides an adequate fit to these data.

nologies) and stored at -20 C. We isolated mRNA from total RNA using NEBNext Poly(A) mRNA magnetic isolation and constructed RNA-seq directional paired-end libraries using NEBNext Ultra Directional RNA Library prep kit. We quantified library size using NEBNext Library Quant Kit (New England BioLabs, Ipswich, MA). Sequencing was performed on the Illumina HiSeq2500 for 1 library of iso-1 strain and Illumina HiSeq3000 and HiSeq4000 for the other 3 libraries. After quality filtering with trimmomatic, we obtained 10, 22, 38, and 54 million reads for each of the 4 libraries. Between 92% and 93% of the reads uniquely mapped to the *D. melanogaster* reference genome annotation r6.12 downloaded from FlyBase.

*Model selection.*—Posterior-predictive simulation indicated that the improvement in model adequacy plateaued for model with three or more active subcomponents (Fig. S8).

*Sensitivity analyses.*—As in our previous findings, our analyses of the *Drosophila*-testis dataset indicate that expression-state estimates under the hierarchical Bayesian mixture model are robust to model overspecification: estimates under models with three or four active subcomponents are virtually identical (Fig. S9). Conversely, expression-state estimates under a model with one or two active subcomponents depart quite strongly from those inferred with three or more active subcomponents, especially for genes with intermediate probabilities of being active.

*Power.*—Our analyses of the *Drosophila*-testis dataset allows us to assess the power of our method to correctly infer expression states of genes that are known to be expressed (or not) in this tissue.

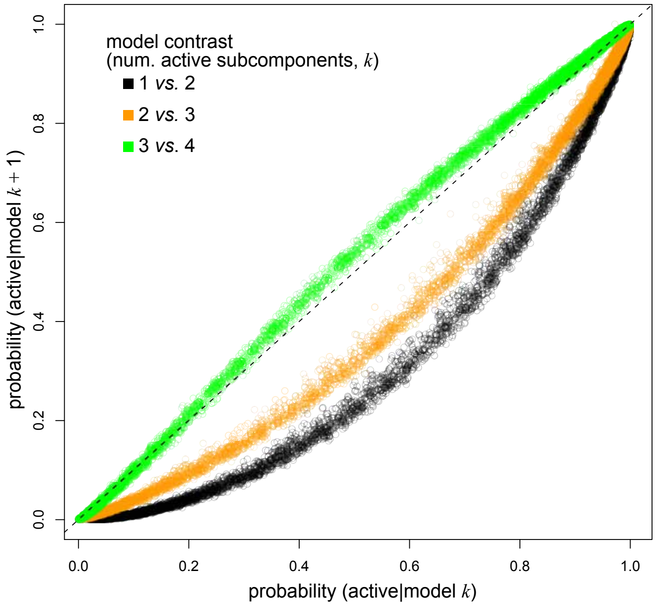


**Figure S9: Sensitivity of expression-state estimates for the *Drosophila*-testis dataset to the assumed number of active subcomponents.** We compared expression-state estimates for genes in the *Drosophila*-testis transcriptome under models with $k = \{1,2,3,4\}$ active subcomponents: the probabilities of assigning genes to the active expression state under a given model and its immediately adjacent, more complex model are plotted on the x- and y-axes, respectively. Specifically, for each gene, $g$, in the the *Drosophila*-testis transcriptome, we plot the probability of active expression, $z_g^a = 1$, inferred under models with 1 *vs.* 2 (black), 2 *vs.* 3 (orange), and 3 *vs.* 4 (green) active subcomponents. Models with $k = \{3,4\}$ active subcomponents produced nearly identical expression-state estimates for all genes. Conversely, estimates under a model with one or two active subcomponents appear to be biased (*i.e.*, causing us to underestimate the uncertainty of the assignment of genes to the in/active expression state).

Specifically, there are 39 genes that are known to be active in the stem-cell niche of the testis based on developmental-genetic evidence (see Table S4 for the list of genes/studies). Additionally, there are odorant-receptor genes (prefix; Or, $n = 59$) and gustatory-receptor genes (prefix; Gr, $n = 60$) that we assume are inactive in the testis; we extracted these genes from the r6.12 annotation. We inferred the expression state of these genes using our hierarchical Bayesian mixture model: the inferred expression states agree closely with their known expression states (Fig. S10), and the power to correctly infer expression states approaches the theoretical maximum for this inference problem (Fig. 4, main text).

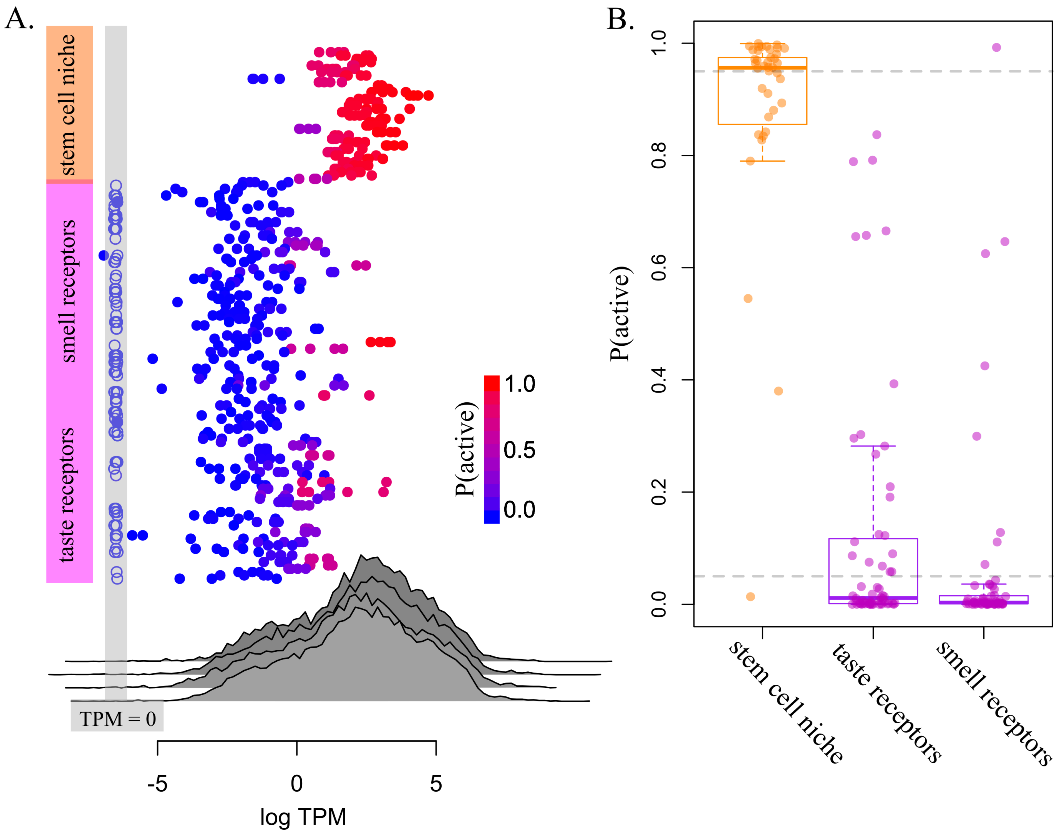


**Figure S10: Assessing the power to correctly infer known expression states of genes in the *Drosophila*-testis dataset.** Panel A. We used our hierarchical Bayesian mixture model to infer the expression state of 39 genes that are known to be expressed in the stem-cell niche of the *Drosophila* testis (see Table S4), and the expression state of 119 odorant- and gustatory-receptor genes that are known to be inactive in the *Drosophila* testis. Here, we depict the observed expression levels of all genes in the four libraries (gray distributions), and the inferred posterior probability that each gene is active (inset legend). Panel B. Boxplots depict the posterior probabilities of active expression for each of the three types of genes.

##### S2.3.3 Analyses of the Primate-Brain Transcriptomic Datasets

We used our method to analyze a published primate-brain transcriptomic dataset [22]. We inferred the expression state of all protein-coding genes in six brain regions—amygdala (AMY), ventral frontal cortex (VFC), dorsal frontal cortex (DFC), superior temporal cortex (STC), striatum (STR), and the area 1 visual cortex (V1C)—sampled from humans, chimpanzees, and rhesus macaques. Our goal is to identify the subset of genes that have a expression state in humans, *i.e.*, where a given gene is inferred to be in/active in humans but not chimpanzees and macaques.

*Transcriptomic data.*—We downloaded the primate orthology table (prNOG) from the EggNOG v. 4.5 database [23], which is based on ensembl release 70 annotations. We downloaded RPKM estimates for six human-brain regions from the Brainspan project (http://www.brainspan.org), as well as published reads [22] for those brain regions in chimp and macaque (from NCBI Bioproject PRJNA236446). We mapped single-end reads to the genome and annotation from ensembl release 70 using `STAR` v. 2.5.3 [24]. We assembled reference-only transcripts using `Stringtie` v. 1.3.4 [25] and estimated relative-expression levels (TPM). We then used `Bedtools` intersect to filter genes that were not present in both the ensembl release 70 and 65 (enforcing at least 98% overlap sequence length).

*Model selection and adequccy.*—We explored the relative and absolute fit of candidate hierarchical Bayesian mixture models with one and two active subcomponents to the three transcriptomic datasets (macaque, chimp and human). Posterior-predictive simulation indicated that a model with two active subcomponents provided an adequate fit to all three datasets (Fig. S12–S13), which we used for all subsequent analyses. We set the prior-mean thresholds for analyses of the the macaque, chimp, and human datasets to $\{\{1\}, \{1,4\}, \{0,3\}\}$, respectively.

*Sensitivity analysis.*—Our analyses of the primate-brain datasets indicate that expression-state estimates under the hierarchical Bayesian mixture model are largely insensitive to the choice of inference model (*i.e.*, the assumed number of active subcomponents: Fig. S14).

*Results.*—We used our method to infer the expression state of all protein-coding genes in six brain regions of the three primate transcriptomic datasets. We then identified the subset of approximately 12,000 1:1:1 orthologous genes in the three species. We then identified the subset of these orthologous genes that have a unique expression state in humans, *i.e.*, where a given gene is inferred to be in/active in humans but not chimpanzees and macaques. Specifically, for each 1:1:1 orthologous gene, $g$, we computed the probability that it was uniquely *active* in humans, $P_{\text{ua}}(z_g^a)$, as:

$$P_{\text{ua}}(z_g^a) = \underbrace{P(z_g^a \mid \mathbf{X}_H)}_{\substack{\text{prob. active} \\ \text{in humans}}} \times \underbrace{\Big(1 - P(z_g^a \mid \mathbf{X}_M)\Big)}_{\substack{\text{prob. inactive} \\ \text{in macaques}}} \times \underbrace{\Big(1 - P(z_g^a \mid \mathbf{X}_C)\Big)}_{\substack{\text{prob. inactive} \\ \text{in chimps}}},$$

where $\mathbf{X}_H, \mathbf{X}_M, \mathbf{X}_C$ refers to the transcriptomic data for humans, macaques, and chimps, respectively. We inferred a gene to be uniquely active in humans if $P_{\text{ua}}(z_g^a) \geq 0.95$. Similarly, we computed the probability that each gene was uniquely *inactive* in humans, $P_{\text{ui}}(z_g^i)$, as:

$$P_{\text{ui}}(z_g^i) = \underbrace{\Big(1 - P(z_g^a \mid \mathbf{X}_H)\Big)}_{\substack{\text{prob. inactive} \\ \text{in humans}}} \times \underbrace{P(z_g^a \mid \mathbf{X}_M)}_{\substack{\text{prob. active} \\ \text{in macaques}}} \times \underbrace{P(z_g^a \mid \mathbf{X}_C)}_{\substack{\text{prob. active} \\ \text{in chimps}}},$$

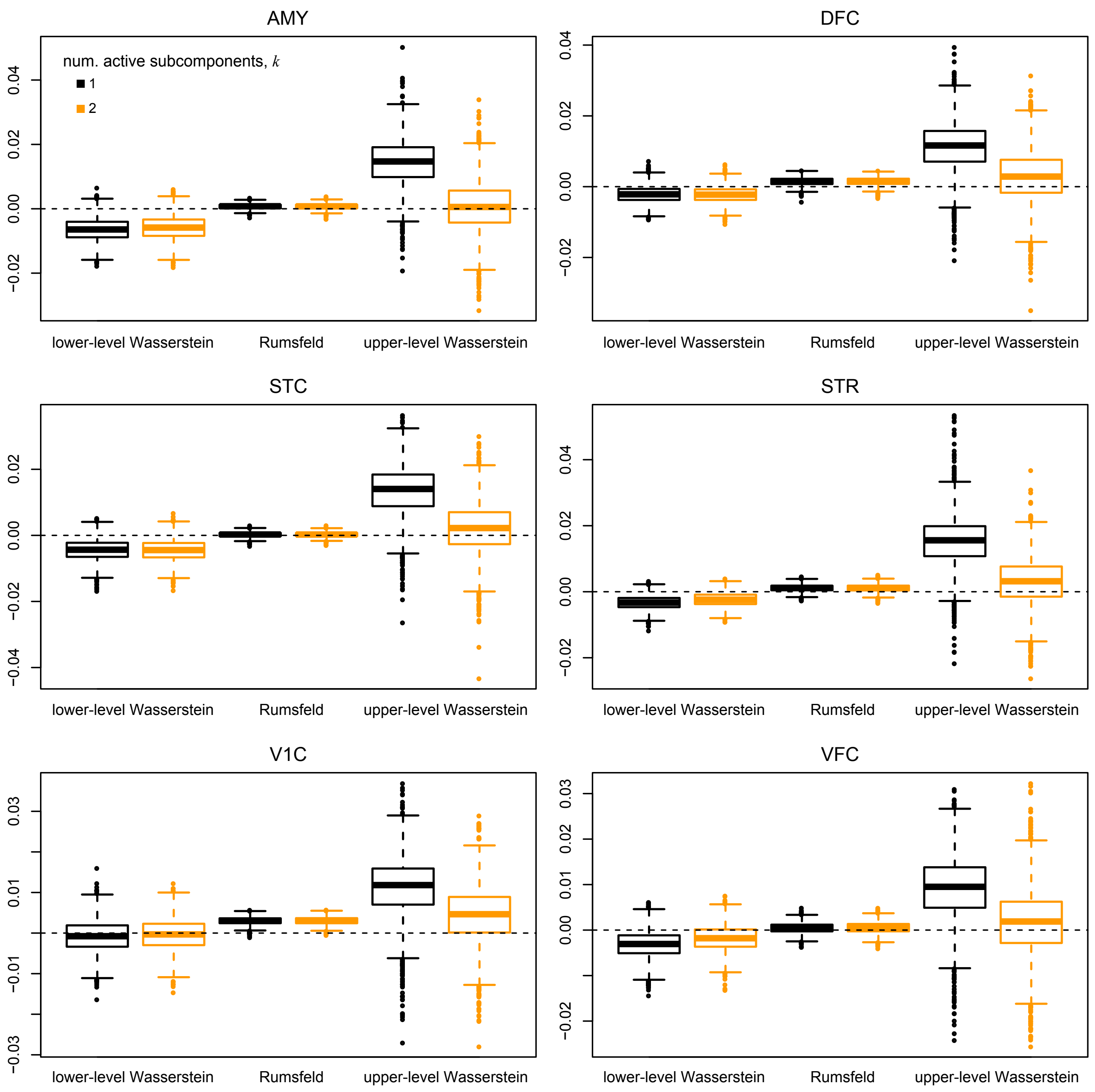


**Figure S11: Selecting among hierarchical mixture models for analyes of the macaque-brain transcriptomic datasets.** We used posterior-predictive simulation to assess the absolute fit of two candidate hierarchial mixture models—with one (black) or two (orange) active subcomponents—to each of the six macaque transcriptomic datasets (one for each of the six brain regions); amygdala (AMY), ventral frontal cortex (VFC), dorsal frontal cortex (DFC), superior temporal cortex (STC), striatum (STR), and the area 1 visual cortex (V1C). For each brain region, we depict boxplots for three summary statistics; the lower-level Wasserstein, the upper-level Wasserstein, and the Rumsfeld statistics. These analyses indicate that a model with two active subcomponents provides an adequate fit to these data.

where we inferred a gene to be uniquely inactive in humans if $P_{\text{ui}}(z^i_g) \geq 0.95$. Across the six brain regions, we identified 9 to 20 genes that were uniquely active in humans and 16 to 23 genes that were uniquely inactive in humans, with the greatest number of unique expression states located in the striatum (Fig. 5, main text). Interestingly, a previous study [22] inferred significant quantitive

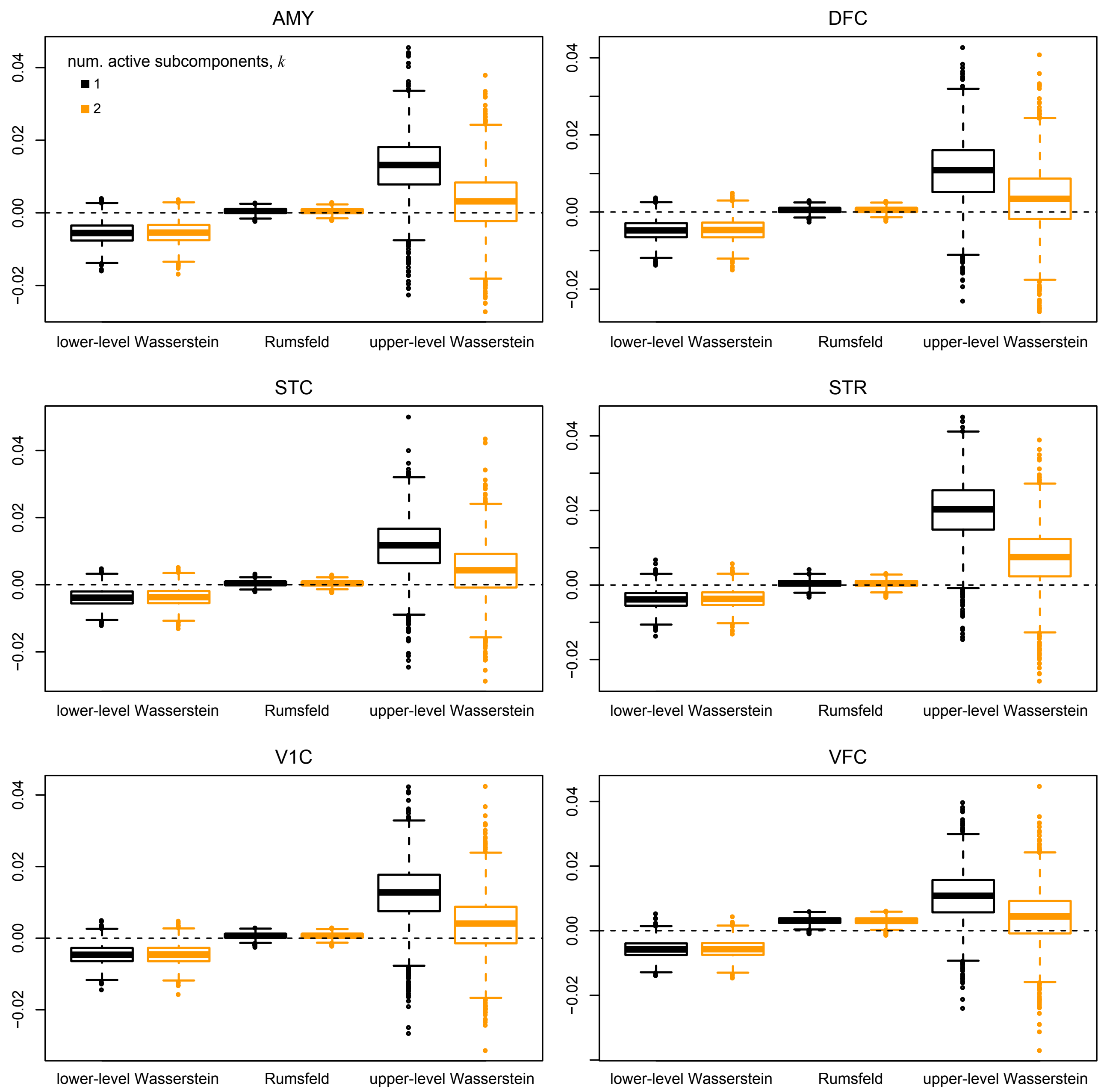


**Figure S12: Selecting among hierarchical mixture models for analyes of the chimpanzee-brain transcriptomic datasets.** We used posterior-predictive simulation to assess the absolute fit of two candidate hierarchial mixture models—with one (black) or two (orange) active subcomponents—to each of the six chimpanzee transcriptomic datasets (one for each of the six brain regions); amygdala (AMY), ventral frontal cortex (VFC), dorsal frontal cortex (DFC), superior temporal cortex (STC), striatum (STR), and the area 1 visual cortex (V1C). For each brain region, we depict boxplots for three summary statistics; the lower-level Wasserstein, the upper-level Wasserstein, and the Rumsfeld statistics. These analyses indicate that a model with two active subcomponents provides an adequate fit to these data.

differences in expression level in $\approx$ 26% of genes in the primate-brain transcriptome; by contrast, analyses using our method reveal that only a tiny fraction of these genes exhibit qualitative difference in expression level.

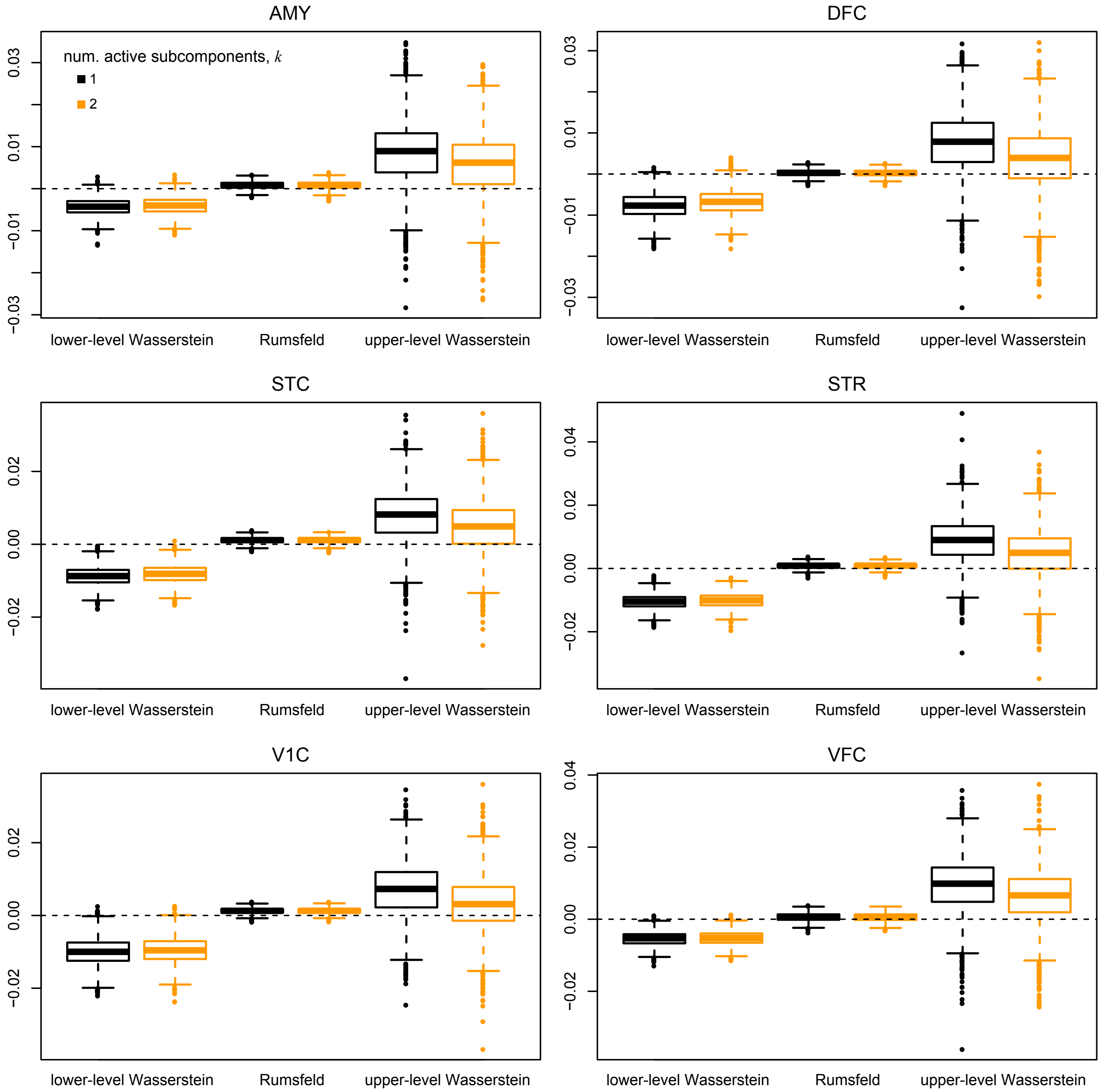


**Figure S13: Selecting among hierarchical mixture models for analyes of the human-brain transcriptomic datasets.** We used posterior-predictive simulation to assess the absolute fit of two candidate hierarchial mixture models—with one (black) or two (orange) active subcomponents—to each of the six human transcriptomic datasets (one for each of the six brain regions); amygdala (AMY), ventral frontal cortex (VFC), dorsal frontal cortex (DFC), superior temporal cortex (STC), striatum (STR), and the area 1 visual cortex (V1C). For each brain region, we depict boxplots for three summary statistics; the lower-level Wasserstein, the upper-level Wasserstein, and the Rumsfeld statistics. These analyses indicate that a model with two active subcomponents provides an adequate fit to these data.

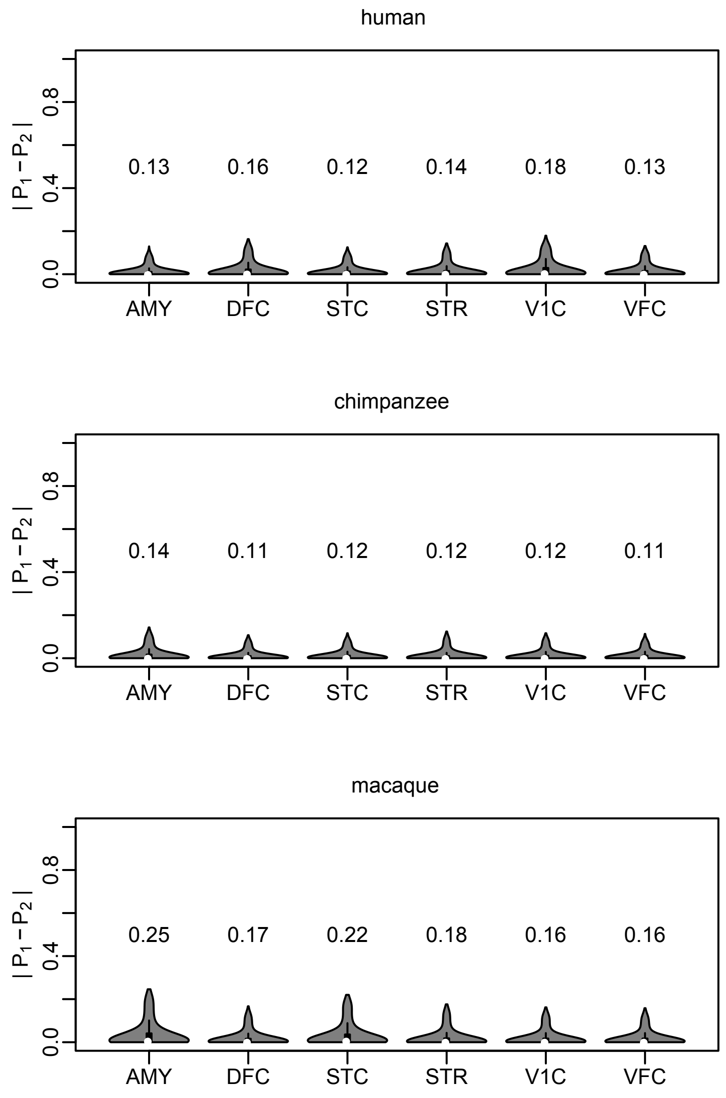


**Figure S14: Sensitivity of expression-state estimates for the primate-brain datasets to the assumed number of active subcomponents.** We compared estimates of the expression state of genes in the primate-brain transcriptomes under models with $k = \{1, 2\}$ active subcomponents. Violin plots depict the absolute difference in expression-state probabilities (*i.e.*, the probability of active expression, $Pz_g^a = 1$) estimated under a model with one ($P_1$) and two active subcomponents ($P_2$). Above each violin plot, we indicate the maximum difference in expression-state probabilities under the two candidate models (which are very similar in all six brain regions). These results indicate that expression-states estimates of genes in the primate-brain transcriptomes are largely insensitive to the choice of model (*i.e.*, the assumed number of active subcomponents.)

**Table S4:** Genes in the *D. melanogaster* stem-cell niche.

| Gene name | FlyBase ID | Reference |
|---|---|---|
| med | FBgn0011655 | [26] |
| put | FBgn0003169 | [26] |
| Dad | FBgn0020493 | [26] |
| dpp | FBgn0000490 | [26] |
| Mad | FBgn0011648 | [26] |
| gbb | FBgn0024234 | [26] |
| Apc1 | FBgn0015589 | [27] |
| Apc2 | FBgn0026598 | [27] |
| Fas3 | FBgn0000636 | [27] |
| lin | FBgn0002552 | [28] |
| bowl | FBgn0004893 | [28] |
| ISWI | FBgn0011604 | [29] |
| Nurf301 | FBgn0000541 | [29] |
| hh | FBgn0004644 | [30] |
| EYA | FBgn0000320 | [30] |
| esg | FBgn0001981 | [31] |
| shg | FBgn0003391 | [32] |
| aop | FBgn0000097 | [33] |
| kek1 | FBgn0015399 | [33] |
| fng | FBgn0011591 | [33] |
| CG2003 | FBgn0039886 | [33] |
| Nrt | FBgn0004108 | [33] |
| ImpL2 | FBgn0001257 | [33] |
| neur | FBgn0002932 | [33] |
| Apt | FBgn0015903 | [33] |
| wnt4 | FBgn0010453 | [33] |
| magu | FBgn0262169 | [33] |
| bnb | FBgn0001090 | [33] |
| zfh1 | FBgn0004606 | [34] |
| STAT92E | FBgn0016917 | [34, 35] |
| chinmo | FBgn0086758 | [34, 36] |
| upd1 | FBgn0004956 | [34] |
| bam | FBgn0000158 | [34, 37] |
| wg | FBgn0004009 | [34] |
| tj | FBgn0000964 | [34] |
| Socs36E | FBgn0041184 | [33, 38] |
| Phf7 | FBgn0031091 | [39] |
| tut | FBgn0052364 | [37] |
| sda | FBgn0015541 | [40] |

#### References


[1] Daniel Hebenstreit and Sarah A. Teichmann. Analysis and simulation of gene expression profiles in pure and mixed cell populations. *Physical Biology*, 8(3):035013, May 2011. ISSN 1478-3975. doi: 10.1088/1478-3975/8/3/035013.

[2] Simon Anders and Wolfgang Huber. Differential expression analysis for sequence count data. *Genome Biology*, 11(10):R106, 2010. ISSN 1474-760X. doi: 10.1186/gb-2010-11-10-r106.

[3] Mark D. Robinson, Davis J. McCarthy, and Gordon K. Smyth. edgeR: A Bioconductor package for differential expression analysis of digital gene expression data. *Bioinformatics*, 26(1):139–140, January 2010. ISSN 1367-4803, 1460-2059. doi: 10.1093/bioinformatics/btp616.

[4] Michael I. Love, Wolfgang Huber, and Simon Anders. Moderated estimation of fold change and dispersion for RNA-seq data with DESeq2. *Genome Biology*, 15(12):550, 2014. ISSN 1474-760X. doi: 10.1186/s13059-014-0550-8.

[5] David C. Hoyle, Magnus Rattray, Ray Jupp, and Andrew Brass. Making sense of microarray data distributions. *Bioinformatics (Oxford, England)*, 18(4):576–584, April 2002. ISSN 1367-4803. doi: 10.1093/bioinformatics/18.4.576.

[6] Chuan Lu and Ross D. King. An investigation into the population abundance distribution of mRNAs, proteins, and metabolites in biological systems. *Bioinformatics (Oxford, England)*, 25(16): 2020–2027, August 2009. ISSN 1367-4811. doi: 10.1093/bioinformatics/btp360.

[7] Nicholas Metropolis, Arianna W. Rosenbluth, Marshall N. Rosenbluth, Augusta H. Teller, and Edward Teller. Equation of State Calculations by Fast Computing Machines. *The Journal of Chemical Physics*, 21(6):1087–1092, June 1953. ISSN 0021-9606, 1089-7690. doi: 10.1063/1.1699114.

[8] W. K. Hastings. Monte Carlo sampling methods using Markov chains and their applications. *Biometrika*, 57(1):97–109, April 1970. ISSN 0006-3444. doi: 10.1093/biomet/57.1.97.

[9] Stuart Geman and Donald Geman. Stochastic Relaxation, Gibbs Distributions, and the Bayesian Restoration of Images. In Martin A. Fischler and Oscar Firschein, editors, *Readings in Computer Vision*, pages 564–584. Morgan Kaufmann, San Francisco (CA), January 1987. ISBN 978-0-08-051581-6. doi: 10.1016/B978-0-08-051581-6.50057-X.

[10] Peter J. Green. Reversible jump Markov chain Monte Carlo computation and Bayesian model determination. *Biometrika*, 82(4):711–732, December 1995. ISSN 0006-3444. doi: 10.1093/biomet/82.4.711.

[11] Ziheng Yang. *Molecular Evolution: A Statistical Approach*. Oxford University Press, 2014. ISBN 978-0-19-960260-5.

[12] Andrew Gelman. Understanding posterior p-values. page 8, 2013.

[13] Andrew Gelman, Xiao-Li Meng, and Hal Stern. Posterior predictive assessment of model fitness via realized discrepancies. *Statistica Sinica*, 6(4):733–760, 1996. ISSN 1017-0405.

[14] Martyn Plummer, Nicky Best, Kate Cowles, and Karen Vines. CODA: Convergence diagnosis and output analysis for MCMC. *R News*, 6:7–11, March 2006. ISSN 1609-3631.

[15] Andrew Gelman and Donald B. Rubin. Inference from iterative simulation using multiple sequences. *Statistical Science*, 7(4):457–472, November 1992. ISSN 0883-4237, 2168-8745. doi: 10.1214/ss/1177011136.

[16] The GTEx Consortium. The Genotype-Tissue Expression (GTEx) pilot analysis: Multitissue gene regulation in humans. *Science*, 348(6235):648–660, May 2015. ISSN 0036-8075, 1095-9203. doi: 10.1126/science.1262110.

[17] Marta Melé, Pedro G. Ferreira, Ferran Reverter, David S. DeLuca, Jean Monlong, Michael Sammeth, Taylor R. Young, Jakob M. Goldmann, Dmitri D. Pervouchine, Timothy J. Sullivan, Rory Johnson, Ayellet V. Segrè, Sarah Djebali, Anastasia Niarchou, The GTEx Consortium, Fred A. Wright, Tuuli Lappalainen, Miquel Calvo, Gad Getz, Emmanouil T. Dermitzakis, Kristin G. Ardlie, and Roderic Guigó. The human transcriptome across tissues and individuals. *Science*, 348(6235):660–665, May 2015. ISSN 0036-8075, 1095-9203. doi: 10.1126/science.aaa0355.

[18] John Lonsdale, Jeffrey Thomas, Mike Salvatore, Rebecca Phillips, Edmund Lo, Saboor Shad, Richard Hasz, Gary Walters, Fernando Garcia, Nancy Young, Barbara Foster, Mike Moser, Ellen Karasik, Bryan Gillard, Kimberley Ramsey, Susan Sullivan, Jason Bridge, Harold Magazine, John Syron, Johnelle Fleming, Laura Siminoff, Heather Traino, Maghboeba Mosavel, Laura Barker, Scott Jewell, Dan Rohrer, Dan Maxim, Dana Filkins, Philip Harbach, Eddie Cortadillo, Bree Berghuis, Lisa Turner, Eric Hudson, Kristin Feenstra, Leslie Sobin, James Robb, Phillip Branton, Greg Korzeniewski, Charles Shive, David Tabor, Liqun Qi, Kevin Groch, Sreenath Nampally, Steve Buia, Angela Zimmerman, Anna Smith, Robin Burges, Karna Robinson, Kim Valentino, Deborah Bradbury, Mark Cosentino, Norma Diaz-Mayoral, Mary Kennedy, Theresa Engel, Penelope Williams, Kenyon Erickson, Kristin Ardlie, Wendy Winckler, Gad Getz, David DeLuca, Daniel MacArthur, Manolis Kellis, Alexander Thomson, Taylor Young, Ellen Gelfand, Molly Donovan, Yan Meng, George Grant, Deborah Mash, Yvonne Marcus, Margaret Basile, Jun Liu, Jun Zhu, Zhidong Tu, Nancy J. Cox, Dan L. Nicolae, Eric R. Gamazon, Hae Kyung Im, Anuar Konkashbaev, Jonathan Pritchard, Matthew Stevens, Timothèe Flutre, Xiaoquan Wen, Emmanouil T. Dermitzakis, Tuuli Lappalainen, Roderic Guigo, Jean Monlong, Michael Sammeth, Daphne Koller, Alexis Battle, Sara Mostafavi, Mark McCarthy, Manual Rivas, Julian Maller, Ivan Rusyn, Andrew Nobel, Fred Wright, Andrey Shabalin, Mike Feolo, Nataliya Sharopova, Anne Sturcke, Justin Paschal, James M. Anderson, Elizabeth L. Wilder, Leslie K. Derr, Eric D. Green, Jeffery P. Struewing, Gary Temple, Simona Volpi, Joy T. Boyer, Elizabeth J. Thomson, Mark S. Guyer, Cathy Ng, Assya Abdallah, Deborah Colantuoni, Thomas R. Insel, Susan E. Koester, A. Roger Little, Patrick K. Bender, Thomas Lehner, Yin Yao, Carolyn C. Compton, Jimmie B. Vaught, Sherilyn Sawyer, Nicole C. Lockhart, Joanne Demchok, and Helen F. Moore. The Genotype-Tissue Expression (GTEx) project. *Nature Genetics*, 45:580–585, May 2013. ISSN 1546-1718. doi: 10.1038/ng.2653.

[19] Helen Parkinson, Ugis Sarkans, Nikolay Kolesnikov, Niran Abeygunawardena, Tony Burdett, Miroslaw Dylag, Ibrahim Emam, Anna Farne, Emma Hastings, Ele Holloway, Natalja Kurbatova, Margus Lukk, James Malone, Roby Mani, Ekaterina Pilicheva, Gabriella Rustici, Anjan Sharma, Eleanor Williams, Tomasz Adamusiak, Marco Brandizi, Nataliya Sklyar, and Alvis Brazma. ArrayExpress update—an archive of microarray and high-throughput sequencing-based functional genomics experiments. *Nucleic Acids Research*, 39(suppl_1):D1002–D1004, January 2011. ISSN 0305-1048. doi: 10.1093/nar/gkq1040.

[20] Linn Fagerberg, Björn M. Hallström, Per Oksvold, Caroline Kampf, Dijana Djureinovic, Jacob Odeberg, Masato Habuka, Simin Tahmasebpoor, Angelika Danielsson, Karolina Edlund,

Anna Asplund, Evelina Sjöstedt, Emma Lundberg, Cristina Al-Khalili Szigyarto, Marie Skogs, Jenny Ottosson Takanen, Holger Berling, Hanna Tegel, Jan Mulder, Peter Nilsson, Jochen M. Schwenk, Cecilia Lindskog, Frida Danielsson, Adil Mardinoglu, Åsa Sivertsson, Kalle von Feilitzen, Mattias Forsberg, Martin Zwahlen, IngMarie Olsson, Sanjay Navani, Mikael Huss, Jens Nielsen, Fredrik Ponten, and Mathias Uhlén. Analysis of the human tissue-specific expression by genome-wide integration of transcriptomics and antibody-based proteomics. *Molecular & Cellular Proteomics*, 13(2):397–406, February 2014. ISSN 1535-9476, 1535-9484. doi: 10.1074/mcp.M113.035600.

[21] Roadmap Epigenomics Consortium, Anshul Kundaje, Wouter Meuleman, Jason Ernst, Misha Bilenky, Angela Yen, Alireza Heravi-Moussavi, Pouya Kheradpour, Zhizhuo Zhang, Jianrong Wang, Michael J. Ziller, Viren Amin, John W. Whitaker, Matthew D. Schultz, Lucas D. Ward, Abhishek Sarkar, Gerald Quon, Richard S. Sandstrom, Matthew L. Eaton, Yi-Chieh Wu, Andreas R. Pfenning, Xinchen Wang, Melina Claussnitzer, Yaping Liu, Cristian Coarfa, R. Alan Harris, Noam Shoresh, Charles B. Epstein, Elizabeta Gjoneska, Danny Leung, Wei Xie, R. David Hawkins, Ryan Lister, Chibo Hong, Philippe Gascard, Andrew J. Mungall, Richard Moore, Eric Chuah, Angela Tam, Theresa K. Canfield, R. Scott Hansen, Rajinder Kaul, Peter J. Sabo, Mukul S. Bansal, Annaick Carles, Jesse R. Dixon, Kai-How Farh, Soheil Feizi, Rosa Karlic, Ah-Ram Kim, Ashwinikumar Kulkarni, Daofeng Li, Rebecca Lowdon, GiNell Elliott, Tim R. Mercer, Shane J. Neph, Vitor Onuchic, Paz Polak, Nisha Rajagopal, Pradipta Ray, Richard C. Sallari, Kyle T. Siebenthall, Nicholas A. Sinnott-Armstrong, Michael Stevens, Robert E. Thurman, Jie Wu, Bo Zhang, Xin Zhou, Arthur E. Beaudet, Laurie A. Boyer, Philip L. De Jager, Peggy J. Farnham, Susan J. Fisher, David Haussler, Steven J. M. Jones, Wei Li, Marco A. Marra, Michael T. McManus, Shamil Sunyaev, James A. Thomson, Thea D. Tlsty, Li-Huei Tsai, Wei Wang, Robert A. Waterland, Michael Q. Zhang, Lisa H. Chadwick, Bradley E. Bernstein, Joseph F. Costello, Joseph R. Ecker, Martin Hirst, Alexander Meissner, Aleksandar Milosavljevic, Bing Ren, John A. Stamatoyannopoulos, Ting Wang, and Manolis Kellis. Integrative analysis of 111 reference human epigenomes. *Nature*, 518 (7539):317–330, February 2015. ISSN 1476-4687. doi: 10.1038/nature14248.

[22] André M. M. Sousa, Ying Zhu, Mary Ann Raghanti, Robert R. Kitchen, Marco Onorati, Andrew T. N. Tebbenkamp, Bernardo Stutz, Kyle A. Meyer, Mingfeng Li, Yuka Imamura Kawasawa, Fuchen Liu, Raquel Garcia Perez, Marta Mele, Tiago Carvalho, Mario Skarica, Forrest O. Gulden, Mihovil Pletikos, Akemi Shibata, Alexa R. Stephenson, Melissa K. Edler, John J. Ely, John D. Elsworth, Tamas L. Horvath, Patrick R. Hof, Thomas M. Hyde, Joel E. Kleinman, Daniel R. Weinberger, Mark Reimers, Richard P. Lifton, Shrikant M. Mane, James P. Noonan, Matthew W. State, Ed S. Lein, James A. Knowles, Tomas Marques-Bonet, Chet C. Sherwood, Mark B. Gerstein, and Nenad Sestan. Molecular and cellular reorganization of neural circuits in the human lineage. *Science*, 358(6366):1027–1032, November 2017. ISSN 0036-8075, 1095-9203. doi: 10.1126/science.aan3456.

[23] Jaime Huerta-Cepas, Damian Szklarczyk, Kristoffer Forslund, Helen Cook, Davide Heller, Mathias C. Walter, Thomas Rattei, Daniel R. Mende, Shinichi Sunagawa, Michael Kuhn, Lars Juhl Jensen, Christian von Mering, and Peer Bork. eggNOG 4.5: A hierarchical orthology framework with improved functional annotations for eukaryotic, prokaryotic and viral sequences. *Nucleic Acids Research*, 44(D1):D286–D293, January 2016. ISSN 0305-1048. doi: 10.1093/nar/gkv1248.

[24] Alexander Dobin, Carrie A. Davis, Felix Schlesinger, Jorg Drenkow, Chris Zaleski, Sonali Jha, Philippe Batut, Mark Chaisson, and Thomas R. Gingeras. STAR: Ultrafast universal RNA-seq

aligner. *Bioinformatics (Oxford, England)*, 29(1):15–21, January 2013. ISSN 1367-4811. doi: 10.1093/bioinformatics/bts635.

[25] Mihaela Pertea, Geo M. Pertea, Corina M. Antonescu, Tsung-Cheng Chang, Joshua T. Mendell, and Steven L. Salzberg. StringTie enables improved reconstruction of a transcriptome from RNA-seq reads. *Nature Biotechnology*, 33(3):290–295, March 2015. ISSN 1546-1696. doi: 10.1038/nbt.3122.

[26] E. Kawase. Gbb/Bmp signaling is essential for maintaining germline stem cells and for repressing bam transcription in the *Drosophila* testis. *Development*, 131(6):1365–1375, February 2004. ISSN 0950-1991, 1477-9129. doi: 10.1242/dev.01025.

[27] Yukiko M. Yamashita, D. Leanne Jones, and Margaret T. Fuller. Orientation of asymmetric stem cell division by the APC tumor suppressor and centrosome. *Science*, 301(5639):1547–1550, September 2003. ISSN 0036-8075, 1095-9203. doi: 10.1126/science.1087795.

[28] Stephen DiNardo, Tishina Okegbe, Lindsey Wingert, Sarah Freilich, and Natalie Terry. Lines and bowl affect the specification of cyst stem cells and niche cells in the *Drosophila* testis. *Development*, 138(9):1687–1696, May 2011. ISSN 0950-1991, 1477-9129. doi: 10.1242/dev.057364.

[29] Christopher M. Cherry and Erika L. Matunis. Epigenetic regulation of stem cell maintenance in the *Drosophila testis* via the nucleosome-remodeling factor NURF. *Cell Stem Cell*, 6(6):557–567, June 2010. ISSN 1934-5909. doi: 10.1016/j.stem.2010.04.018.

[30] Marc Amoyel, Justina Sanny, Michael Burel, and Erika A. Bach. Hedgehog is required for CySC self-renewal but does not contribute to the GSC niche in the *Drosophila* testis. *Development*, 140(1):56–65, January 2013. ISSN 0950-1991, 1477-9129. doi: 10.1242/dev.086413.

[31] Justin Voog, Sharsti L. Sandall, Gary R. Hime, Luís Pedro F. Resende, Mariano Loza-Coll, Aaron Aslanian, John R. Yates, Tony Hunter, Margaret T. Fuller, and D. Leanne Jones. Escargot restricts niche cell to stem cell conversion in the *Drosophila* testis. *Cell Reports*, 7(3):722–734, May 2014. ISSN 2211-1247. doi: 10.1016/j.celrep.2014.04.025.

[32] Justin Voog, Cecilia D'Alterio, and D. Leanne Jones. Multipotent somatic stem cells contribute to the stem cell niche in the *Drosophila* testis. *Nature*, 454(7208):1132–1136, August 2008. ISSN 1476-4687. doi: 10.1038/nature07173.

[33] Natalie A. Terry, Natalia Tulina, Erika Matunis, and Stephen DiNardo. Novel regulators revealed by profiling *Drosophila* testis stem cells within their niche. *Developmental Biology*, 294(1):246–257, June 2006. ISSN 0012-1606. doi: 10.1016/j.ydbio.2006.02.048.

[34] Judith L. Leatherman and Stephen DiNardo. Zfh-1 controls somatic stem cell self-renewal in the *Drosophila* testis and nonautonomously influences germline stem cell self-renewal. *Cell Stem Cell*, 3(1):44–54, July 2008. ISSN 1934-5909. doi: 10.1016/j.stem.2008.05.001.

[35] Natalia Tulina and Erika Matunis. Control of stem cell self-renewal in *Drosophila* spermatogenesis by JAK-STAT signaling. *Science*, 294(5551):2546–2549, December 2001. ISSN 0036-8075, 1095-9203. doi: 10.1126/science.1066700.

[36] Maria Sol Flaherty, Pauline Salis, Cory J. Evans, Laura A. Ekas, Amine Marouf, Jiri Zavadil, Utpal Banerjee, and Erika A. Bach. Chinmo is a functional effector of the JAK/STAT pathway that regulates eye development, tumor formation, and stem cell self-renewal in *Drosophila*. *Developmental Cell*, 18(4):556–568, April 2010. ISSN 1534-5807. doi: 10.1016/j.devcel.2010.02.006.

[37] Di Chen, Chan Wu, Shaowei Zhao, Qing Geng, Yu Gao, Xin Li, Yang Zhang, and Zhaohui Wang. Three RNA binding proteins form a complex to promote differentiation of germline stem cell lineage in *Drosophila*. *PLOS Genetics*, 10(11):e1004797, November 2014. ISSN 1553-7404. doi: 10.1371/journal.pgen.1004797.

[38] Melanie Issigonis, Natalia Tulina, Margaret de Cuevas, Crista Brawley, Laurel Sandler, and Erika Matunis. JAK-STAT signal inhibition regulates competition in the *Drosophila* testis stem cell niche. *Science*, 326(5949):153–156, October 2009. ISSN 0036-8075, 1095-9203. doi: 10.1126/science.1176817.

[39] Shu Yuan Yang, Ellen M. Baxter, and Mark Van Doren. Phf7 controls male sex determination in the *Drosophila* germline. *Developmental Cell*, 22(5):1041–1051, May 2012. ISSN 1534-5807. doi: 10.1016/j.devcel.2012.04.013.

[40] Cindy Lim, Shiv Gandhi, Martin L. Biniossek, Lijuan Feng, Oliver Schilling, Siniša Urban, and Xin Chen. An aminopepetidase in the *Drosophila* testicular niche acts in germline stem cell maintenance and spermatogonial dedifferentiation. *Cell reports*, 13(2):315–325, October 2015. ISSN 2211-1247. doi: 10.1016/j.celrep.2015.09.001.
